# Subtyping of major SARS-CoV-2 variants reveals different transmission dynamics

**DOI:** 10.1101/2022.04.10.486823

**Authors:** Hsin-Chou Yang, Jen-Hung Wang, Chih-Ting Yang, Yin-Chun Lin, Han-Ni Hsieh, Po-Wen Chen, Hsiao-Chi Liao, Chun-houh Chen, James C. Liao

## Abstract

SARS-CoV-2 continues to evolve, causing waves of the pandemic. Up to March 2022, eight million genome sequences have accumulated, which are classified into five major variants of concern. With the growing number of sequenced genomes, analysis of the big dataset has become increasingly challenging. Here we developed systematic approaches for comprehensive subtyping and pattern recognition for transmission dynamics. By analyzing the first two million viral genomes as of July 2021, we found that different subtypes of the same variant exhibited distinct temporal trajectories. For example, some Delta subtypes did not spread rapidly, while others did. We identified sets of characteristic single nucleotide variations (SNVs) that appeared to enhance transmission or decrease efficacy of antibodies for some subtypes of the Delta and Alpha variants. We also identified a set of SNVs that appeared to suppress transmission or increase viral sensitivity to antibodies. These findings are later confirmed in an analysis of six million genomes as of December 2021. For the Omicron variant, the dominant type in the world, we identified the subtypes with enhanced and suppressed transmission in an analysis of seven million genomes as of January 2022 and further confirmed the findings in a later analysis of eight million genomes as of March 2022. While the “enhancer” SNVs exhibited an enriched presence on the spike protein, the “suppressor” SNVs are mainly elsewhere. Disruption of the SNV correlation largely destroyed the enhancer-suppressor phenomena. These results suggest the importance of fine subtyping of variants, and point to potential complex interactions among SNVs.

## Introduction

Relative to the original Wuhan strain, SARS-CoV-2 Variants of Concern (Alpha, Beta, Gamma, Delta, and Omicron) and other known variants (e.g., Eta, Iota, Kappa, Lambda, Epsilon, Zeta, Theta, and Mu) have been identified and caused multiple waves of the pandemic. These variants are reported to confer high transmissibility and possible antibody escape, thus posing challenges to the pandemic control measures. Therefore, tracking variants and predicting their risks are crucially important for pandemic control and the development of pharmacological treatments.

Analysis of the SARS-CoV-2 genome sequences has provided unprecedented opportunities for tracking variants [1–3], characterizing the viral genomes [4–6]; investigating molecular and cellular mechanisms [7–9], understanding the viral origin and evolution [2, 10–16], and scrutinizing many other aspects related to the pandemic. These utilities demonstrate the importance of analyzing the viral genome database [17].

Since the beginning of the pandemic, Global Initiative on Sharing Avian Influenza Data (GISAID) (https://www.gisaid.org/) has provided a data depository for viral genome sequences from confirmed cases. As of July 2021, more than 2 million sequences have accumulated, each containing 30K nucleotides, and as of March 2022, eight million sequences have been reported, which provide an opportunity for fine subtyping of SARS-CoV-2 variants. However, as the size of the dataset grows, analysis of the big data has become increasingly challenging. Likelihood-based subtyping approaches are popular but require more model assumptions and intensive computation compared to a model-free approach.

We develop a systematic approach for using correlated SNV sets (CSSs) with allelic association for dimension reduction of the large collection of genomes. The CSSs allow a computationally tractable ways for viral sub-typing and pattern recognition for transmission dynamics. Using this method, we found that within the commonly identified Alpha (aka B.1.1.7), Delta (aka B.1.617.2), and Omicron (aka B.1.1.529) variants, the temporal trajectories differ significantly among their subtypes. We further identified sets of SNVs that behave as transmission “enhancers”, which are associated with increased temporal trajectory of the Alpha, Delta, and Omicron variants, respectively, and sets of transmission suppressors, which are associated with the “suppression” of the variants with the transmission enhancers. These findings suggest the importance of fine subtyping and possible SNV interactions that be important determinants of viral fitness in the context of public health measures.

## Results

### Using CSSs for dimension reduction

Allelic association of SNVs is a hallmark of rising variants (**Supplementary Text 1**, **Figs. S1 – S4**, and **Table S1**). This characteristic allows us to use allelic association as a way to reduce the dimension of the big data and subtype the variants. We grouped SNVs with pairwise associations R^2^ > 0.5, and used an exponential weighted moving average (EWMA) to detect CSSs, while ignoring SNV sets with occurrence frequency lower than 20 (Refer to the **Materials and Methods** section and **Fig. S5A**). The sensitivity, specificity, and robustness of CSS detection are discussed in Supplementary information (**Fig. S5A**).

The genome of viral strains can then be represented by a combination of SNVs in CSSs with a residual term (**Fig. S6**). Through a three-stage dimension reduction, a 29,409 by 2,119,724 matrix of genome sequence is reduced to a 1,366 by 9,848 matrix of CSS (**Fig. S6**). Note that the definition of CSSs can change depending on the purpose of analysis to include any subset of the genomic database, for example, strains identified in different time span, different countries, or different segments of the genome. Additionally, the thresholds for allelic association can also vary to highlight the features of interest.

We identified a total of 1,057 CSSs, each containing 4 – 33 SNVs with a total of 1,366 signature SNVs. We found that 1,053 of 1,057 CSSs can characterize >99.9% of the dominant strain type (Type VI defined in our previous work [5]), which accounts for 2,000,622 (94.38%) of the strains since March, 2020. The statistics are provided (**Table S2**).

**Fig. 1A** shows that the frequency of strains represented by CSSs increased with time, and CSSs almost completely represent the genome variations after July 2020. **Fig. 1B** shows that the residual SNVs became insignificant. Temporal change of the numbers of CSSs provides information for the dynamic evolutionary processes. We analyzed seven datasets that were collected from Dec 2019 to 7 Apr, 5 May, 26 May, 23 Jun, 15 Dec in 2021 and 12 Jan, 23 Feb in 2022 with a sample size of 1,047K, 1,391K, 1,805K, 2,119K, 6,166K, 7,026K, and 8,475K, respectively, to infer the number of CSSs (**Fig. S7**). As of 23 Jun, 2021 (n = 2,119K), compared to the Alpha strain that the increase in the number of CSSs attenuates, the Delta variant exhibits a steep increase in number of CSSs, indicating a continued evolution of the Delta variant. As of 15 Dec, 2021 (n = 6,166K), compared to the Alpha and Delta strains, the Omicron variant exhibits a much steeper increase in number of CSSs, illustrating a rapid transmission of the Omicron variant. Apart from the Alpha, Delta, and Omicron, other variants have a more limited change and evolutional progress. If we use each CSS to define a subtype, these subtypes can effectively represent the whole population in recent dates (**Supplementary Text 2** and **Fig. S8**). Therefore, CSSs can serve as a basis for both dimension reduction and subtyping, which captures the genome evolution in a computationally tractable manner.

**Figure 1.**
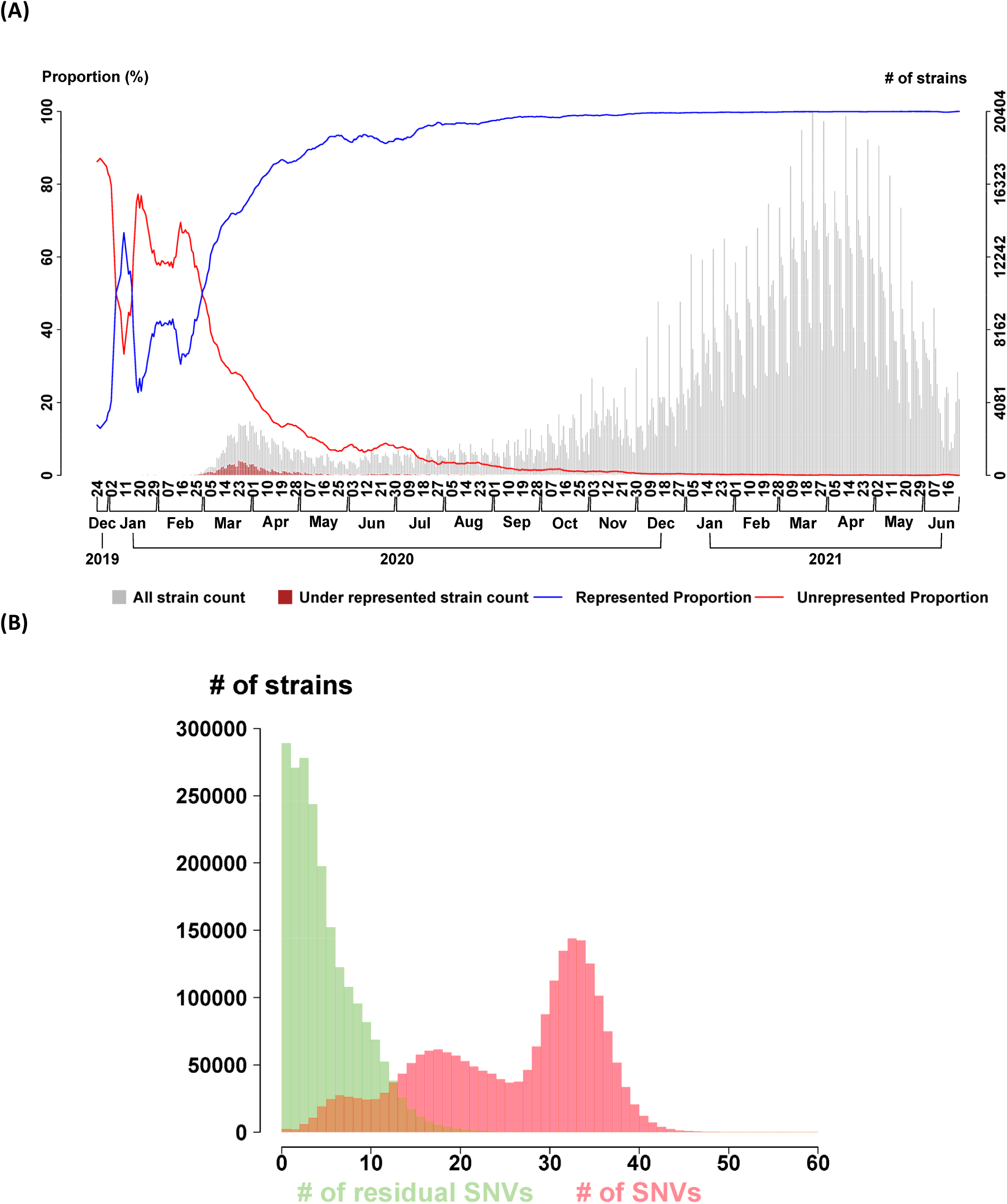
Distributions of the represented strains and SNVs by correlated SNV sets (CSSs) (n = 2,119K genomes as of 23 Jun, 2021). (A) Represented viral strains by CSSs. The temporal proportions of viral strains represented (blue curve) and underrepresented (red curve) by CSSs are displayed. The number of total strains (gray bar) and the number of under-represented stains (red bar) are displayed with the two histograms in the background. **(B) Represented SNVs by CSSs**. Distributions of the number of residual SNVs (i.e., the SNVs under represented by any of CSSs, green bar) and the number of entire SNVs in a strain (i.e., union of SNVs represented and under represented by any of CSSs, red bar) are displayed. The median number of residual SNVs is 4. The median number of entire SNVs is 29.

Importantly, although the CSSs are assigned to the variant with the same Pango nomenclature, they may exhibit different temporal trajectories. All of them carry the core SNVs of the CSSs but some additional SNVs may be different and influence the fitness and transmission of the CSSs. Examples for the Delta variant (aka B.1.617.2), Alpha variant (aka B.1.1.7), and Omicron variant (aka B.1.1.529) are given in **Fig. 2A**, **Fig. 3A**, and **Fig. 4A**, respectively. Detailed composition of the Delta CSSs, Alpha CSSs, and Omicron CSSs are provided in **Fig. 2B**, **Fig. 3B**, and **Fig. 4B**, respectively.

**Figure 2.**
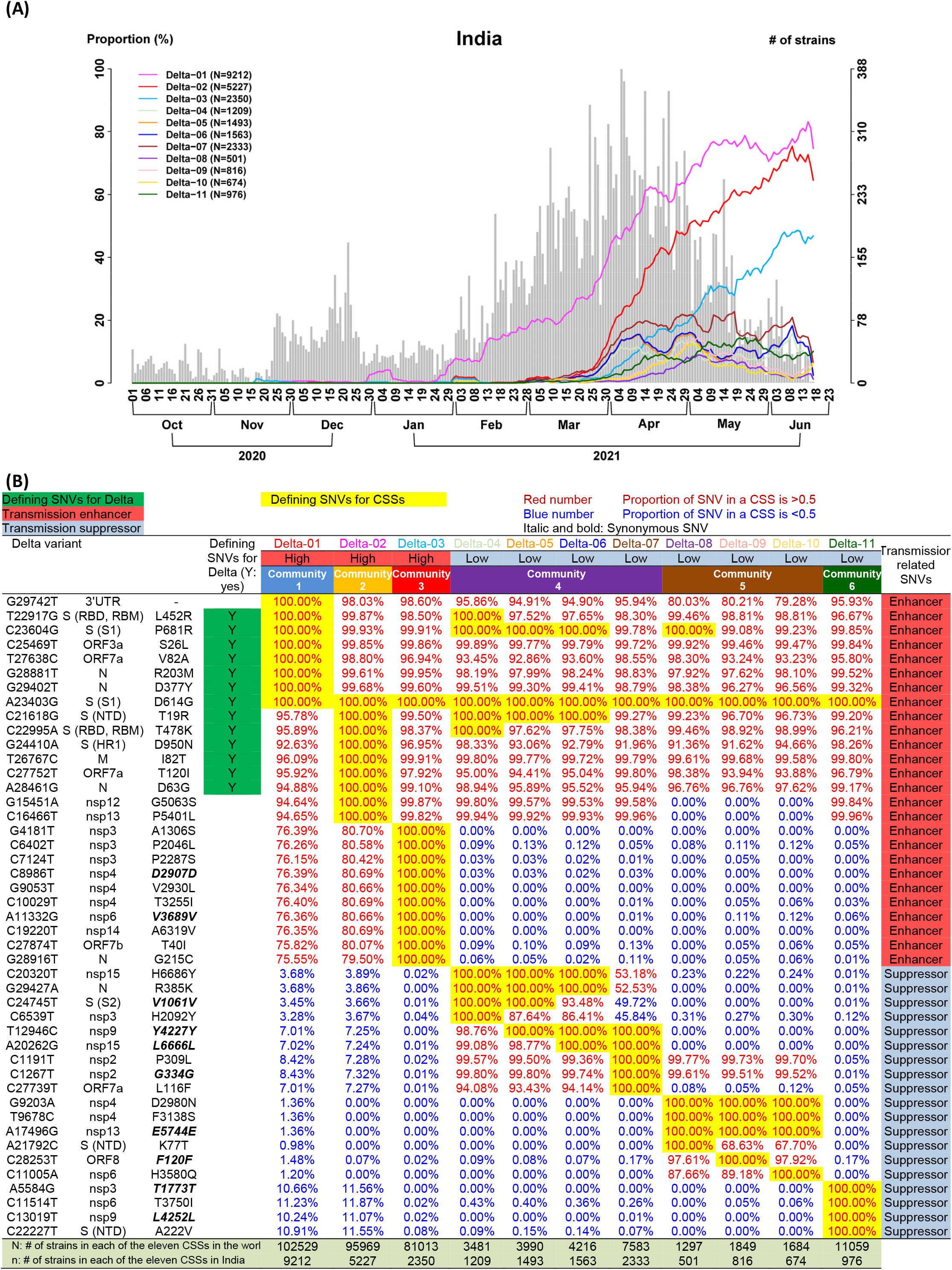
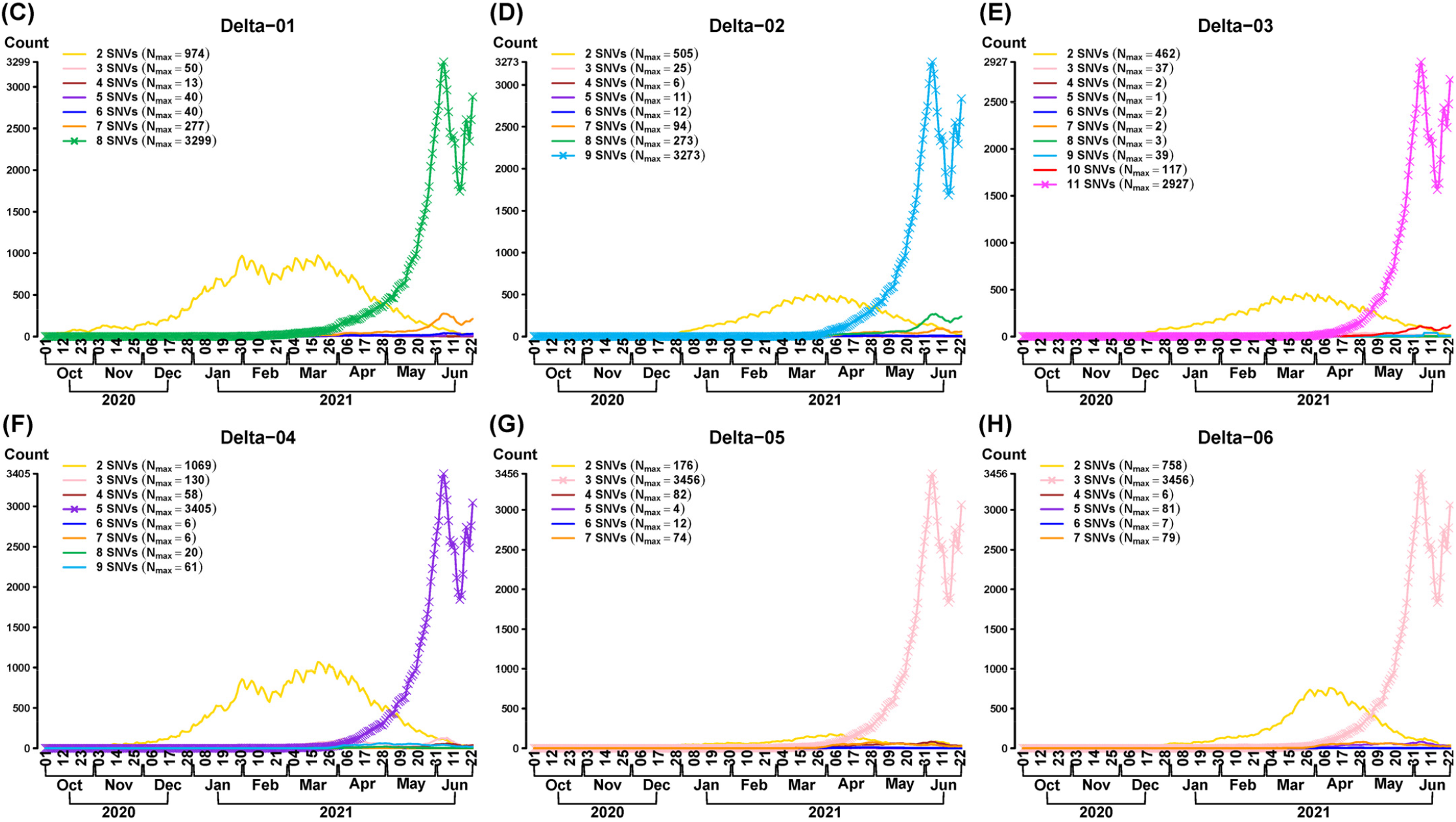
Differential temporal trajectories for the Delta subtypes (n = 2,119K genomes as of 23 Jun, 2021). (A) Eleven Delta (aka B.1.617.2) subtypes (i.e., correlated SNV sets; CSSs) in India. Three CSSs (Delta-01 ~ Delta-03) have an increasing temporal trajectory and eight CSSs (Delta-04 ~ Delta-11) have a much lower temporal trajectory, revealing that CSSs provide a more detailed information for the subtypes and their transmission patterns. **(B) Transmission enhancer and suppressor SNVs.** Eleven CSSs are separated into two categories: the first three CSSs (Delta-01 ~ Delta-03) have a high temporal trajectory and the remaining eight CSSs have a low temporal trajectory. For each of the eleven CSSs, nucleotide change, the protein that a SNV is located, and amino acid change for the signature SNVs are listed and followed by the proportions of the signature SNVs in each of the eleven CSSs. If the proportion is greater than 80%, then the numbers are marked by red color, otherwise, the numbers are marked by blue color. The signature SNVs for each of the eleven CSSs are marked by yellow color. The number of strains in each of the eleven CSSs in the world (N) and in the original country that the Delta variant was discovered (n) are listed. (**C – E**) **Missing SNVs in the three CSSs carrying the transmission enhancer SNVs.** For the first three CSSs, Delta-01 carries 8 transmission enhancer SNVs, Delta-02 carries 9 transmission enhancer SNVs, and Delta-03 carries 11 transmission enhancer SNVs. The highest curve (the curve with a symbol x) indicates the temporal trajectory of this CSS. When some of the transmission enhancer SNVs are missing (the curve without a symbol x), the temporal trajectories are dramatically reduced. This phenomenon explains the transmission enhancer effect. Nmax indicates the maximum number of strains in the temporal trajectory. (**F – H**) **Missing SNVs in the first three CSSs carrying the transmission suppressor SNVs.** On the other hand, for Delta-04, Delta-05, and Delta-06, their CSS signature SNVs contain 5, 3, and 3 transmission enhancer SNVs and 4, 4, and 4 transmission suppressor SNVs, respectively. The highest curve (the curve with a symbol x) indicates the temporal trajectory for a set of strains (not the identified CSSs) that they carry 5, 3, and 3 transmission enhancer SNVs but miss 4, 4, and 4 transmission suppressor SNVs from the three CSSs (Delta-04, Delta-05, and Delta-06), respectively. After 4, 4, and 4 transmission suppressor SNVs are added to from the three CSSs with a total of 9, 7, and 7 defining signature SNVs, respectively, the temporal trajectories of the three CSSs are dramatically reduced. This phenomenon explains the transmission suppressor effect.

**Figure 3.**
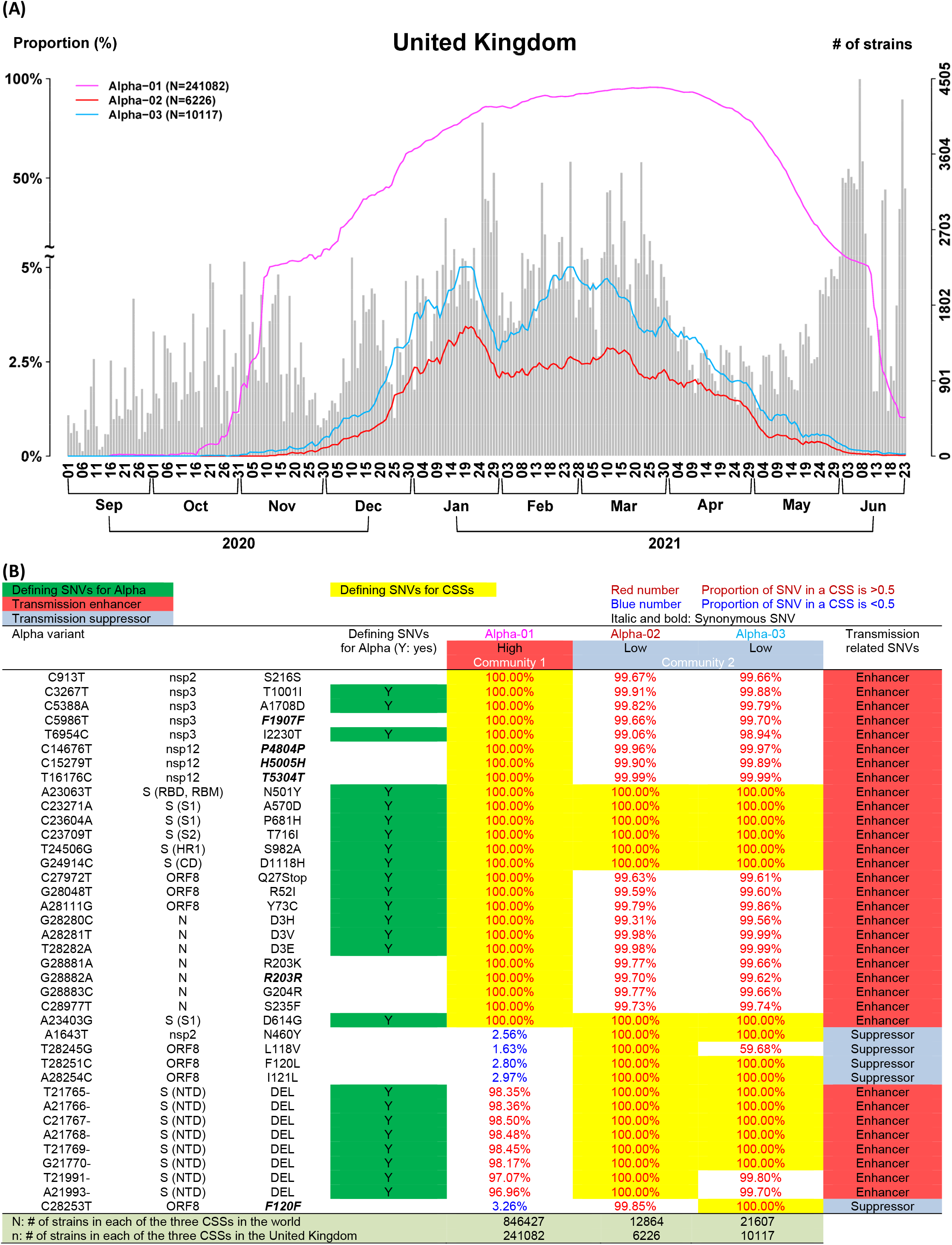
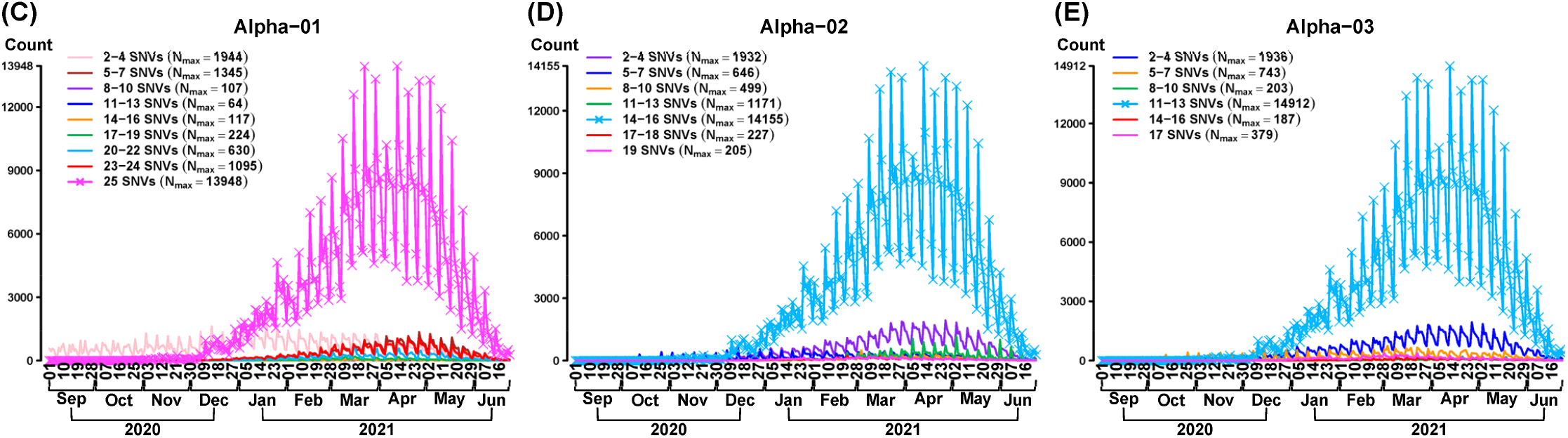
Differential temporal trajectories for the Alpha subtypes (n = 1,047K genomes as of 7 Apr, 2021). (A) Three Alpha (aka B.1.1.7) subtypes (i.e., correlated SNV sets; CSSs) in the United Kingdom. One CSS (Alpha-01) has an increasing temporal trajectory and two CSSs (Alpha-02 and Alpha-03) have a suppressed temporal trajectory, revealing that CSSs provide a more detailed information for the subtypes and their transmission patterns. **(B) Transmission enhancer and suppressor SNVs.** Three CSSs are separated into two categories: the first CSS has a high temporal trajectory and the remaining two CSSs have a low temporal trajectory. For each of the three CSSs, nucleotide change, the protein that a SNV is located, and amino acid change for the signature SNVs are listed and followed by the proportions of the signature SNVs in each of the three CSSs. If the proportion is greater than 80%, then the numbers are marked by red color, otherwise, the numbers are marked by blue color. The signature SNVs for each of the three CSSs are marked by yellow color. The number of strains in each of the three CSSs in the world (N) and in the original country that the Alpha variant was discovered (n) are listed. **(C) Missing SNVs in the three CSSs carrying the transmission enhancer SNVs**. For the first CSS, Alpha-01 carries 25 transmission enhancer SNVs. The highest curve (the curve with a symbol x) indicates the temporal trajectory of this CSS. When some of the transmission enhancer SNVs are missing (the curve without a symbol x), the temporal trajectories are dramatically reduced. This phenomenon explains a transmission enhancer effect. **(D – E) Missing SNVs in the three CSSs carrying the transmission suppressor SNVs**. On the other hand, for the reminding two CSSs (Alpha-02 and Alpha-03), their CSS signature SNVs contain 15 and 13 transmission enhancer SNVs and 4 and 4 transmission suppressor SNVs, respectively. The highest curve (the curve with a symbol x) indicates the temporal trajectory for sets of strains (not the identified CSSs) that they carry 15 and 13 transmission enhancer SNVs but miss 4 and 4 transmission suppressor SNVs in from the two CSSs (Alpha-02 and Alpha-03), respectively. After 4 and 4 transmission suppressor SNVs are added to form the two CSSs with a total of 19 and 17 defining signature SNVs, respectively, the temporal trajectories of the two CSSs are dramatically reduced. This phenomenon explains a transmission suppressor effect.

### The Delta transmission enhancer and suppressor SNVs

To characterize and subtype strain variations in more detail, the CSS approach can be applied to individual countries. As the highly contagious variant Delta was first discovered in India, we further subtyped the Delta strain sequenced in India using the CSS approach, which resulted in eleven subtypes (**Figs. 2A** and **2B**). For illustration, we focus on the first six subtypes. The first three subtypes (Delta-01, Delta-02, and Delta-03) exhibited increasing temporal trajectories and the other three subtypes (Delta-04, Delta-05, and Delta-06) had much lower temporal trajectories (**Fig. 2A**). Remarkably, the same pattern of the differential temporal trajectories for the subtypes was found in many other countries consistently (**Fig. S9**), indicating that the subtypes and their differences are reproducible.

The strains in the six Delta subtypes carry all or the majority of the signature SNVs defined for the Delta variant (T19R, T478K, D950N, D614G, L452R, and P681R in the spike protein, 12 SNVs in other proteins, and 1 in the 3’ untranslated region) (**Fig. 2B**). Yet, the subtypes exhibited distinct temporal trajectories.

The first three subtypes with rising temporal trajectories (Delta-01 to 03 in **Figs. 2A** and **2B**) are defined by eight, nine, and eleven signature SNVs, respectively, with 100% allelic associations. We define these signature SNVs as “transmission enhancers”, since they are strongly associated with the rapid rise in proportion. The remaining three subtypes with lower temporal trajectories (Delta-04 to 06 in **Figs. 2A** and **2B**) all contain a set of 100% associated signature SNVs, in addition to the enhancer SNVs in some strains. It appears that these signature SNVs “suppressed” the rise of the temporal trajectories. Thus, we define them as “transmission suppressors”.

As the enhancer SNVs are 100% associated in Delta-01 to 03, we looked for similar strains without the complete set of the enhancer SNVs. The result shows that strains missing any one of the 8 enhancer SNVs in Delta-01 (**Fig. 2C**), missing any of the 9 enhancer SNVs in Delta-02 (**Fig. 2D**), or missing any of the 11 enhancer SNVs in Delta-03 (**Fig. 2E**) dramatically reduced the temporal trajectory. This phenomenon suggests that the enhancer SNVs work cooperatively to gain viral fitness (i.e., a synergistic or positive cooperativity effect), and disfavors the possible hitchhiking passenger roles played by some SNVs.

In contrast to the first three subtypes, the other three subtypes (Delta-04 to 06) exhibit different temporal patterns (**Fig. 2A**). The majority of the strains in these three subtypes also carried the signature SNVs in the spike protein for the Delta variant (i.e., the T19R-L452R-T478K-D614G-P681R-D950N haplotype) (**Fig. 2B**). Nevertheless, the temporal trajectories of the three subtypes are suppressed dramatically after acquiring the suppressor SNVs located mainly on the non-spike proteins (**Figs. 2F – 2H**).

Delta-04 carries the transmission suppressors C20320T (nsp15), G29427A (N), C24745T (S), C6539T (nsp3), and exhibited the low temporal trajectory (**Figs. 2A - 2B**). Delta-05 and Delta-06 carried overlapping but different sets of transmission suppressors with some synonymous SNVs (**Fig. 2B**). These two subtypes also generally suppressed the rise of the temporal proportion, although not as effective as Delta-04 (**Fig. 2A**).

Note that the defining SNVs for Delta-04 to 06 subtypes (CSSs) contain both suppressors and enhancers. We examined the effect of acquiring defining SNVs, starting from strains containing only the enhancers but not the suppressors (**Figs. 2F - 2H)**. Strains containing the 5, 3, and 3 enhancers in the defining SNV sets of Delta-04 to Delta-06, respectively, but not the suppressors all exhibited a rising temporal pattern. However, when they acquired all the suppressor SNVs, the rising patterns were suppressed. We did not find sufficient samples or presence of partial set of suppressors. Therefore, whether the transmission suppressor act independently or cooperatively to reduce the temporal trajectories remains inconclusive. However, since we did not find partial suppression of the temporal patterns, it is likely that the suppressors also work cooperatively (see a discussion in the subsection *“Suppression effect”).* These transmission suppressor SNVs play a suppressing role on the viral transmission through genetic epistasis with other signature SNVs. Since some of the suppressor SNVs in Delta-04 to 06 are synonymous SNVs, the effect may come from codon usage or RNA-level interactions, although the hitchhiking effect cannot be ruled out.

Remarkably, the identified enhancer and suppressor SNVs can be further confirmed in an analysis of six million SARS-CoV-2 genomes (**Fig. S10**). This reflects the robustness of the identified enhancer and suppressor SNVs.

### The Alpha transmission enhancer and suppressor SNVs

We also applied the CSS-based approach to subtype the Alpha variants identified in UK (**Fig. 3A**), the country where Alpha was first discovered. The first subtype (Alpha-01) presents the much higher temporal trajectory than the other two subtypes (Alpha-02 and Alpha-03). These Alpha subtypes were also found in many other countries, and showed the similar temporal trajectory pattern (**Fig. S11**). We defined the signature SNVs of Alpha-01 as transmission enhancers, as they enhanced the transmission of the variant (**Fig. 3B**). Alpha-02 or 03 contained some of these enhancers, but most importantly, they acquired a set of suppressors that appeared to suppress the rise of the temporal trajectory (**Fig. 3B**).

The proportion of the strains pertaining to the first subtype was reduced dramatically if any of the transmission enhancers were missing (**Fig. 3C**). This finding suggests that the enhancers work cooperatively. Similar to the Delta subtypes, we did not find sufficient samples or presence of partial set of suppressors (**Figs. 3D – 3E**). Therefore, whether the transmission suppressor act independently or cooperatively to reduce the temporal trajectories remains inconclusive.

### The Omicron transmission enhancer and suppressor SNVs

As of 12 Jan, 2022 (n = 7,026K), we applied the CSS-based approach to subtype the Omicron variants in the United Kingdom with a larger sample size compared to other countries (**Fig. 4A**). Omicron is known as the variant with a large number of signature SNVs especially in the spike protein. The first and second subtypes (Omicron-01 and 02) present the much higher temporal trajectory than the third subtype (Omicron-03). These Omicron subtypes were also found in many other countries, and showed the consistent temporal trajectory patterns (**Fig. S12**). We defined the signature SNVs of Omicron-01 and 02 as transmission enhancers, as they enhanced the transmission of the variant (**Fig. 4B**). Omicron-03 contained some of these enhancers, but most importantly, they acquired a set of suppressors that appeared to suppress the rise of the temporal trajectory (**Fig. 4B**). Furthermore, these findings were successfully confirmed in an analysis of eight million SARS-CoV-2 genomes accumulated as of 23 Feb, 2022 (n = 8,475K) (**Fig. S13**).

**Figure 4.**
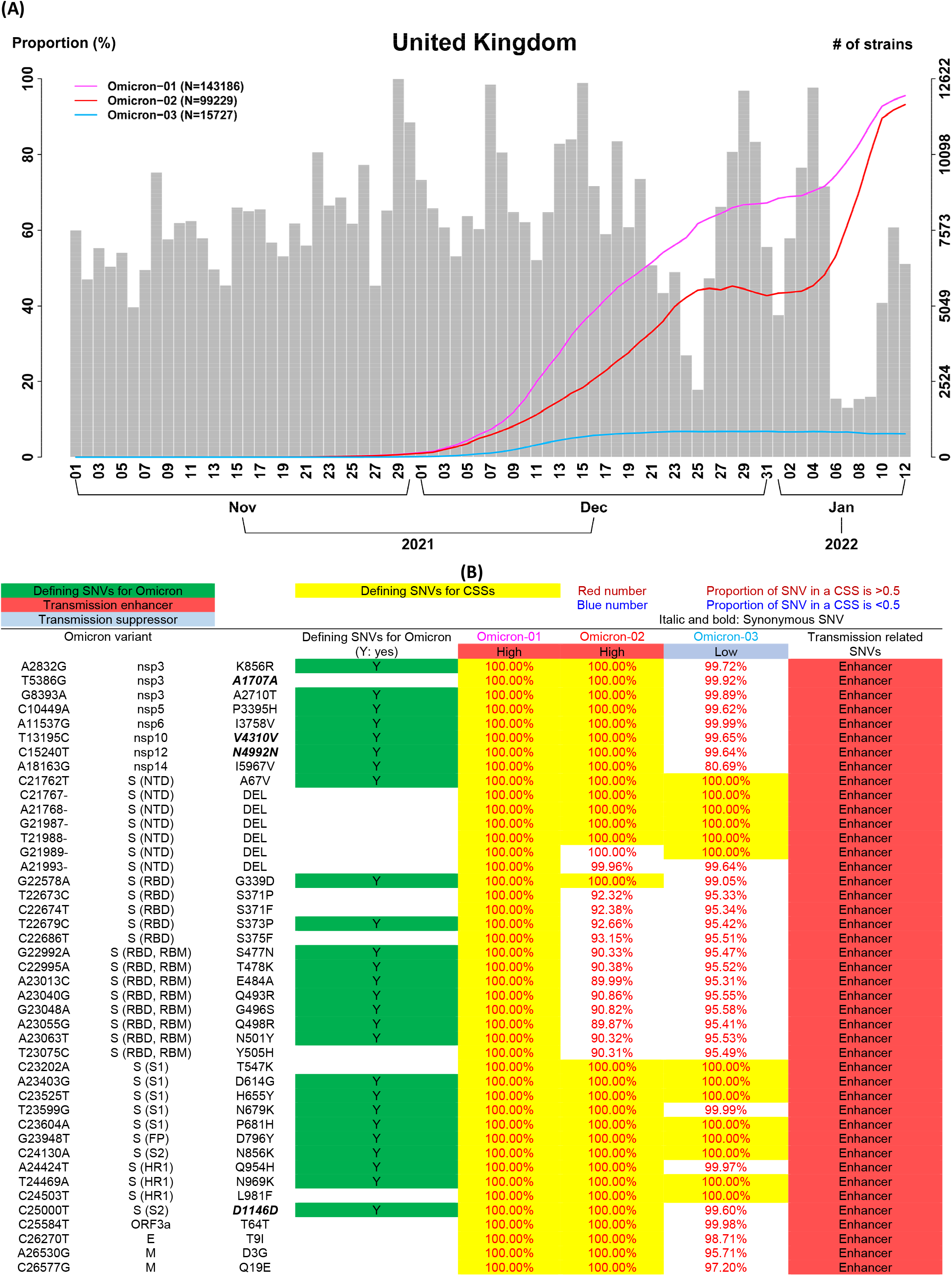
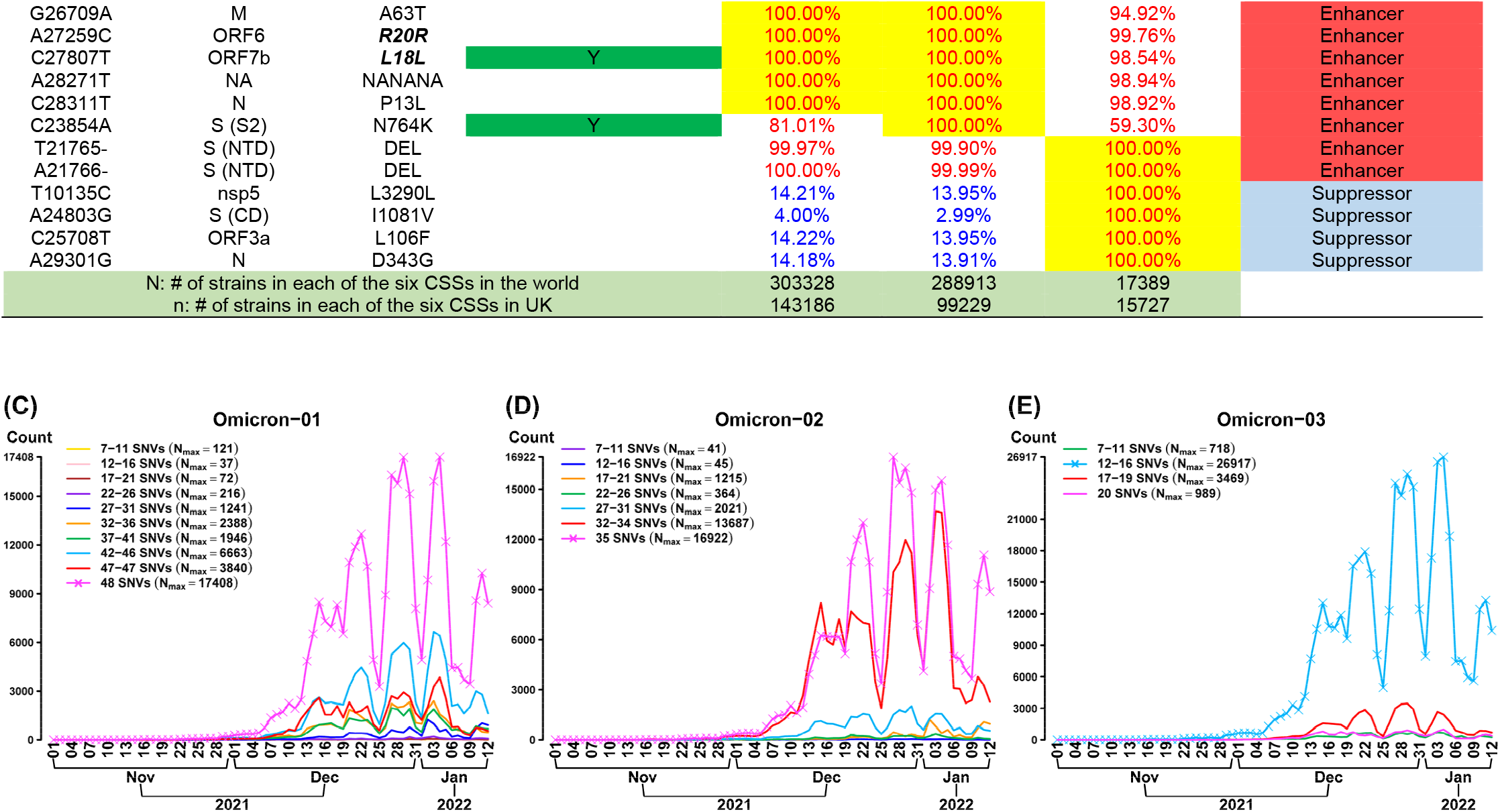
Differential temporal trajectories for the Omicron subtypes (n = 7,026K genomes as of 12 Jan, 2022). (A) Three Omicron (aka B.1.1.529) subtypes (i.e., correlated SNV sets; CSSs) in the United Kingdom. Two CSSs (Omicron-01 and Omicron-02) have an increasing temporal trajectory and one CSS (Omicron-03) have a much lower temporal trajectory, revealing that CSSs provide a more detailed information for the subtypes and their transmission patterns. **(B) Transmission enhancer and suppressor SNVs.** Three CSSs are separated into two categories: the first two CSSs (Omicron-01 and Omicron-02) have a high temporal trajectory and the remaining one CSS has a low temporal trajectory. For each of the three CSSs, nucleotide change, the protein that a SNV is located, and amino acid change for the signature SNVs are listed and followed by the proportions of the signature SNVs in each of the three CSSs. If the proportion is greater than 80%, then the numbers are marked by red color, otherwise, the numbers are marked by blue color. The signature SNVs for each of the three CSSs are marked by yellow color. The number of strains in each of the three CSSs in the world (N) and in the United Kingdom (n) are listed. (**C – D**) **Missing SNVs in the two CSSs carrying the transmission enhancer SNVs.** For the first two CSSs, Omicron-01 carries 48 transmission enhancer SNVs and Omicron-02 carries 35 transmission enhancer SNVs. The highest curve (the curve with a symbol x) indicates the temporal trajectory of this CSS. When some of the transmission enhancer SNVs are missing (the curve without a symbol x), the temporal trajectories are dramatically reduced. This phenomenon explains a transmission enhancer effect. N_max_ indicates the maximum number of strains in the temporal trajectory. (**E**) **Missing SNVs in the CSS carrying the transmission suppressor SNVs.** On the other hand, for Omicron-03, the CSS signature SNVs contain 16 transmission enhancer SNVs and 4 transmission suppressor SNVs. The highest curve (the curve with a symbol x) indicates the temporal trajectory for a set of strains (not the identified CSS) that they carry 16 transmission enhancer SNVs but miss 4 transmission suppressor SNVs from the CSS (Omicron-03). After 4 transmission suppressor SNVs are added to form the CSS with a total of 20 defining signature SNVs, the temporal trajectory of the CSS is dramatically reduced. This phenomenon explains a transmission suppressor effect.

The proportion of the strains pertaining to the first two subtypes were reduced dramatically if any of the transmission enhancers were missing (**Fig. 4C**). This finding suggests that the enhancers work cooperatively. Similar to the Delta and Alpha subtypes, we did not find sufficient samples or presence of partial set of suppressors (**Figs. 4D – 4E**). Therefore, whether the transmission suppressor act independently or cooperatively to reduce the temporal trajectories remains inconclusive. However, the current observation reveals that the four suppressor SNVs appeared together since the first emergence in November 2021. The set of four suppressor SNVs has a stronger effect on transmission suppression compared to the subsets that they miss any suppressor SNVs (see a discussion in the subsection *“Suppression effect”).*

A further analysis of a larger sample size (n = 8,475K) as of 23 Feb, 2022 identified seven Omicron CSSs (**Figs. S14A** and **S14B**), named as Omicron-04 ~ Omicron-10. The first six Omicron subtypes Omicron-04 ~ Omicron-9 exhibited a pattern of enhanced transmission and the final subtype Omicron-10 suppressed transmission. Patterns of enhanced and suppressed transmission in these CSSs were also confirmed in many countries (**Figs. S14C – S14V**). Compared to the previous results (n = 7,026K) in **Fig. 4**, Omicron-04, Omicron-05, and Omicron-10 are the analogue of Omicron-02, Omicron-01, and Omicron-03, respectively. Omicron-06 ~ Omicron-09 are the subtypes additionally identified in the new data, where Omicron-09 is enriched by the Omicron sub-lineage BA.2 strains.

### Suppression effect

Transmission suppression can be contributed by a single spike SNV or a set of suppressor SNVs. Delta-11 (**Fig. 2**) and Omicron-03 (**Fig. 4**) are illustrated as examples. Delta with A222V(S) has a higher temporal trajectory than Delta-11 which carries a full set of transmission suppressor SNVs: T1773T(nsp3)–T3750I(nsp6)–L4252L(nsp9)–A222V(S), indicating that the set of four transmission suppressor SNVs work cooperatively and have a stronger synergistic effect in suppressing transmission suppression than a single SNV A222V(S) (**Fig. 5**). The result also explains the necessity of a CSS analysis compared to a SNV by SNV analysis that it fails to account for genetic epistasis. Both Omicron-03 and Omicron with I1081V(S) have a lower temporal trajectory compared to the one in Omicron with L3290L(nsp5)–L106F(ORF3a)–D343G(N), indicating that I1081V(S) and/or L3290L(nsp5)–I1081V(S)–L106F(ORF3a)–D343G(N) have an effect in suppressing a viral transmission of Omicron (**Fig. 6**). Because of a high allelic association in the transmission suppressor SNVs L3290L(nsp5)–I1081V(S)–L106F(ORF3a)–D343G(N), more data are needed to distinguish that the suppressing effect is contributed by I1081V solely and/or the full set of transmission suppressor SNVs.

**Figure 5.**
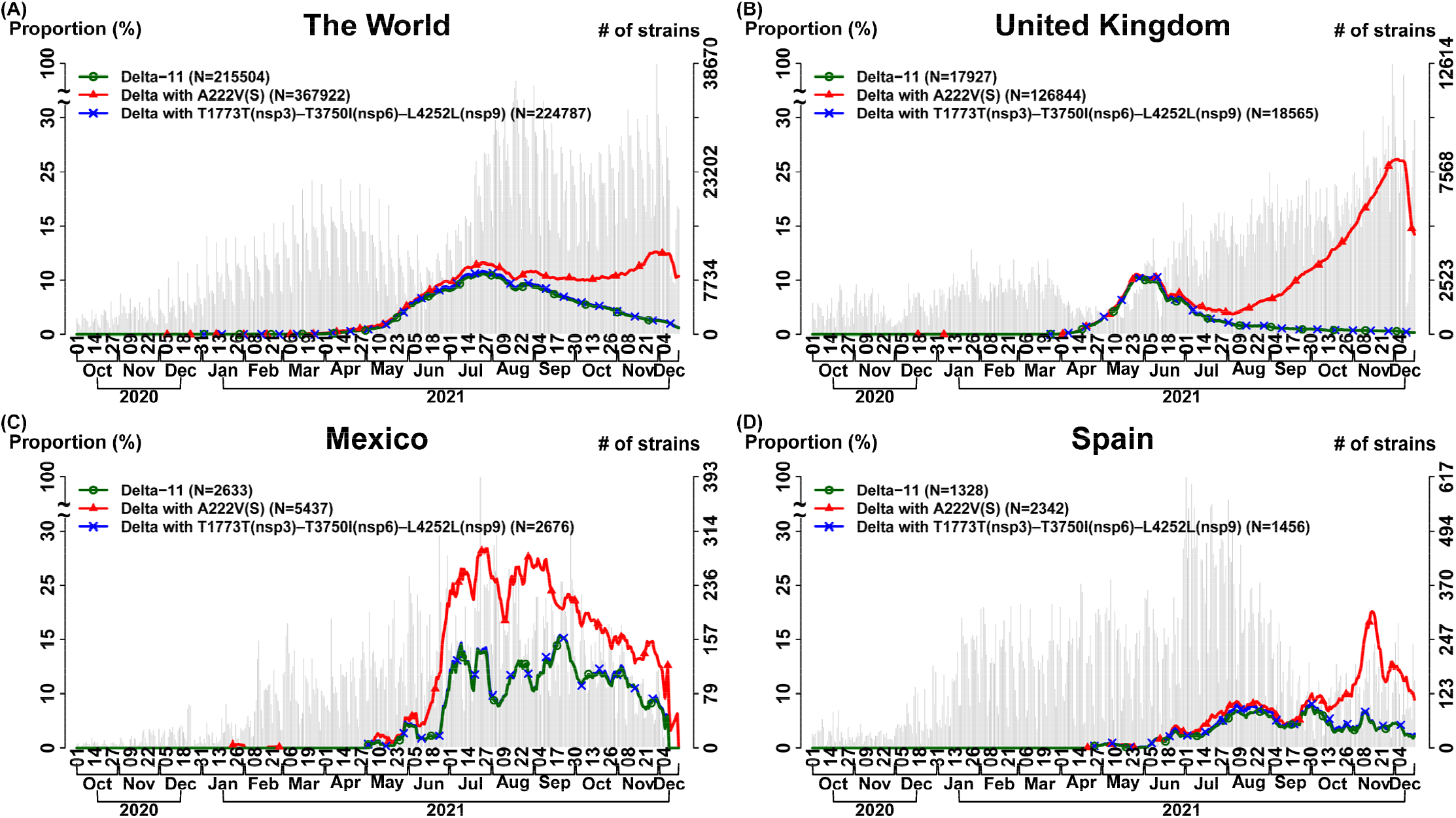
Temporal trajectories for the transmission suppressor SNVs in the Delta variant (n = 6,166K genomes as of 15 Dec, 2021). We analyzed the effects of transmission suppression contributed by a single SNV and a set of SNVs. Delta-11 in **Fig. 2** is illustrated as an example. In each subfigure, three curves indicate the temporal trajectories for three subgroups: (i) “Delta-11” (**Fig. 2B**) carries a full set of transmission suppressor SNVs: T1773T(nsp3)–T3750I(nsp6)–L4252L(nsp9)–A222V(S) (green color and circle symbol); (ii) “Delta–11 with T1773T(nsp3)–T3750I(nsp6)–L4252L(nsp9)” indicates the Delta strains that they carry the signature SNVs similar to Delta-11 but miss a suppressor SNV A222V(S), i.e., the Delta strains which carry only the suppressor triplet T1773T(nsp3)–T3750I(nsp6)–L4252L(nsp9) (blue color and x symbol); (iii) “Delta with A222V(S)” indicates the Delta strains with a transmission suppressor SNV in the spike protein A222V (red color and triangle symbol). The number of total strains per date (gray bar) is displayed with the histogram in the background. “Delta with A222V(S)” has a higher temporal trajectory than “Delta–11”, indicating that the set of four transmission suppressor SNVs work cooperatively and have a stronger synergistic effect in suppressing transmission suppression than a single SNV A222V(S). The curves for “Delta–11” and “Delta–11 with T1773T(nsp3)–T3750I(nsp6)–L4252L(nsp9)” are very close, reflecting that the four suppressor SNVs T1773T(nsp3)–T3750I(nsp6)–L4252L(nsp9)–A222V(S) have a high allelic association, and therefore it’s hard to observe any missing suppressor SNVs from the set of transmission suppressor SNVs. (**A**) **The World**; (**B**) **The United Kingdom**; (**C**) **Mexico**; (**D**) **Spain**.

**Figure 6.**
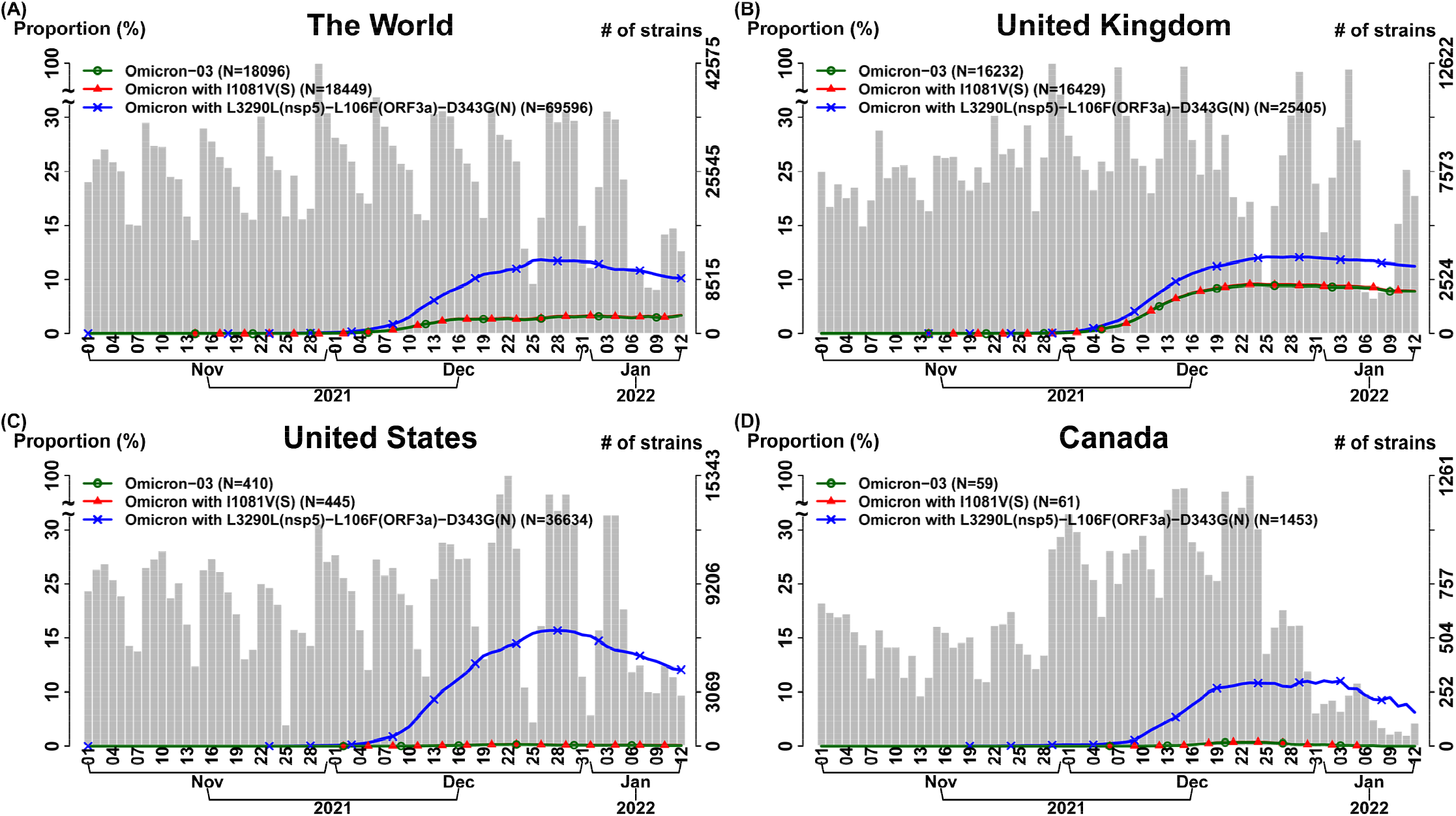
Temporal trajectories for the transmission suppressor SNVs in the Omicron variant (n = 7,026K genomes as of 12, Jan, 2022). We analyzed the effects of transmission suppression contributed by a single SNV and a set of SNVs. Omicron-03 in **Fig. 4** is illustrated as an example. In each subfigure, three curves indicate the temporal trajectories for the three subgroups as follows: (i) “Omicron-03” (**Fig. 4B**) carries a full set of four transmission suppressor SNVs: L3290L(nsp5)–I1081V(S)-L106F(ORF3a)–D343G(N) (green color and circle symbol); (ii) “Omicron-03 with L3290L(nsp5)–L106F(ORF3a)–D343G(N)” indicates the Omicron strains that they carry the signature SNVs similar to Omicron-03 but miss a suppressor SNV I1081V in the spike protein, i.e., it only carries the suppressor triplet L3290L(nsp5)–L106F(ORF3a)–D343G(N) (blue color and x symbol); (iii) “Omicron with I1081V(S)” indicates the Omicron strains carrying a transmission suppressor SNV in the spike protein I1081V (red color and triangle symbol). The number of total strains per date (gray bar) is displayed with the histogram in the background. “Omicron-03” and “Omicron with I1081V(S)” are very close, representing that the suppressor SNV I1081V(S) and triplet L3290L(nsp5)-L106F(ORF3a)-D343G(N) co-appeared in the Omicron variants. They have a lower temporal trajectory compared to the one in “Omicron-03 with L3290L(nsp5)–L106F(ORF3a)–D343G(N)”, indicating that I1081V(S) and/or L3290L(nsp5)–I1081V(S)–L106F(ORF3a)–D343G(N) have an effect in suppressing a viral transmission of Omicron. More data are needed to distinguish that the suppressing effect is contributed by I1081V solely and/or the full set of transmission suppressor SNVs: L3290L(nsp5)–I1081V(S)–L106F(ORF3a)–D343G(N). (**A**) **The World**; (**B**) **The United Kingdom**; (**C**) **The United States**; (**D**) **Canada**.

## Discussion

SARS-CoV-2 is an RNA virus and can readily acquire mutations during the replication process and generate new variants and subtypes. The subtypes arising from the recent common ancestral variant may have highly correlated SNVs but remarkably different genomic sequences and transmission patterns. In this study, we developed a systematic dimension reduction approach to characterizing the viral subtypes based on the correlated SNV sets with allelic association and monitoring their emergence and growth. We also developed a pattern recognition approach to grouping CSSs and detecting the sets of transmission enhancers and suppressors. By analyzing 8 million genome sequences of SARS-CoV-2, we provided real-world evidence for the viral subtypes. The identified subtypes exhibit differential temporal trajectories. The patterns can be characterized by the sets of transmission enhancers and suppressors located both on the spike protein and elsewhere. This highlights the importance of SNVs in both spike and non-spike proteins.

Spike-protein signature SNVs are often used as a proxy for diagnosing variants and explaining increasing viral transmissibility. Our result shows that almost all CSSs contain SNVs on the spike protein (1,053/1,057 = 99.62%), and only 4 CSSs do not contain any defining SNVs on spike. In total, 37.16% of the defining SNVs of a CSS are located in the spike protein, which is larger than the proportion of SNVs in the spike protein in the whole genome (3,822/29,903 = 12.78%). Remarkably, spike-protein is highly related to allelic association; spike-protein is characterized by the high ratio of non-synonymous SNVs vs. synonymous SNVs with allelic association, frequent intergenic allelic association (i.e., Spike-Nucleocapsid association), and frequent intragenic allelic association. These results indicate that spike is constantly under selection and that it is the most important protein coded by the viral genome which determines the overall viral fitness and transmission effectiveness.

The non-spike protein signature SNVs have been commonly found in the important variants, however, their roles have been largely under-appreciated. Our results reveal that the non-spike-proteins signature SNVs provide subtle information for the subtypes of a variant. In addition, the non-spike-proteins signature SNVs are also relevant to the viral transmissibility. Allelic association provides a direction for investigating mechanistic interactions. The non-spike-proteins SNVs can directly elevate or reduce the viral transmissibility through a direct genetic epistasis with other SNVs (i.e., genetic buffering). On the other hand, the observed association of non-spike protein SNVs may be explainable by genetic hitchhiking. However, this explanation is disfavored because we did not find “hitchhikers” of less than the whole set of suppressors. The current vaccines have been designed targeting at the spike protein. Our results suggest that the non-spike-proteins SNVs should also be considered to improve the sensitivity of SARS-CoV-2 variants such as Omicron, Delta, and Alpha to pharmacological intervention.

Our results show that the ratio of nonsynonymous versus synonymous amino acid change (r) is much higher in the transmission enhancer SNVs compared to the suppressor SNVs. Interestingly, the Alpha variant has a high r value (r = 4.5) for the transmission enhancer SNVs, and the Delta variant has an even higher r value (r = 11.5). Moreover, the Delta variant has a r value (r = 1.375) for the transmission suppressor SNVs much lower than the Alpha variant (r = 4). A higher nonsynonymous vs. synonymous proportion may suggest a possible nature selection [18]. The results reveal that the Delta variant may have a higher positive selection in the transmission enhancer SNVs and a lower negative selection in the transmission suppressor SNVs. This partially reflects the dominance of Delta compared to other variants. Compared to the previous variants, Omicron is known as a variant with a significant enrichment of spike signature SNVs. SNVs in the receptor binding domain (RBD) of the spike protein (S) can alter the affinity to the angiotensin converting enzyme 2 (ACE2) receptor and potentially cause vaccine escape [7, 19, 20]. Omicron has exhibited a more rapid transmission than previous Variants of Concern.

Because of a lasting evolution of SARS-CoV-2, there is an unmet need to systematically track the dynamic changes of the viral subtypes, understand their signature SNVs, and improve our understanding and controlling for the COVID-19 pandemic. However, the computation becomes a hurdle when the number of genomes exceeds a million scale. We find that allelic association is a hallmark for an emergence and growth of a subtype and therefore can be employed to detect a strain subtype. Viral strains in a subtype share the signature SNVs with high allelic association (i.e., CSS). A CSS-based approach provides a multilocus analysis that it is more informative than a SNV by SNV analysis. A SNV by SNV analysis ignores allelic association and genetic epistasis, causing an increased false positive in identifying transmission suppressors and an underestimated effect of transmission suppression.

In addition, a large number of SNVs are neutral and do not benefit the fitness gain of SARS-CoV-2 [21]. It is redundant to consider all SNVs in the 30K genome of SARS-CoV-2. The CSS-based analysis addresses this problem. A CSS-based analysis based on a set of correlated signature SNVs significantly overcomes the computational bottleneck in the strain-by-strain and SNV-by-SNV whole-genome analysis.

## Materials and Methods

### Data download and preprocessing

We downloaded and preprocessed 1,180K, 1,589K, 1,872K, 2,215K, 6,316K, 7,205K, and 8,940K whole-genome sequences data from the Global Initiative on Sharing Avian Influenza Data (GISAID) database (https://www.gisaid.org/) on 21 Apr, 19 May, 09 Jun, 07 Jul, 29 Dec in 2021 and 26 Jan, 02 Mar in 2022, respectively (**Fig. S15**). Strain information was extracted from the meta information in GISAID. After discarding the duplicated samples, the samples with an aligned sequence of <29K bases, and the samples without sample recruitment date, it remained the complete sequences of 1,047K, 1,391K, 1,805K, 2,119K, 6,166K, 7,026K, and 8,475K genomes, respectively. Multiple sequence alignment was performed by using MAFFT v.7 [22]. The Wuhan-Hu-1 that the strain was originally isolated in China and had 29,903 nucleotides [23] was employed as the reference genome. Our major sequence analysis discarded two ends (5’ leader and 3’ terminal sequences) and focused on the SNV base positions from 266 to 29,674. Nucleotides different from the Wuhan-Hu-1 strain were assigned as a SNV. Deletions were also detected. Annotation of the SNVs was collected from CNCB (https://bigd.big.ac.cn/ncov/release_genome). Statistical analyses were performed by using our self-developed R codes.

### CSS analysis

CSS subtyping was established based on the proposed analysis procedures (**Fig. S5A**). Matrix representation for a dimension reduction of the proposed CSS analysis procedure is provided (**Fig. S6**). Identification of preliminary SNV groups (PSGs) based on allelic association and determination of CSSs by using an exponential weighted moving average (EWMA) [24] are explained. We calculated variation (allele) frequencies of SNVs in different countries and in the whole world by using a direct allele counting. For the SNVs with a frequency >0.01, we calculated pairwise allelic association for any pairs of SNVs by using the square of the correlation coefficient [25] as follows:

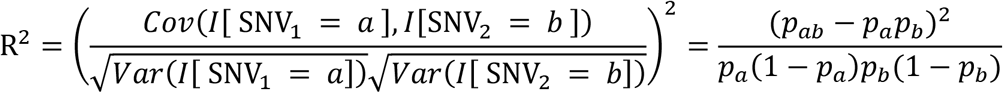

where *Cov* and *Var* indicate covariance and variance, respectively. *I*[*A*] indicates an indicator variable with a value of 1 if event *A* holds, 0 otherwise. Frequency *p_ab_* indicates occurrence frequency (haplotype frequency) with allele *a* at the first SNV and allele *b* at the second SNV. *p_a_* (*p_b_*) indicates the frequency of allele *a* (*b*) at the first (second) SNV.

An algorithm was developed to group SNVs as a preliminary SNV group (PSG) that a PSG contains more than 3 SNVs and all pairwise allelic associations ≥ 0.5. We removed very rare PSGs which contained less than and equal to 20 strains in the last 90 days in a country or in the world. On the one hand, signature SNVs in PSGs were extended if the SNVs on the spike protein had a very high proportion. On the other hand, a subset of PSGs were also regarded as a PSG if the subset had a proportion ≥ 0.1. EWMA control chart for testing H_0_:E(*Z_t_*)≤*μ*_0_ vs. H_*α*_:E(*Z_t_*) >*μ*_0_ was applied to detect correlated SNV sets (CSSs) and track the growth change of a variant over time as follows: Let *Y_t_* denote the temporal proportion at time point t for a PSG. The EWMA statistic

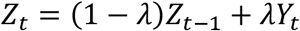

represents a weighted average characteristic of the past and current occurrence proportion of a variant, where *λ* indicates a (smoothing) weight for the current temporal proportion. A PSG was identified as an emerging CSS (i.e., Z_t_ is out-ofcontrol) if

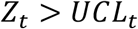

where the upper control limit *UCL_t_ = μ_0_* + 6 -σ(Z_t_) and

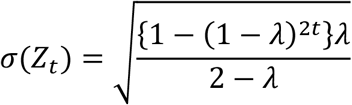

where *σ_γ_* denotes the standard deviation of *Y_t_*. Default *μ_0_* = 0.01 and *λ =* 0.2 were considered. The analysis was conducted by using R package qcc [26].

Once CSSs had been determined, genome sequence of a CSS was determined by substituting the genome of the Wuhan-Hu-1 strain with the signature SNVs with high allelic association of the CSS. A CSS-based phylogenetic analysis (**Fig. S8**) based on maximum parsimony (MP) was conducted by using MEGA X [27]. Subtree-Pruning-Regrafting algorithm [28] was employed for a tree topology search heuristic.

### Detection of transmission enhancer SNVs and transmission suppressor SNVs

Transmission enhancer SNVs and transmission suppressor SNVs were detected by using the proposed procedures (**Fig. S5B**). Decision rule, parameter vector, and parameter updating by using Particle Swarm Optimization [29] are explained below.

On the basis of the identified CSSs and their temporal proportions for a variant, we proposed a decision rule to classify the CSSs into enhanced, suppressed, and undetermined CSSs for a variant. First, we only included the CSSs with a temporal proportion >*θ*_4_ at some time (hitting time) and focused on the temporal proportions after the first hitting time. Second, CSSs were initially grouped if an average difference of their temporal proportions over time was <*θ*_2_. Third, when all CSSs belong to the same group: (i) if the maximum temporal proportion was <*θ*_3_, then these CSSs were classified as the undetermined CSS group. (ii) If the maximum temporal proportion was located between *θ*_3_ and *θ*_4_: (ii-a) we further monitored the slope of the temporal trajectory. Let *D* denote the increment that we subtracted the sum of negative slopes from the sum of positive slopes. If *D* was ≤*θ*_5_, then the CSSs were classified as the undetermined CSS group. (ii-b) If *D* was >*θ*_5_, then the CSSs were classified as the enhanced CSS group. (iii) If the maximum temporal proportion was ≥*θ*_4_, then the CSSs were classified as the enhanced CSS group. Finally, when CSSs were grouped into multiple groups: (i) if the maximum temporal proportion was <*θ*_4_: (i-a) If *D* was ≤*θ*_5_, then these CSSs were classified as the undetermined CSS group. (i-b) If *D* for all CSS were >0_5_, then these CSSs were classified as the enhanced CSS group. (i-c) If *D* for some CSSs were >*θ*_5_ and some CSSs were ≤*θ*_5_, then the former CSSs were classified as the enhanced CSS group and the latter CSSs were classified as the suppressed CSS group. (ii) if the maximum temporal proportion was ≥*θ*_4_, then the CSSs were classified as the enhanced CSS group. Therefore, given a parameter vector ***θ*** = (*θ*_1_, *θ*_2_, *θ*_3_, *θ*_4_, *θ*_5_), CSSs can be classified into enhanced, suppressed, or undetermined CSSs.

An optimal parameter vector is critical for the decision rule (a classifier of CSSs). The parameter vector ***θ*** = (*θ*_1_, *θ*_2_, *θ*_3_, *θ*_4_, *θ*_5_) was updated and optimized by using Particle Swarm Optimization [29] as follows:

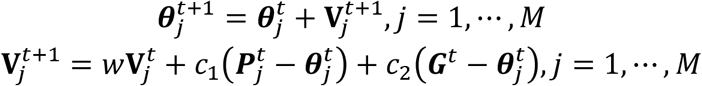

where *w* is the weight of 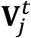 (default: *w* = 0.9); acceleration constant for individuals (“particle”) *c*_1_~Uniform(0, *a*) (default: *a* = 0.2); acceleration constant for population (“swarm”) *c*_2_~Uniform(0, *b*) (default: *b* = 0.2); *M* is the number of initial random particles (default: *M* = 200). 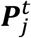 is the best state for individual *j* at iteration *t* and **G**^*t*^ is the best state for population at iteration *t*. In each updating of 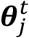 and 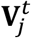, the best states 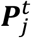 and ***G**^f^* were updated simultaneously as follows:

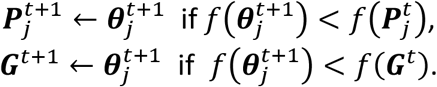

Here we considered an objective function (i.e., misclassification frequency in *n* CSSs) as follows:

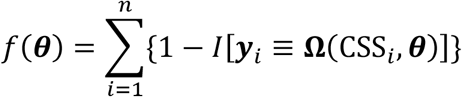

where ***y**_i_* and **Ω**(*CSS_i_*, ***θ***) indicate the true and predicted states (i.e., “enhancer”, “suppressor”, or “undetermined”) of the *i*-th CSS given a parameter vector ***θ***. The true state of a CSS was initially determined based on a heuristic discussion about the pattern of the temporal trajectory of the CSS in a multidisciplinary expert team. The predicted state of a CSS was obtained according to the aforementioned decision rule given a parameter vector ***θ***. For a CSS, if the true and predicted states are identical, then the misclassification error 1 – *I*[***y**_i_*≡**Ω**(*CSS_i_*,***θ***)] is 1, otherwise, 0. The parameter updating procedure was iterated to minimize the misclassification error (*f*(***θ***)). The iteration was stop if: 1) *f*(***G**^t^*’) reached the minimum of *f* (In our case, 0 is the minimum of *f*, this represents every predicted state of CSSs is the true state.) or 2) it reached the maximum number of iterations (default: 25), ***G**^f^* is the optimal estimator of *θ.* The optimization was performed by using R package pso [30].

The optimal parameter vector was plugged into the decision rule to find the candidate transmission enhancers and suppressors for variant(s) and their corresponding signature SNVs. Finally, the results need to be confirmed in at least 80% of the studied countries and further confirmed in a later dataset with a larger sample size. The established decision rule can be directly apply to determine the CSS states, or serve as a good initial in an adaptive decision rule for more other variants.

## Acknowledgments

We thank Drs. Ling-Jyh Chen, Shang-Te Danny Hsu, and Kay-Hooi Khoo for their useful discussions. We thank Global Initiative on Sharing Avian Influenza Data (GISAID) for providing the rich data for the genome sequences of SARS-CoV-2. We thank National Center for High-performance Computing of National Applied Research Laboratories of Taiwan for providing computational resources.

## Data and Code Availability

The analysis data are available publicly at Global Initiative on Sharing Avian Influenza Data (GISAID) (https://www.gisaid.org/). The codes for our analysis are available upon request.

## Highlight Points

- We develop a dimension reduction method based on correlated SNV sets (CSSs) for a viral subtyping and pattern recognition for transmission dynamics.
- Multiple analyses of two – eight million SARS-CoV-2 genomes identify transmission enhancer SNVs and suppressor SNVs for the contagious variants Omicron, Delta, and Alpha.
- The finding is further confirmed in the subsequent analyses with larger sample sizes and in more countries.
- This study improves our understanding about SARS-CoV-2 and controlling for the COVID-19 pandemic.

## Authors’ Contributions

J.C.L. and H.C.Y. conceived the study, designed the research, and wrote the manuscript. J.H.W., C.T.Y., Y.C.L., H.N.H., B.W.C., H.C.L., and C.h.C. performed data analysis. J.C.L. and H.C.Y. supervised the project.

---

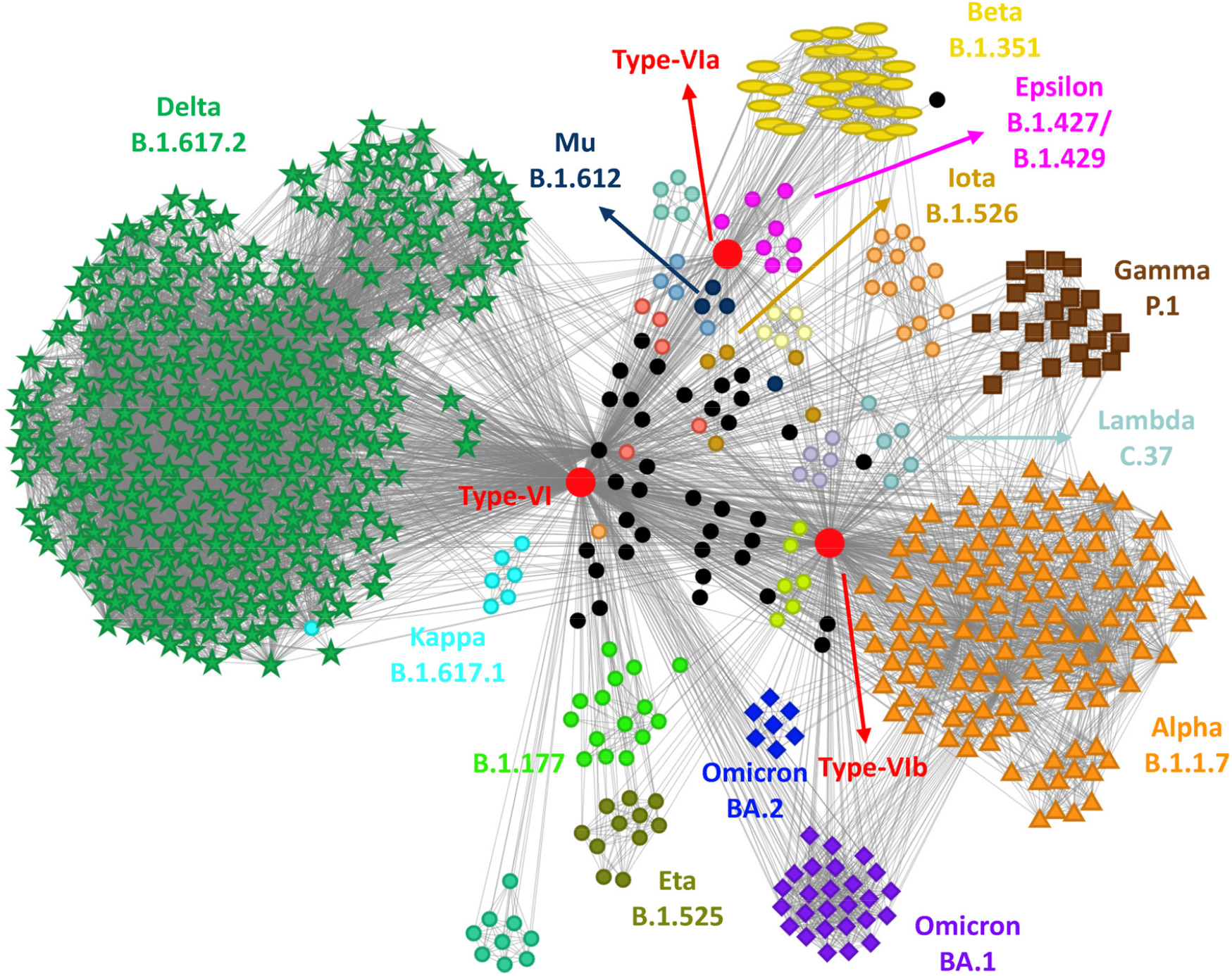
Cover. Network of SARS-CoV-2 variants based on the correlated SNV sets (CSSs) (n = 7,026K genomes as of 12 Jan, 2022). In the network of CSSs, each node indicates one CSS. A node that it has many links to other nodes is called a hub of CSSs. The hub of CSSs with the highest degree is the three red nodes in the center. The hubs include the Type VI variant and the two sub-VI groups: VIa and VIb (Yang et al., 2020, PNAS). Nodes are connected if the CSSs share strains. Variant type for a CSS is assigned according to the variant type of the majority of the viral strains in the CSS. The same variant types are displayed with the same node color. Special node symbols for the five Variants of Concern are given – Alpha (aka B.1.1.7, triangle), Beta (aka B.1.351, ellipse), Gamma (aka P.1, square), Delta (aka B.1.617.2, asterisk), and Omicron (aka B.1.1.529, diamond) including two sub-lineages BA.1 and BA.2. Other variants share the same symbol circle. Representative communities in correspondence to the well-known variants are formed, including Alpha (aka B.1.1.7), Beta (aka B.1.351), Gamma (aka P.1), Delta (aka B.1.617.2), Omicron (aka B.1.1.529), Eta (aka B.1.525), Iota (aka B.1.526), Kappa (aka B.1.617.1), Lambda (aka C.37), Epsilon (aka B.1.429), and Mu (aka B.1.621). Contrast to the well-recognized variants, CSSs provide further information about the subtypes of the variants and also the abundance and heterogeneity of the CSS communities. Interestingly, if two distant CSSs have a link not through the hubs, it suggests that some of the strains in the connected CSSs may have undergone genetic recombination.

