## Supplemental information for "Subtyping of major SARS-CoV-2 variants reveals different transmission dynamics"


| | | |
|---|---|---|
| **Figure S1.** | Allelic association and co-mutation (haplotype) frequency (n = 2,119K) …………………… | 2 |
| **Figure S2.** | Genomic distributions of allelic association (n = 2,119K) ………………………………………… | 5 |
| **Figure S3.** | Intragenic and intergenic allelic associations (n = 2,119K) ……………………………………… | 7 |
| **Figure S4.** | Intergenic allelic association between the spike and nucleocapsid proteins (n = 2,119K) ……………………………………………………………………………………………………………… | 11 |
| **Figure S5.** | Flow charts of the proposed correlated SNV sets (CSSs) analysis and auto-detection algorithm for transmission enhancer SNVs and transmission suppressor SNVs ……….. | 14 |
| **Figure S6.** | Matrix representation of dimension reduction by using correlated SNV sets (CSSs) …. | 16 |
| **Figure S7.** | Number of correlated SNV sets (CSSs) for the well-known variants in seven datasets. | 18 |
| **Figure S8.** | Phylogenetic dendrogram of correlated SNV sets (CSSs) (n = 2,119K) ……………………… | 19 |
| **Figure S9.** | Differential temporal trajectories for the eleven Delta subtypes with transmission enhancer and/or suppressor SNVs in many countries (n = 2,119K) ………………………….. | 20 |
| **Figure S10.** | Differential temporal trajectories for the eleven Delta subtypes with transmission enhancer and/or suppressor SNVs in many countries (n = 6,166K) ………………………….. | 23 |
| **Figure S11.** | Differential temporal trajectories for the three Alpha subtypes with transmission enhancer and/or suppressor SNVs in many countries (n = 2,119K) ………………………….. | 26 |
| **Figure S12.** | Differential temporal trajectories for the three Omicron subtypes with transmission enhancer and/or suppressor SNVs in many countries (n = 7,026K) ………………………….. | 29 |
| **Figure S13.** | Differential temporal trajectories for the three Omicron subtypes with transmission enhancer and/or suppressor SNVs in many countries (n = 8,475K) …………………………. | 32 |
| **Figure S14.** | Differential temporal trajectories for the seven Omicron subtypes with transmission enhancer and/or suppressor SNVs in a larger sample size (n = 8,475K) ……………………. | 34 |
| **Figure S15.** | Data download and preprocessing ……………………………………………………………………… | 40 |
| **Table S1.** | Proportions of intragenic and intergenic allelic association (n = 2,119K) …………………. | 41 |
| **Table S2.** | Distribution of the viral strains and SNVs represented by correlated SNV sets (CSSs) (n = 2,119K) ……………………………………………………………………………………………………… | 42 |
| **Supp Text 1.** | Allelic association is a characteristic of rising variants ……………………………………….. | 43 |
| **Supp Text 2.** | Using correlated SNV sets (CSSs) to subtype existing variants …………………………………. | 44 |

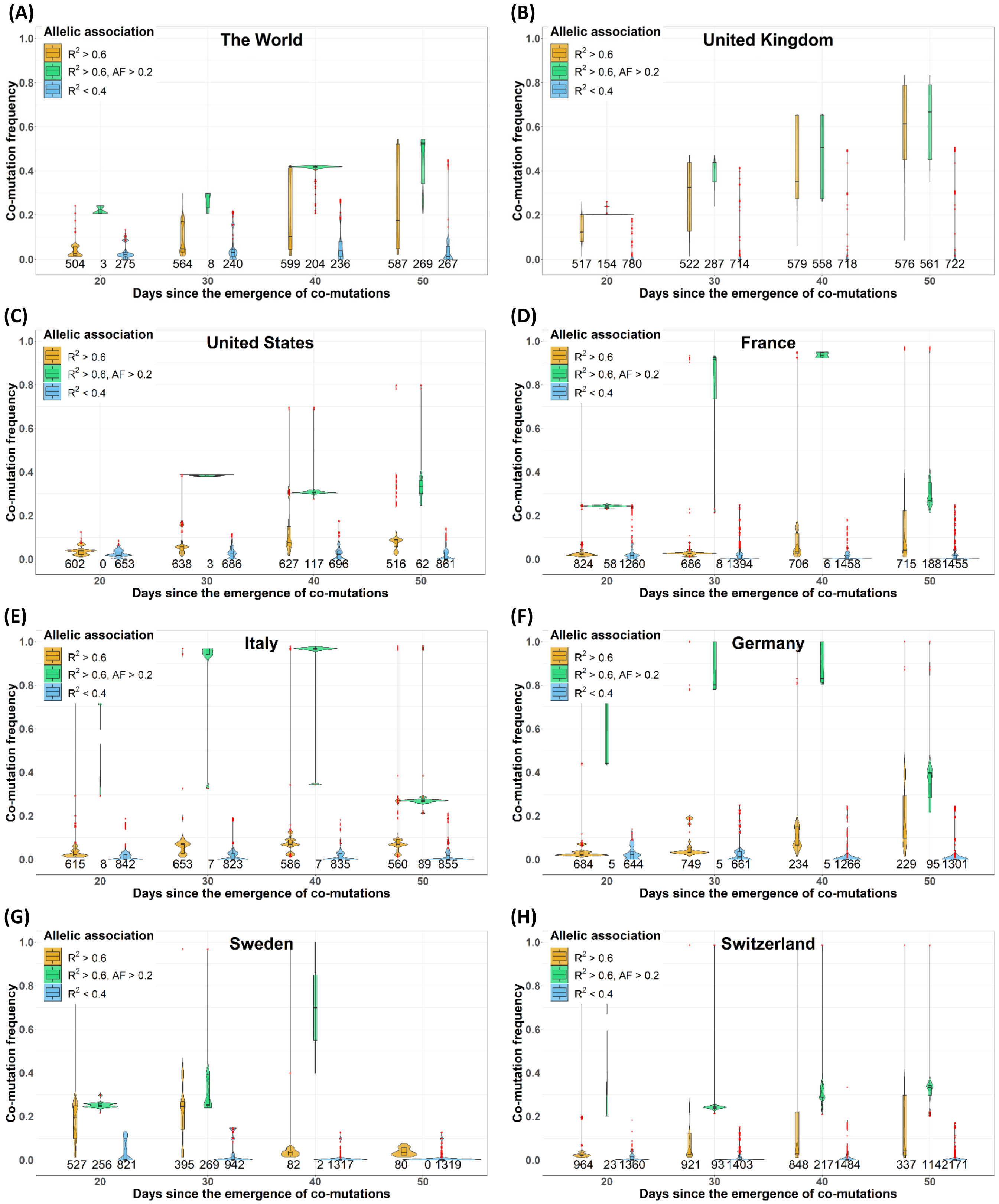
(A)
The World
(B)
United Kingdom
(C)
United States
(D)
France
(E)
Italy
(F)
Germany
(G)
Sweden
(H)
Switzerland
Allelic association
$R^2 > 0.6$
$R^2 > 0.6$, AF > 0.2
$R^2 < 0.4$
Co-mutation frequency
Days since the emergence of co-mutations
504 3 275 564 8 240 599 204 236 587 269 267
517 154 780 522 287 714 579 558 718 576 561 722
602 0 653 638 3 686 627 117 696 516 62 861
824 58 1260 686 8 1394 706 6 1458 715 188 1455
615 8 842 653 7 823 586 7 835 560 89 855
684 5 644 749 5 661 234 5 1266 229 95 1301
527 256 821 395 269 942 82 2 1317 80 0 1319
964 23 1360 921 93 1403 848 217 1484 337 114 2171

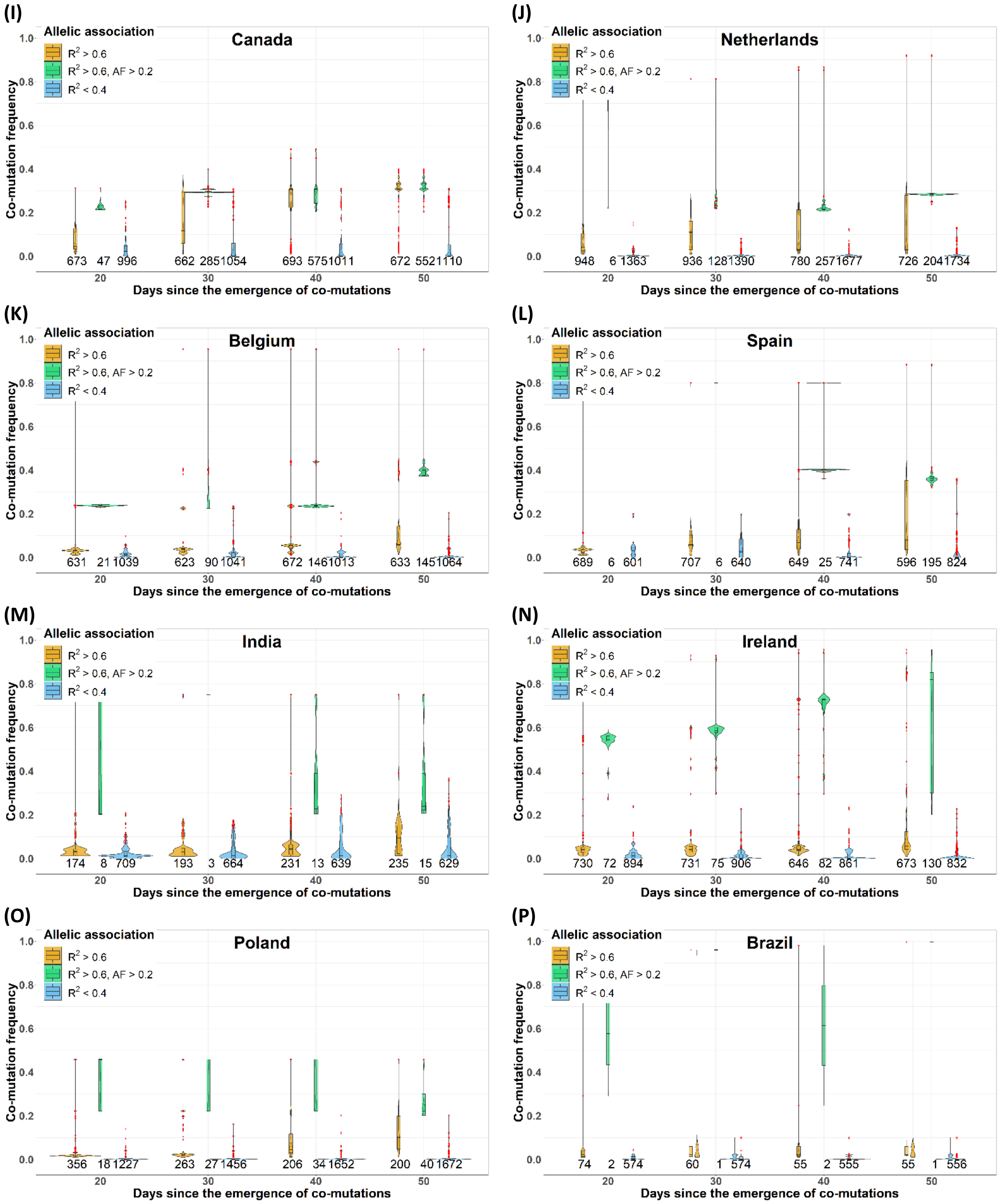
(I)
Canada
Allelic association
R2 > 0.6
R2 > 0.6, AF > 0.2
R2 < 0.4
Co-mutation frequency
673 47 996 662 2851054 693 5751011 672 5521110
Days since the emergence of co-mutations
(J)
Netherlands
936 1281390 948 6 1363 780 2571677 726 2041734
(K)
Belgium
631 21 1039 623 90 1041 672 1461013 633 1451064
(L)
Spain
689 6 601 707 6 640 649 25 741 596 195 824
(M)
India
174 8 709 193 3 664 231 13 639 235 15 629
(N)
Ireland
730 72 894 731 75 906 646 82 861 673 130 832
(O)
Poland
356 18 1227 263 27 1456 206 34 1652 200 40 1672
(P)
Brazil
74 2 574 60 1 574 55 2 555 55 1 556

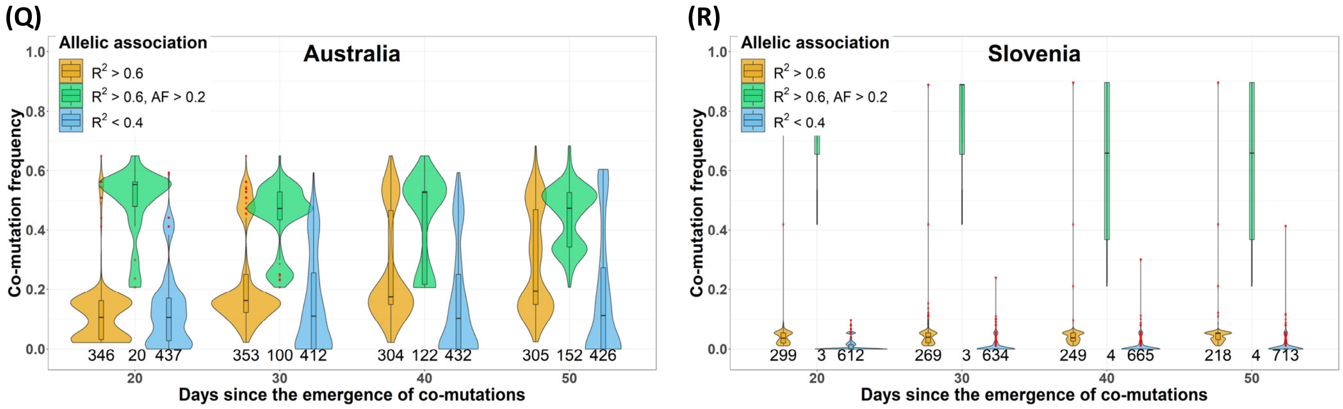


**Figure S1. Allelic association and co-mutation (haplotype) frequency (n = 2,119K).** The relationship between the allelic association and co-mutation frequency of the pairs of SNVs were examined based on the genome sequences of 2,119K SARS-CoV-2 viral strains collected in the GISAID (https://www.gisaid.org/). Only SNVs with a variation frequency > 0.01 were included. Four durations in time, from 20d to 50d with an increment of 10d, since a pair of SNVs started to accumulate its co-mutation frequency were considered. In each duration in time, allelic association $R^2$ and co-mutation frequency of pairs of SNVs were calculated. Pairs of SNVs were divided into the high-allelic-association group with $R^2 > 0.60$ (left-hand side, orange violin plot), $R^2 > 0.60$ and the maximum co-mutation frequency > 0.2 (middle, green violin plot), and low-allelic-association group with $R^2 < 0.4$ (right-hand side, blue violin plot). The vertical axis is the maximum co-mutation frequency ranged from 0 to 1. The horizontal axis is the time duration (days) ranged from 20d to 50d with an increment of 10d. The value beneath a violin plot indicates the number of the pairs of SNVs used for drawing the violin plot. We find that the co-mutation frequencies of pairs of SNVs start to increase only after that $R^2$ increases or has attained a high level. In other words, the necessary condition for an increase of the co-mutation frequency is an existence of allelic association. For the cases of a low allelic association, the co-mutation frequency is rarely increased to a high level. A few exceptions occur if a pair of SNVs are located in a nearby region, or the two SNVs occur sequentially rather than simultaneously. In other words, when one of the mutations already existed in the population, a sequential mutation weakens the allelic association of the two SNVs. The observed pattern reveals that allelic association of SNVs is the driving force for an increasing co-mutation frequency. In other words, a co-occurrence of SNVs benefits to their transmission to the next strains compared to a case of a single or sequential mutation in the most of the cases. (**A**) The World; (**B**) The United Kingdom; (**C**) The United States; (**D**) France; (**E**) Italy; (**F**) Germany; (**G**) Sweden; (**H**) Switzerland; (**I**) Canada; (**J**) The Netherlands; (**K**) Belgium; (**L**) Spain; (**M**) India; (**N**) Ireland; (**O**) Poland; (**P**) Brazil; (**Q**) Australia; (**R**) Slovenia.

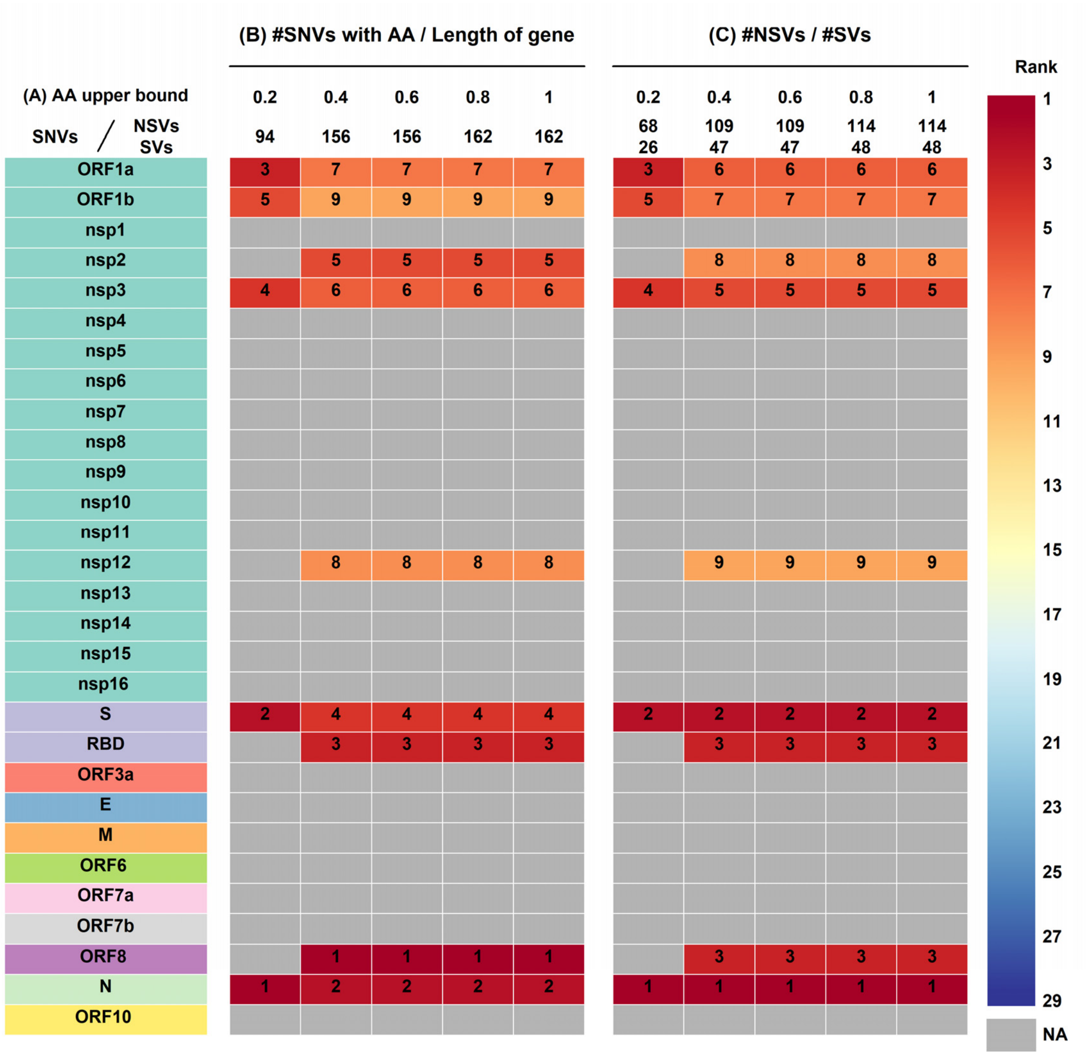

**Figure S2. Genomic distributions of allelic association (n = 2,119K).** (A) The names of genomic regions (protein symbols) are listed and followed by the two heatmaps. (B) The first heatmap shows the genomic distribution of pairs of SNVs with allelic association (A.A.) within each genomic region. Six lower bounds from 0.2 to 1 with an increment of 0.2 were considered for the coefficient of allelic association R2. The numbers of the pairs of SNVs with allelic association are listed beneath the values of the lower bound. The values in the heatmap were calculated as follows: "the number of the pairs of SNVs with allelic association" was divided by "the number of SNVs in a specific protein region (i.e., the length of a protein)". Then the ratios were ranked across different protein regions. A small-value rank indicates that the genomic region has a higher proportion of pairs of SNVs with an intragenic allelic association. The results show that ORF8, N (the nucleocapsid protein), S (the spike protein) especially for RBD (receptor binding domain), S, and nsp2 (the non-structural protein 2) are the top genomic regions that they have a high proportion of intra-gene allelic association. The proportions are 1.64%, 1.27%, 1.03%, 0.76%, and 0.42%, respectively. (C) In the second heatmap, the numbers of non-synonymous SNVs (NSVs) and synonymous SNVs (SVs) with allelic association with other SNVs are listed beneath the values of the lower bound of R2. The values in the heatmap were calculated as follows: "the numbers of non-synonymous SNVs with allelic association" was divided by "the numbers of synonymous SNVs with allelic association". Then the ratios were ranked across different protein regions. A small-value rank indicates that the genomic region has a higher ratio of non-synonymous vs. synonymous SNVs that they have allelic association with other SNVs in the same protein. "NA" indicates that the number of SNVs in allelic association in the protein region is not greater than 5. The results show that N (the nucleocapsid protein), S (the spike protein), RBD (receptor binding domain) in S, ORF8, and nsp3 have the highest ratio of non-synonymous SNVs vs. synonymous SNVs with allelic association. The ratios are 15, 8.67, 5, 5, and 1.5, respectively.

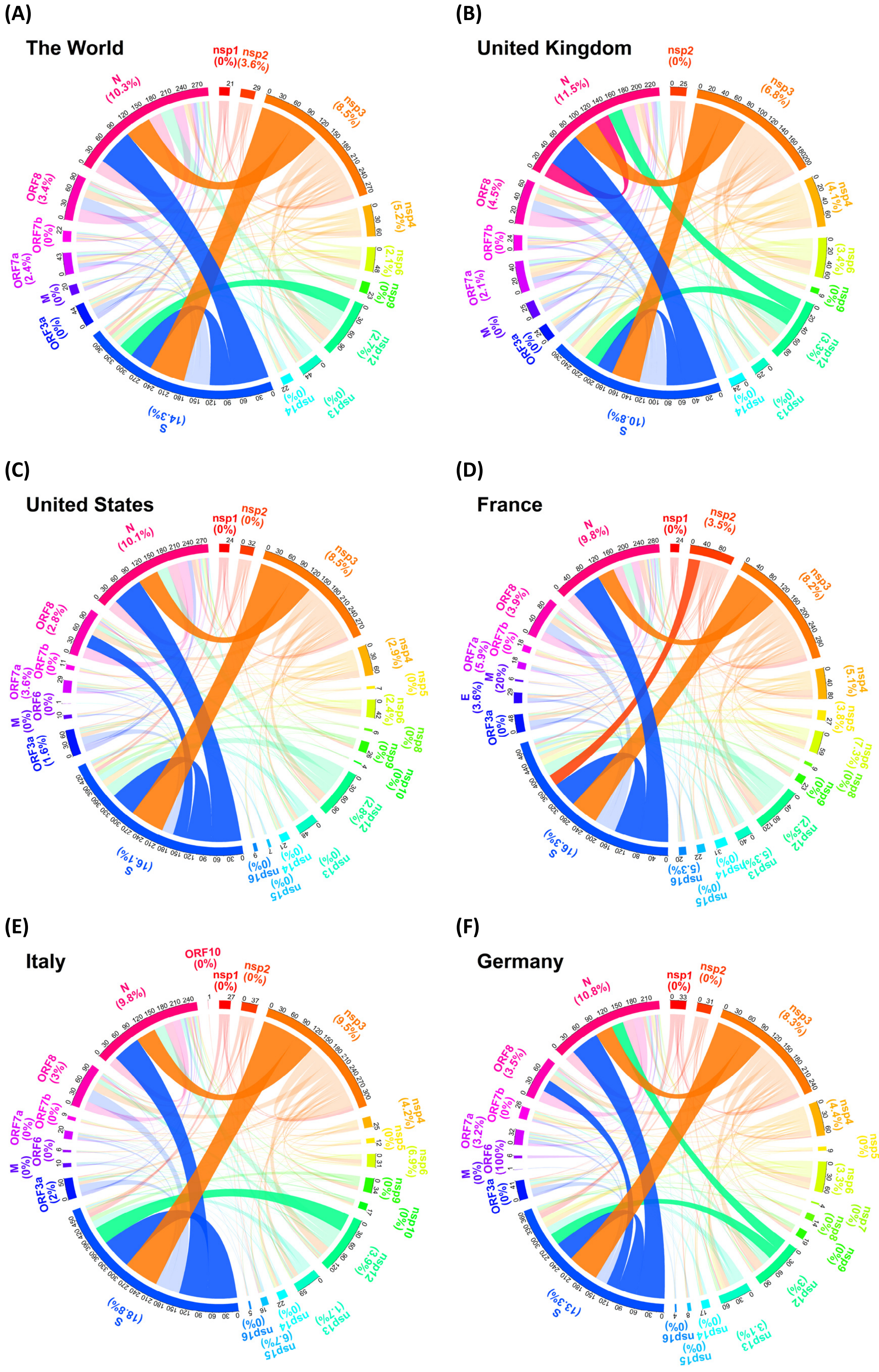

(A)
The World
(B)
United Kingdom
(C)
United States
(D)
France
(E)
Italy
(F)
Germany

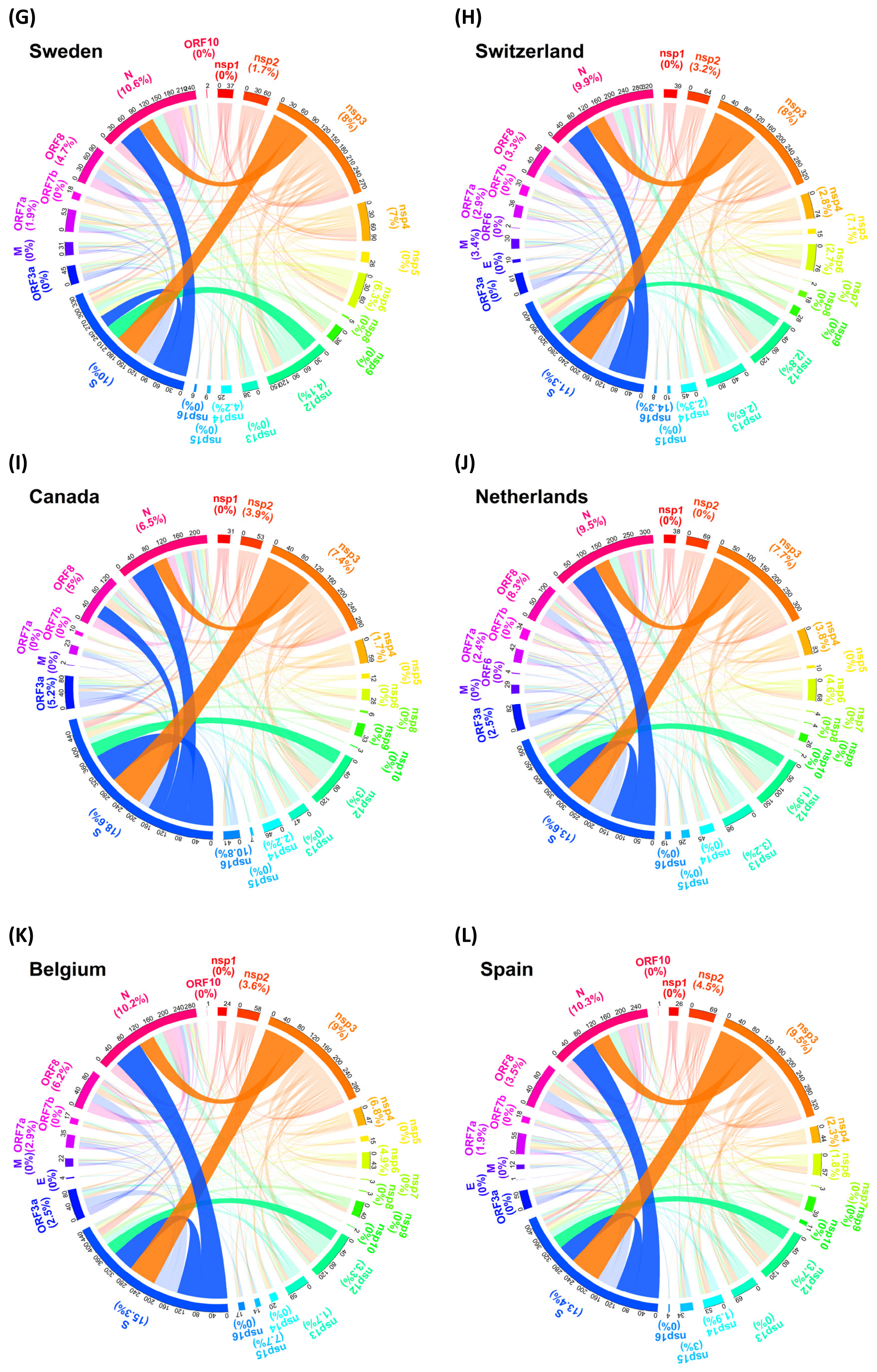
(G)
Sweden
(H)
Switzerland
(I)
Canada
(J)
Netherlands
(K)
Belgium
(L)
Spain

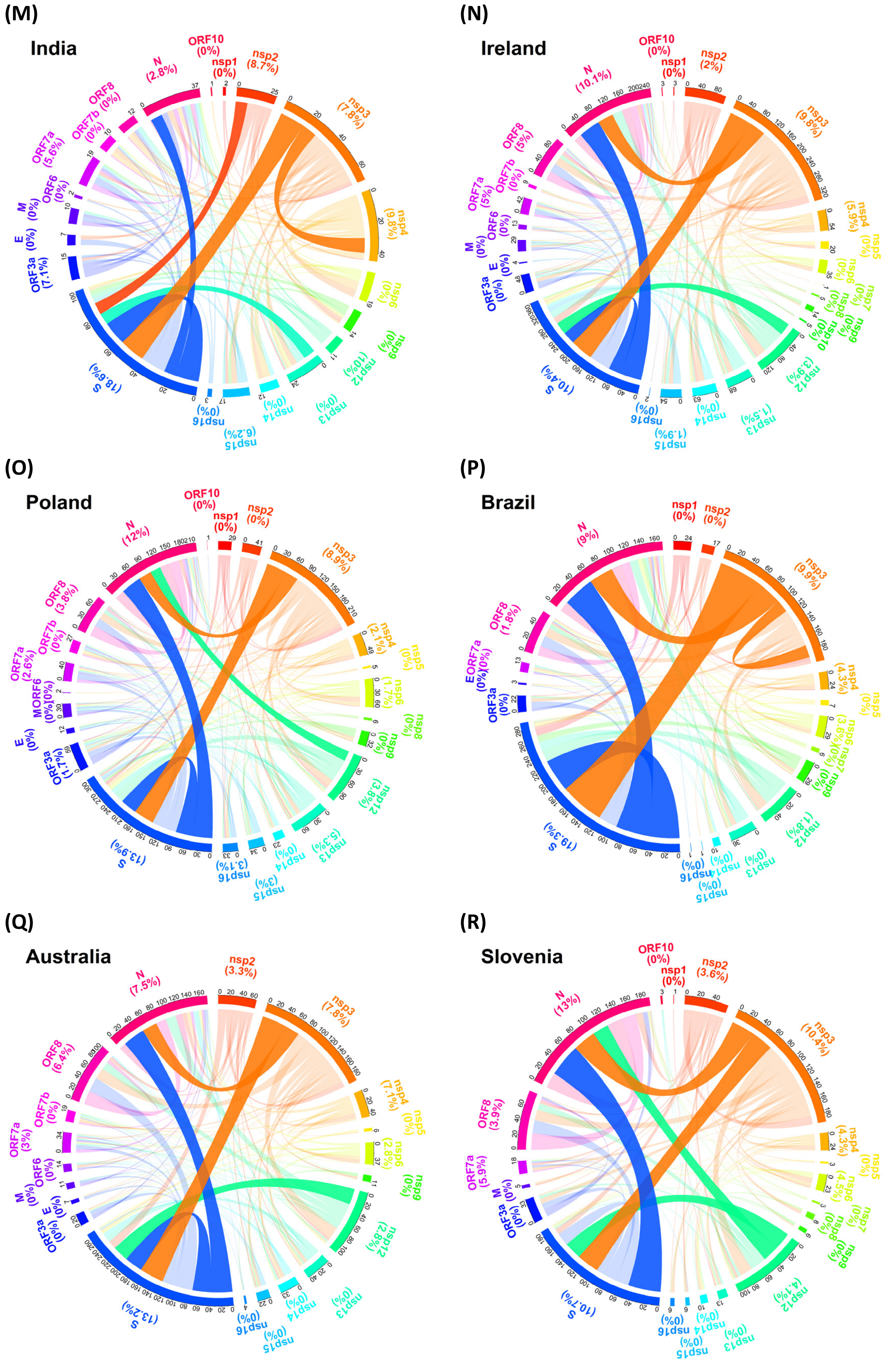
(M)
India
(N)
Ireland
(O)
Poland
(P)
Brazil
(Q)
Australia
(R)
Slovenia

**Figure S3. Intragenic and intergenic allelic associations (n = 2,119K).** In a chord diagram, a line is connected if two SNVs within a gene or between two genes have allelic association. Genes are displayed with different colors. The outer arc length reflects the number of SNVs with allelic association for a gene. Among the proteins involved in intergenic allelic association, spike-nucleocapsid allelic association are the most frequent, e.g., a high frequency of intergenic allelic association between the spike and nucleocapsid proteins. Spike protein is also the region that exhibits the most abundant SNVs that they have intragenic allelic association. The statistics are provided (Table S1). The results of allelic association reveal that the SNVs in the spike protein pivotally involve in inter- and intra-genic allelic associations. Spike protein SNVs has appealed significant attention because the SNVs directly relate to a viral binding and entry to the host cells so as to an increased viral transmissibility. (A) The World; (B) The United Kingdom; (C) The United States; (D) France; (E) Italy; (F) Germany; (G) Sweden; (H) Switzerland; (I) Canada; (J) The Netherlands; (K) Belgium; (L) Spain; (M) India; (N) Ireland; (O) Poland; (P) Brazil; (Q) Australia; (R) Slovenia.

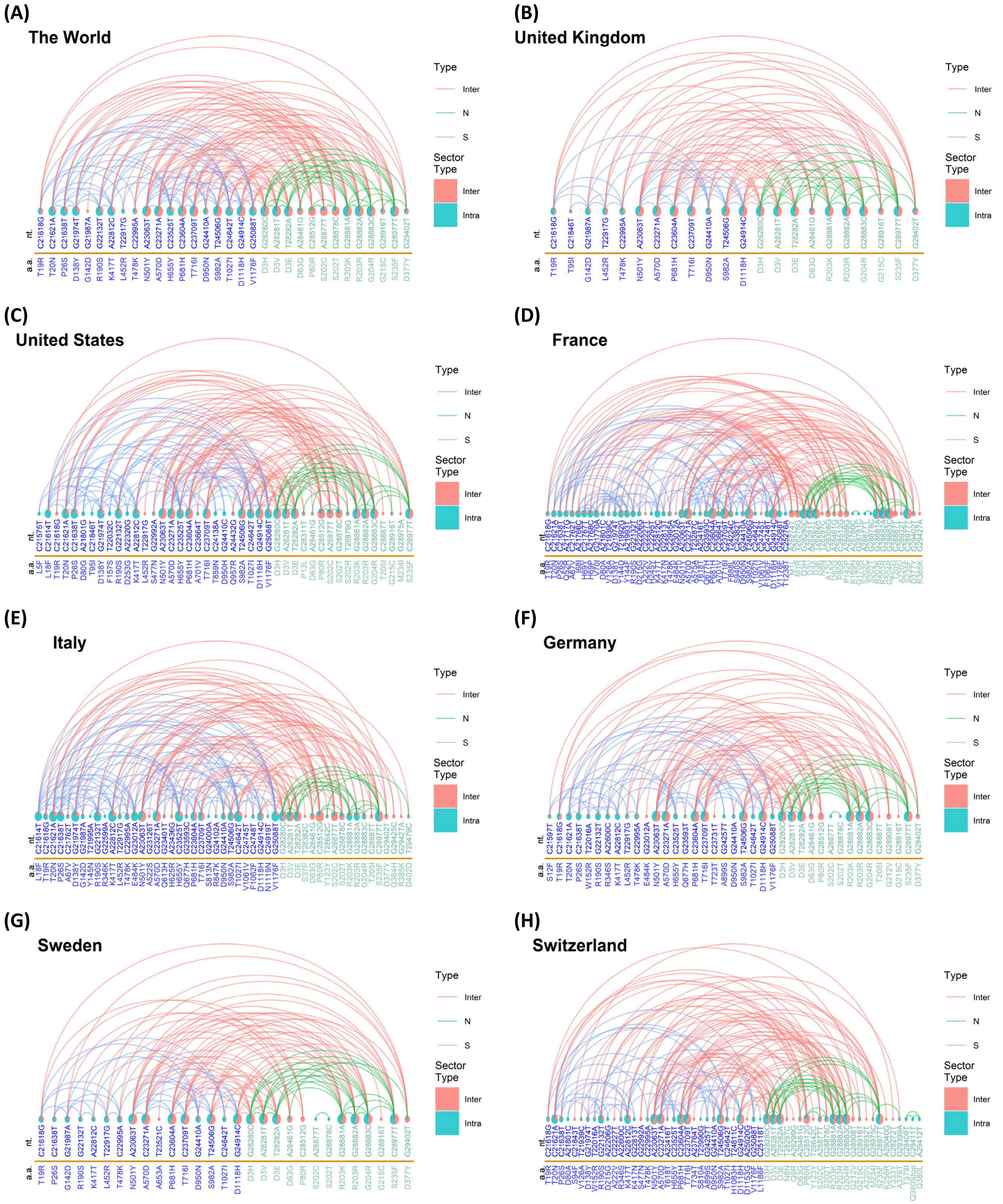
(A)
The World
(B)
United Kingdom
(C)
United States
(D)
France
(E)
Italy
(F)
Germany
(G)
Sweden
(H)
Switzerland
Type
Inter
N
S
Sector Type
Inter
Intra
nt.
a.a.

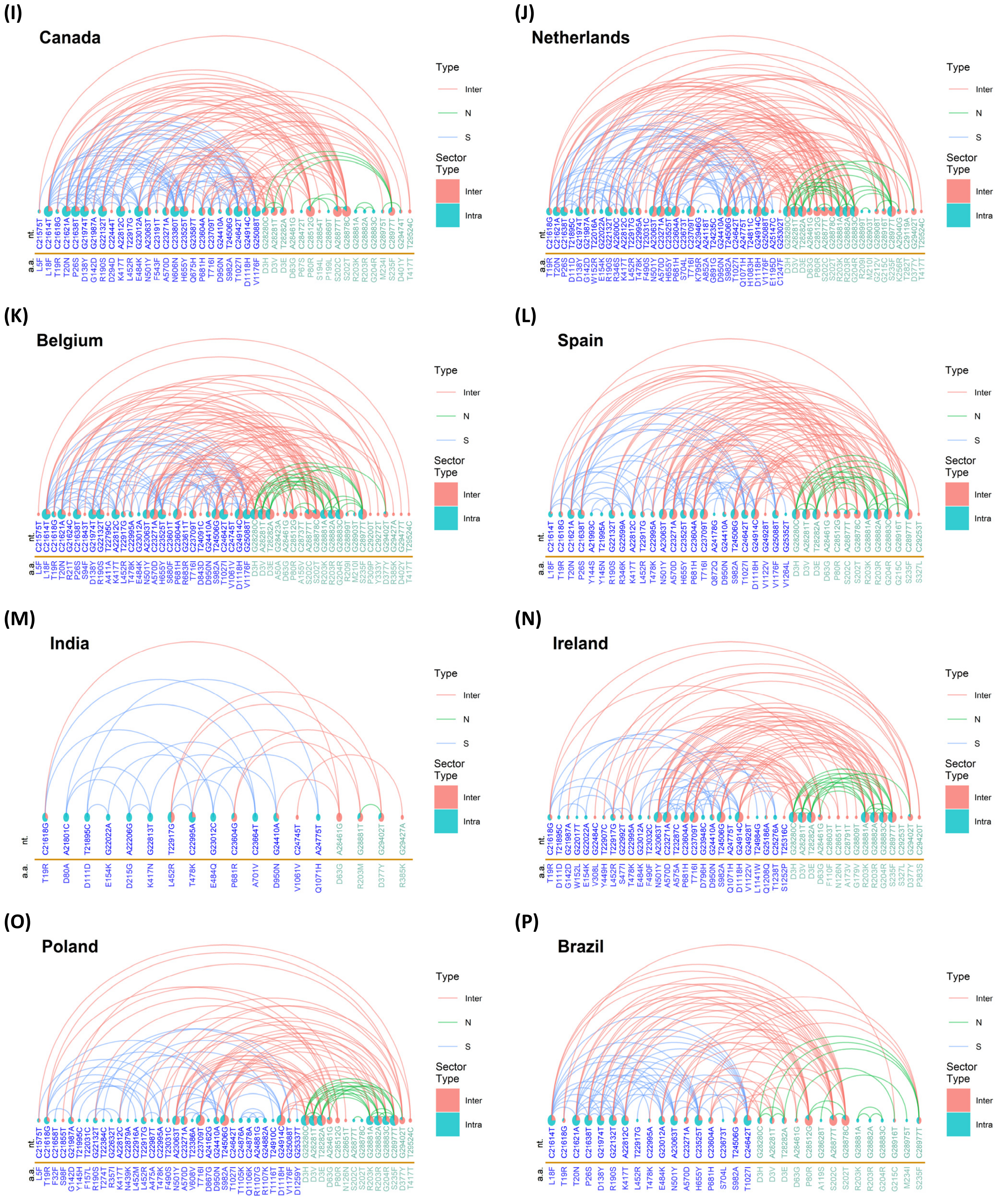

(I)
Canada
(J)
Netherlands
(K)
Belgium
(L)
Spain
(M)
India
(N)
Ireland
(O)
Poland
(P)
Brazil
Type
Inter
N
S
Sector Type
Inter
Intra
nt.
a.a.

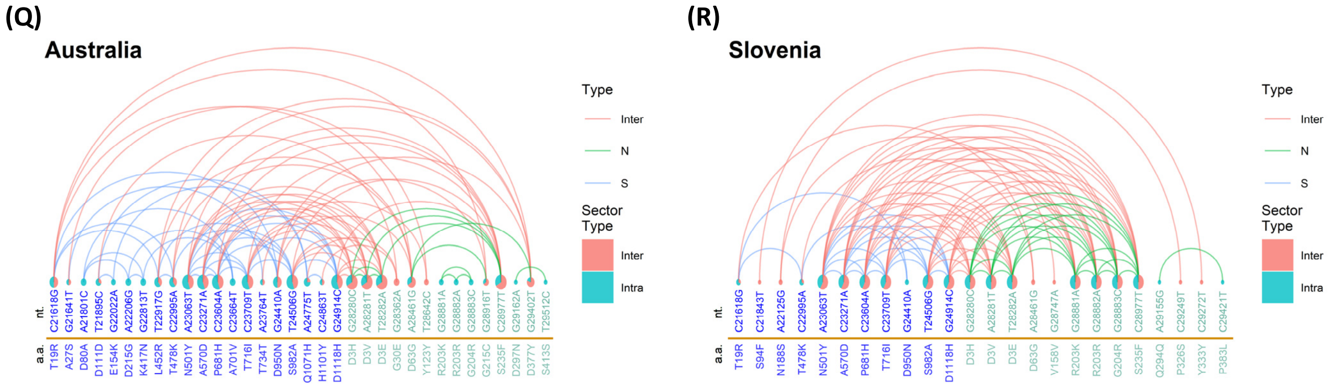


**Figure S4. Intergenic allelic association between the spike and nucleocapsid proteins (n = 2,119K).** Intragenic allelic associations for the SNVs in the spike protein (S) are indicated by a blue link. Intragenic allelic associations for the SNVs in the nucleocapsid protein (N) are indicated by a green link. Intergenic allelic association between the spike and nucleocapsid proteins are indicated by a red link. For each SNV, pie chart reflects the proportions of intragenic (red) and intergenic (green) allelic associations. Among the proteins involved in intergenic allelic association, spike-nucleocapsid allelic associations are the most frequent, e.g., a high frequency of intergenic allelic association between the spike and nucleocapsid proteins. (**A**) The World; (**B**) The United Kingdom; (**C**) The United States; (**D**) France; (**E**) Italy; (**F**) Germany; (**G**) Sweden; (**H**) Switzerland; (**I**) Canada; (**J**) The Netherlands; (**K**) Belgium; (**L**) Spain; (**M**) India; (**N**) Ireland; (**O**) Poland; (**P**) Brazil; (**Q**) Australia; (**R**) Slovenia.

(A)

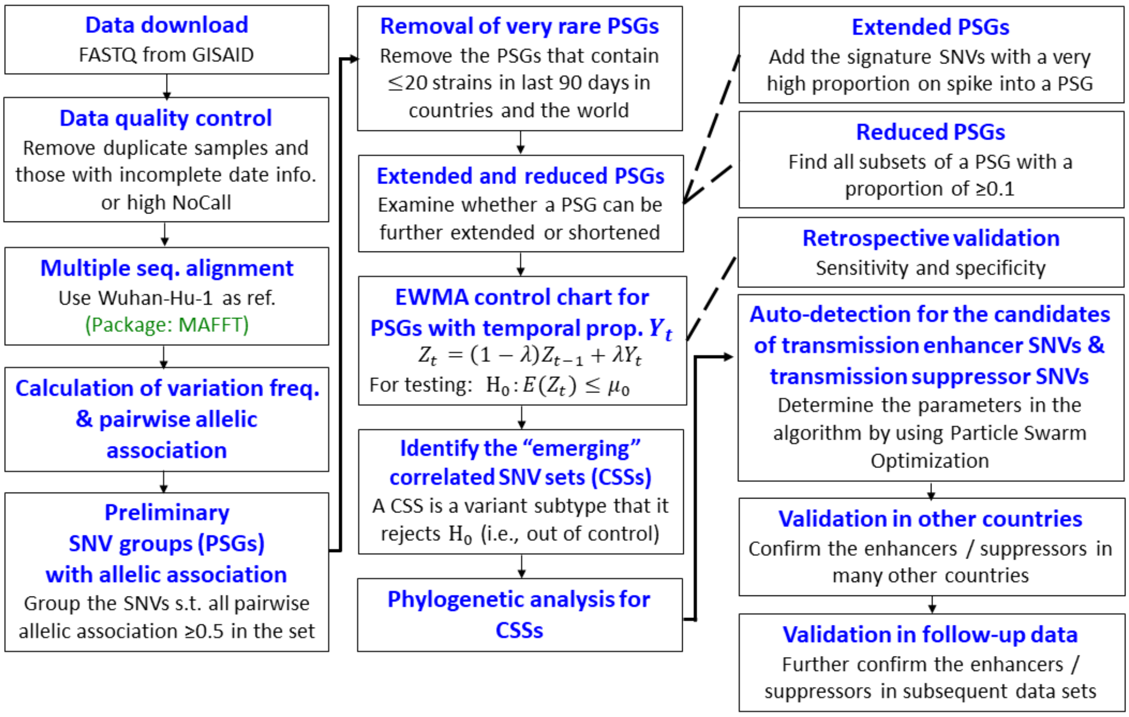
Data download
FASTQ from GISAID
Data quality control
Remove duplicate samples and those with incomplete date info. or high NoCall
Multiple seq. alignment
Use Wuhan-Hu-1 as ref.
(Package: MAFFT)
Calculation of variation freq. & pairwise allelic association
Preliminary SNV groups (PSGs) with allelic association
Group the SNVs s.t. all pairwise allelic association ≥0.5 in the set
Removal of very rare PSGs
Remove the PSGs that contain ≤20 strains in last 90 days in countries and the world
Extended and reduced PSGs
Examine whether a PSG can be further extended or shortened
EWMA control chart for PSGs with temporal prop. $Y_t$
$Z_t = (1-\lambda)Z_{t-1} + \lambda Y_t$
For testing: $H_0: E(Z_t) \le \mu_0$
Identify the "emerging" correlated SNV sets (CSSs)
A CSS is a variant subtype that it rejects $H_0$ (i.e., out of control)
Phylogenetic analysis for CSSs
Extended PSGs
Add the signature SNVs with a very high proportion on spike into a PSG
Reduced PSGs
Find all subsets of a PSG with a proportion of ≥0.1
Retrospective validation
Sensitivity and specificity
Auto-detection for the candidates of transmission enhancer SNVs & transmission suppressor SNVs
Determine the parameters in the algorithm by using Particle Swarm Optimization
Validation in other countries
Confirm the enhancers / suppressors in many other countries
Validation in follow-up data
Further confirm the enhancers / suppressors in subsequent data sets

(B)

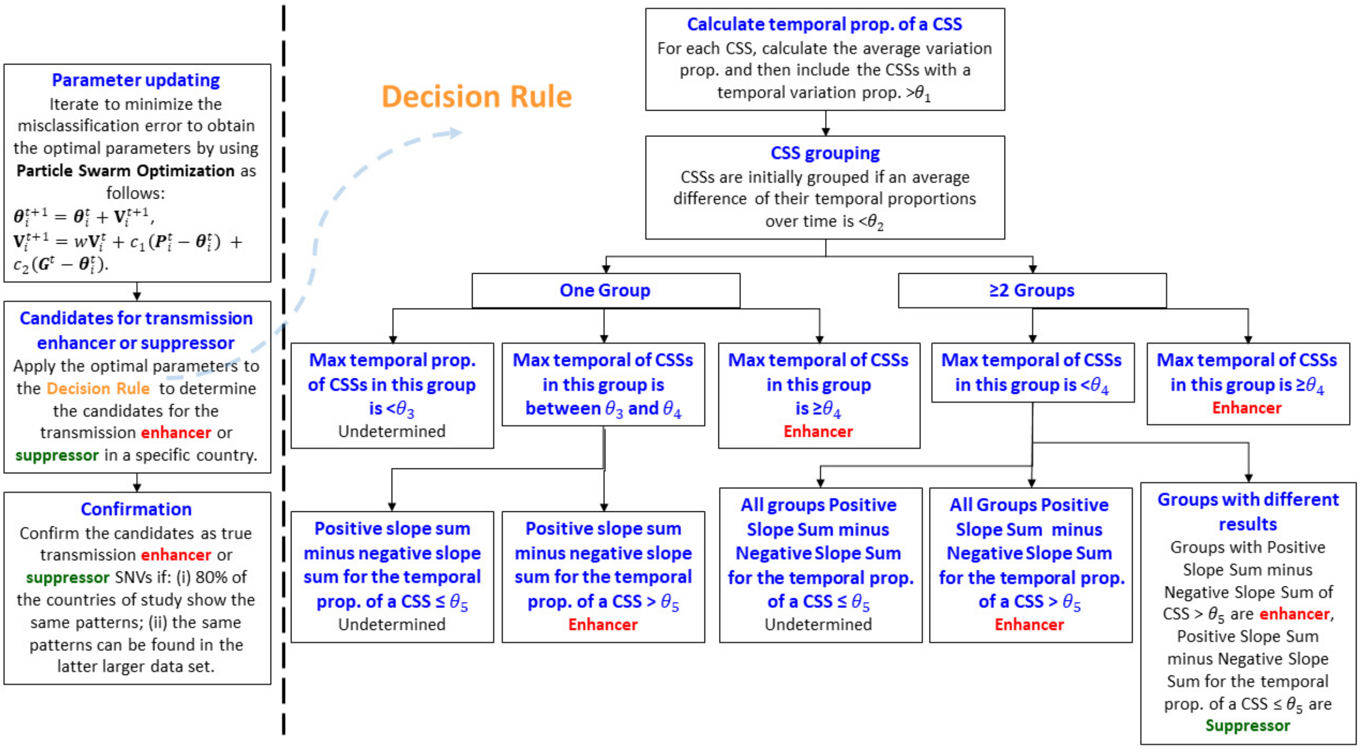
Parameter updating
Iterate to minimize the misclassification error to obtain the optimal parameters by using Particle Swarm Optimization as follows:
$\boldsymbol{\theta}_i^{t+1} = \boldsymbol{\theta}_i^t + \mathbf{V}_i^{t+1}$,
$\mathbf{V}_i^{t+1} = w\mathbf{V}_i^t + c_1(\boldsymbol{P}_i^t - \boldsymbol{\theta}_i^t) + c_2(\boldsymbol{G}^t - \boldsymbol{\theta}_i^t)$.
Candidates for transmission enhancer or suppressor
Apply the optimal parameters to the Decision Rule to determine the candidates for the transmission enhancer or suppressor in a specific country.
Confirmation
Confirm the candidates as true transmission enhancer or suppressor SNVs if: (i) 80% of the countries of study show the same patterns; (ii) the same patterns can be found in the latter larger data set.
Decision Rule
Calculate temporal prop. of a CSS
For each CSS, calculate the average variation prop. and then include the CSSs with a temporal variation prop. >$\theta_1$
CSS grouping
CSSs are initially grouped if an average difference of their temporal proportions over time is <$\theta_2$
One Group
≥2 Groups
Max temporal prop. of CSSs in this group is <$\theta_3$
Undetermined
Max temporal of CSSs in this group is between $\theta_3$ and $\theta_4$
Max temporal of CSSs in this group is ≥$\theta_4$
Enhancer
Max temporal of CSSs in this group is <$\theta_4$
Max temporal of CSSs in this group is ≥$\theta_4$
Enhancer
Positive slope sum minus negative slope sum for the temporal prop. of a CSS ≤ $\theta_5$
Undetermined
Positive slope sum minus negative slope sum for the temporal prop. of a CSS > $\theta_5$
Enhancer
All groups Positive Slope Sum minus Negative Slope Sum for the temporal prop. of a CSS ≤ $\theta_5$
Undetermined
All Groups Positive Slope Sum minus Negative Slope Sum for the temporal prop. of a CSS > $\theta_5$
Enhancer
Groups with different results
Groups with Positive Slope Sum minus Negative Slope Sum of CSS > $\theta_5$ are enhancer, Positive Slope Sum minus Negative Slope Sum for the temporal prop. of a CSS ≤ $\theta_5$ are Suppressor

**Figure S5. Flow charts of the proposed correlated SNV sets (CSSs) analysis and auto-detection algorithm for transmission enhancer SNVs and transmission suppressor SNVs. (A) The proposed correlated SNV sets (CSSs) analysis.** We developed a CSSs analysis pipeline. The analysis was tested based on the genome sequences of more than 2 million SARS-CoV-2 viral sequences collected before 23 Jun, 2021 (n = 2,214K) from the GISAID (https://www.gisaid.org/) and further applied to several larger datasets subsequently. After data quality control, 2,119,724 strains remained. An exponential weighted moving average (EWMA) control chart was employed to detect CSSs. The method tests the null hypothesis $\mathbf{H_0}$ that the expected temporal trajectory of a CSS is smaller than and equal to a prespecified low bound $\boldsymbol{\mu_0}$. If the null hypothesis is rejected statistically, then the SNV set is identified as an emerging CSS (i.e., eCSS, for short, we call it as a CSS). The sensitivity varies from 99.66% to 100%, and specificity is 100% for a prespecified low bound of the temporal trajectory of a CSS $\boldsymbol{\mu_0}$ varied from 0.01 to 0.1 (the CSSs with a temporal trajectory lower than the low bounds are too rare and removed from the analysis). Both of the sensitivity and specificity are high across different thresholds for $\boldsymbol{\mu_0}$, demonstrating its high robustness of the proposed analysis. Note that, in order to reduce the influence of sequencing errors, very rare CSSs (less than 20 strains) are ignored. Conceptually, CSS is constructed based on allelic association of SNVs (i.e., co-occurrence of mutations). Allelic association has been observed in the real SARS-CoV-2 variants such as variants Alpha (aka B.1.1.7 and 501Y.v1), Beta (aka B.1.351 and 501Y.v2), Gamma (aka P.1 and 501Y.v3), Delta (aka B.1.617.2), and Omicron (aka B.1.1.529). The CSSs analysis provides a computationally efficient approach to tracking the viral subtypes and classification, assessing risks and surveillance, and understanding their evolutionary processes. In addition, our results show that our detection for CSSs has a high sensitivity, specificity, and robustness. **(B) The proposed auto-detection algorithm for identifying the CSSs with transmission enhancer SNVs and transmission suppressor SNVs.** We developed an auto-detection algorithm for identifying transmission enhancers SNVs and transmission suppressor SNVs. In a preliminary data set, CSSs were labelled as an enhancer, suppressor, or undetermined based on a heuristic discussion for the pattern of the temporal trajectory of CSSs in a multidisciplinary expert team. Iterations was performed to optimize the parameter vector $\boldsymbol{\theta} = (\theta_1, \theta_2, \theta_3, \theta_4, \theta_5)$ with the minimum misclassification error by using Particle Swarm Optimization (Kennedy and Eberhart, 1995). The optimal parameter vectors were applied to the proposed decision tree algorithm in the right-hand-side in this subfigure. According to the decision rule, CSSs were assigned into enhancer, suppressor, or undetermined. Finally, the candidate enhancers and suppressors should be further confirmed if 80% of the studied countries showed the same pattern of temporal proportion of a CSS and the results can be further confirmed in the later larger dataset.

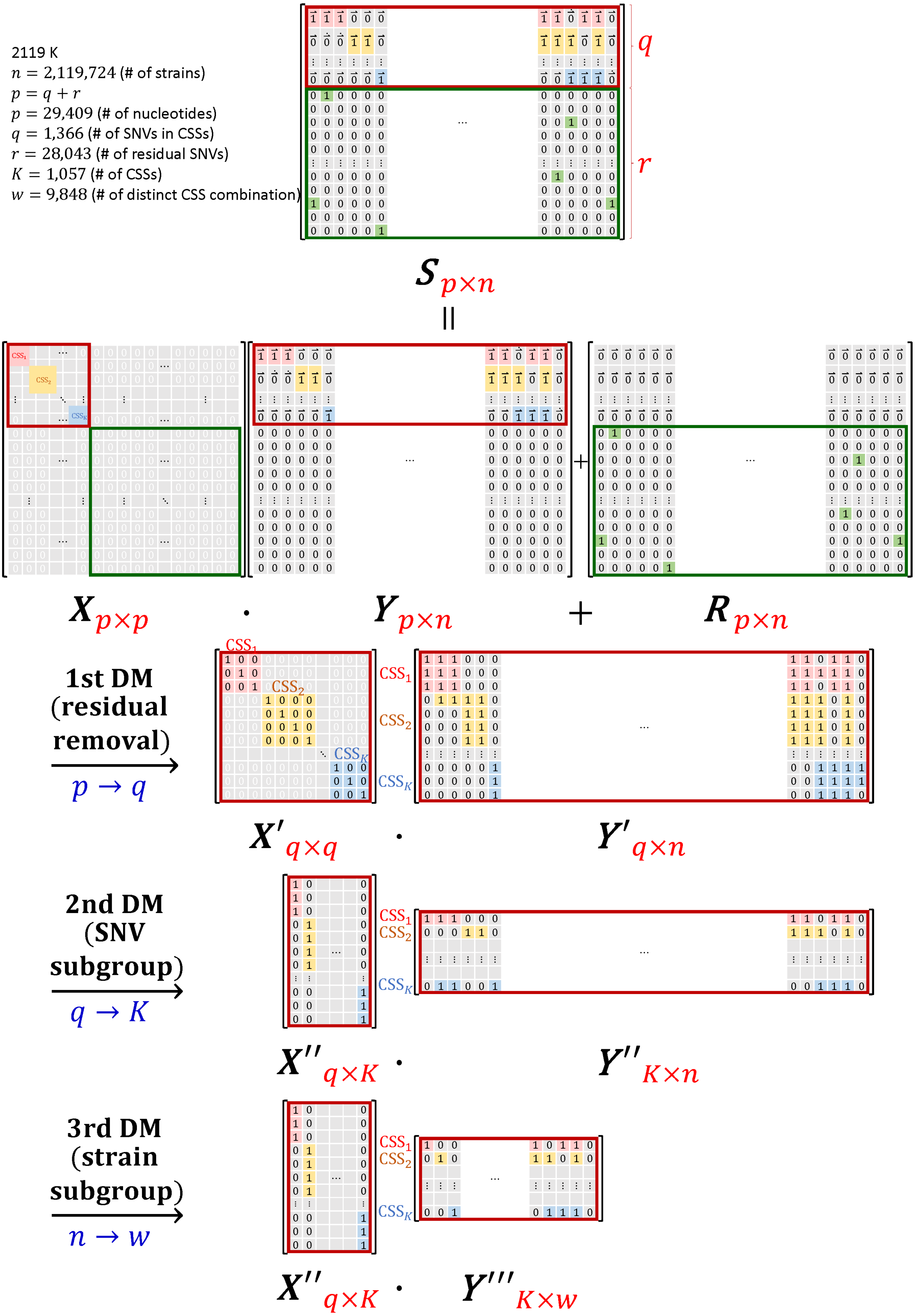

2119 K
$n$ = 2,119,724 (# of strains)
$p = q + r$
$p$ = 29,409 (# of nucleotides)
$q$ = 1,366 (# of SNVs in CSSs)
$r$ = 28,043 (# of residual SNVs)
$K$ = 1,057 (# of CSSs)
$w$ = 9,848 (# of distinct CSS combination)
$q$
$r$
$S_{p\times n}$
||
$X_{p\times p}$ · $Y_{p\times n}$ + $R_{p\times n}$
1st DM (residual removal)
$p \rightarrow q$
$X'_{q\times q}$ · $Y'_{q\times n}$
2nd DM (SNV subgroup)
$q \rightarrow K$
$X''_{q\times K}$ · $Y''_{K\times n}$
3rd DM (strain subgroup)
$n \rightarrow w$
$X''_{q\times K}$ · $Y'''_{K\times w}$
$CSS_1$
$CSS_2$
$CSS_K$

**Figure S6. Matrix representation of dimension reduction by using correlated SNV sets (CSSs).** This diagram demonstrates a three-step dimension reduction (DM). Let $p$ indicates the number of nucleotides in viral genome after removing 3' leader and 5' end sequence regions ($p = 29{,}409$); $q$ indicates the number of SNVs in CSSs ($q = 1{,}366$), where $q = \sum_{k=1}^{K} c_k$ and $c_k$ is the # of SNVs in the $k$-th CSS and $K$ is the number of CSSs ($K = 1{,}057$); $r$ indicates the number of SNVs not in any of CSSs ($r = 28{,}043$), where $p = q + r$; $n$ indicates the total number of strains in this data set ($n = 2{,}119{,}724$). $\boldsymbol{S}$ indicates the matrix of genome sequences with $p$ rows and $n$ columns. Rows have been rearranged by the order of CSSs. The upper part of $\boldsymbol{S}$ (i.e., the red-frame square with $q$ rows and $n$ columns) is constructed by the SNVs in CSSs. For each strain (column), the genomic regions in a CSS are assigned as a vector of 1's ($\vec{1}$), 0's ($\vec{0}$), or $\dot{\vec{0}}$ if this CSS is completely found, completely unfound, or partially unfound in this viral strain, respectively. $\boldsymbol{S}$ is partitioned as $\boldsymbol{S} = \boldsymbol{X} \cdot \boldsymbol{Y} + \boldsymbol{R}$, where $\boldsymbol{R}$ indicates the residual matrix. Here $\boldsymbol{X} \cdot \boldsymbol{Y}$ and $\boldsymbol{R}$ indicate the component that can and cannot be represented by CSSs, respectively. In $\boldsymbol{X}$ and $\boldsymbol{Y}$, for the CSSs part (i.e., the red-frame square), the $k$-th diagonal block is an identity matrix designed for the $k$-th CSS in $\boldsymbol{X}$ and the $i$-th column indicates the indictor vector of CSSs for the $i$-th strain in $\boldsymbol{Y}$. Others are zero-valued. In $\boldsymbol{R}$, for the residual part (i.e., the green-frame square), the $i$-th column indicates the residual vector for the $i$-th strain. Residual, the unrepresented SNVs, for the $i$-th strain is the cardinal number of the $i$-th column vector (i.e., the number of 1's in a column). Others are zero-valued. In the first DM, $\boldsymbol{X}'$ and $\boldsymbol{Y}'$ provide a SNVs dimension reduction for $\boldsymbol{X}$ and $\boldsymbol{Y}$ for the viral sequences after ignoring the residual SNVs. The number of SNVs is reduced from $p = 29{,}409$ to $q = 1{,}366$. In the second DM, $\boldsymbol{X}''$ and $\boldsymbol{Y}''$ provide a dimension reduction for $\boldsymbol{X}'$ and $\boldsymbol{Y}'$ that the signature SNVs are grouped into CSSs. That is, $q = 1{,}366$ is reduced to $K = 1{,}057$. In the last DM, $\boldsymbol{X}''$ and $\boldsymbol{Y}'''$ provide a dimension reduction for $\boldsymbol{X}''$ and $\boldsymbol{Y}''$ that the large number of strains are grouped into the subtypes according to CSSs. That is, $n = 2{,}119{,}724$ is reduced to $w = 1{,}057$. This analysis was conducted based on the genome sequences of more than 2 million of SARS-CoV-2 viral strains collected before 23 Jun, 2021 ($n = 2{,}119{,}724$) from the GISAID (https://www.gisaid.org/). The majority of the viral strains (>98.28%) can be characterized by a single CSS or a few numbers of the 1,057 CSSs consisting of 1,384 distinct signature SNVs (from 1,432 signature SNVs that some SNVs have various alternative alleles). We find that 1,053 of 1,057 CSSs can characterize >99.9% of the current dominant strain type (Type VI). Type VI became the dominant strain (2,000,622 strains and 94.38%) since March, 2020. Among 2,000,622 Type VI strains, 98.79% of the strains can be further represented by 171 primary CSSs which are constructed by 557 SNVs. The remaining 882 secondary CSSs can be represented by the 171 primary CSSs. The proportion explained by CSSs is further increased to 99.95% if additional 42 SNVs with a variation frequency of >0.01 are accounted. The CSSs can cover the well-recognized variants such as B.1.1.7 (aka variant 501Y.v1 or Alpha), B.1.351 (aka variant 501Y.v2 or Beta), P.1 (aka variant 501Y.v3 or Gamma), and B.1.617.2 (aka variant Delta) and exhibit the subtypes and heterogeneity of these variants.

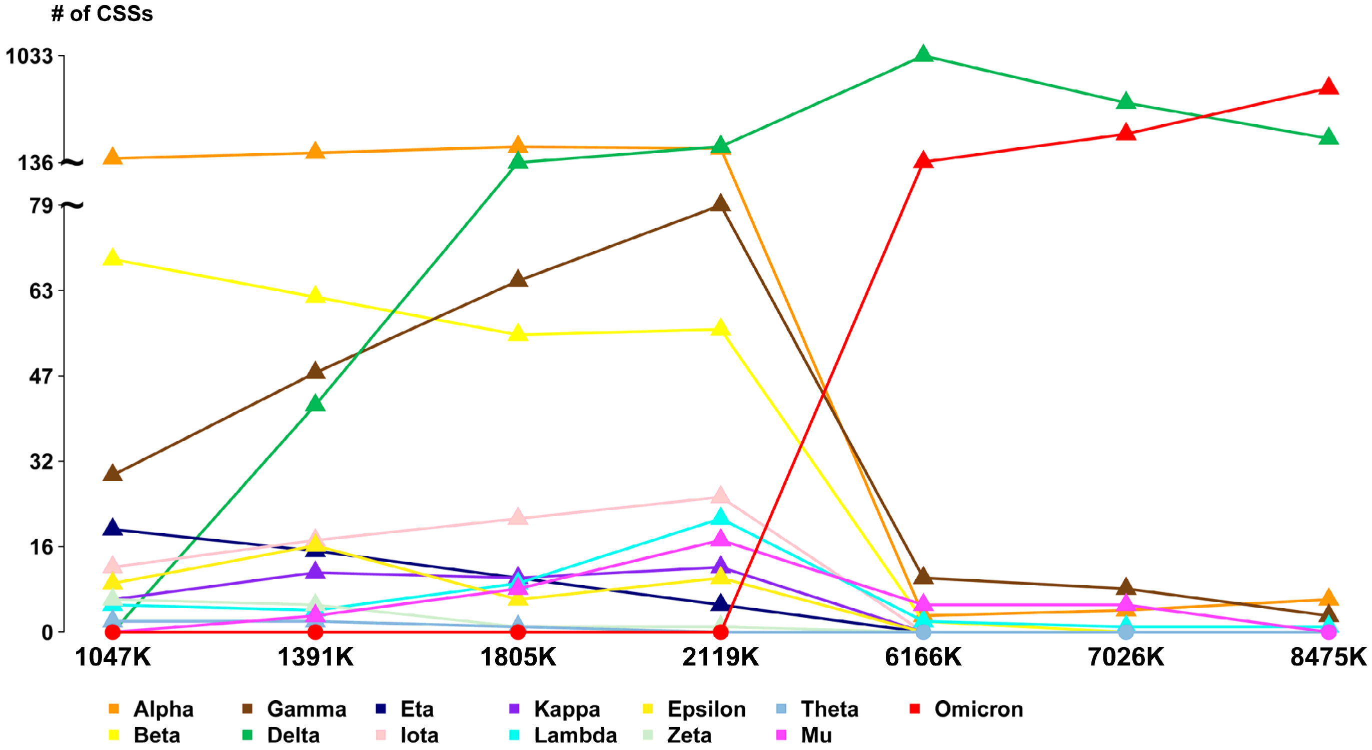


**Figure S7. Number of correlated SNV sets (CSSs) for the well-recognized variants in seven datasets.** This figure shows the temporal change of the numbers of CSSs in the seven datasets that the data were collected from Dec 2019 to 7 Apr, 5 May, 26 May, 23 Jun, 2021, 15 Dec for year 2021 and 1 Jan and 23 Feb for year 2022 with a sample size of n = 1,047K, 1,391K, 1,805K, 2,119K, 6,166K, 7,026K, and 8,475K, respectively. As of 23 June, 2021 (n = 2,119K), compared to the Alpha strain that the increase in the number of CSSs attenuates, the Delta variant exhibits a steep increase, indicating of a large heterogeneity and continued evolution of the Delta variant. As of December, 2021 (n = 6,166K), compared to the Alpha and Delta strains, the Omicron variant exhibits a much steeper increase in number of CSSs, illustrating a rapid transmission of the Omicron variant.

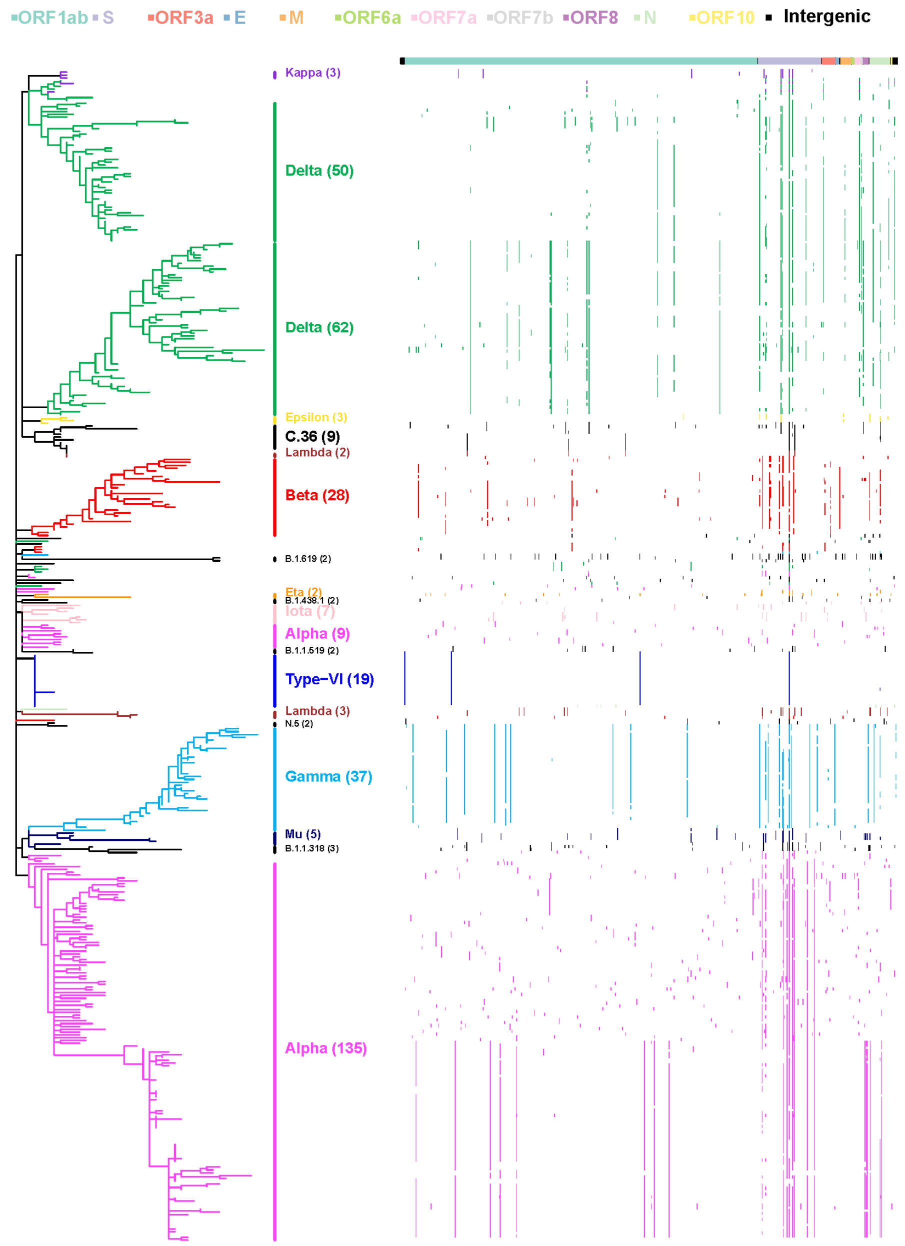


**Figure S8. Phylogenetic dendrogram of correlated SNV sets (CSSs) (n = 2,119K).** This sub-figure contains three components: (i) Maximum parsimony (MP) dendrogram for the CSSs. CSSs are sorted using terminal nodes order from the MP dendrogram. (ii) PANGO nomenclature. A CSS may contain a number of strains with various PANGO nomenclature from the meta information provided by GISAID. The PANGO nomenclature for the majority of the viral strains in a CSS is assigned. (iii) single nucleotide variation (SNV) matrix map for 417 CSSs with 669 SNV sites (For a clear view, only the CSSs with a strain size of ≥200 are shown). SNVs are sorted according to their physical positions in the viral genome, including 265 sites in the 5' leader and 229 sites in the 3' terminal sequences. Protein names are listed in the top panel of the SNV matrix map. If signature SNVs for CSSs have a nucleotide change different from the Wuhan-Hu-1 strain, dots are displayed in the SNV matrix map. The color of signature SNVs in CSSs is assigned according to the color of the CSS group in the MP dendrogram. Allelic association of the signature SNVs can be observed through a co-occurrence of signature SNVs in a certain proportion of viral strains or CSSs. Phylogenetic analysis based on maximum parsimony (MP) was conducted by using MEGA X (Kumar et al., Mol Biol Evol 35, 2018). Subtree-Pruning-Regrafting algorithm (Nei and Kumar, Oxford university press, 2000) was employed for a tree topology search heuristic.

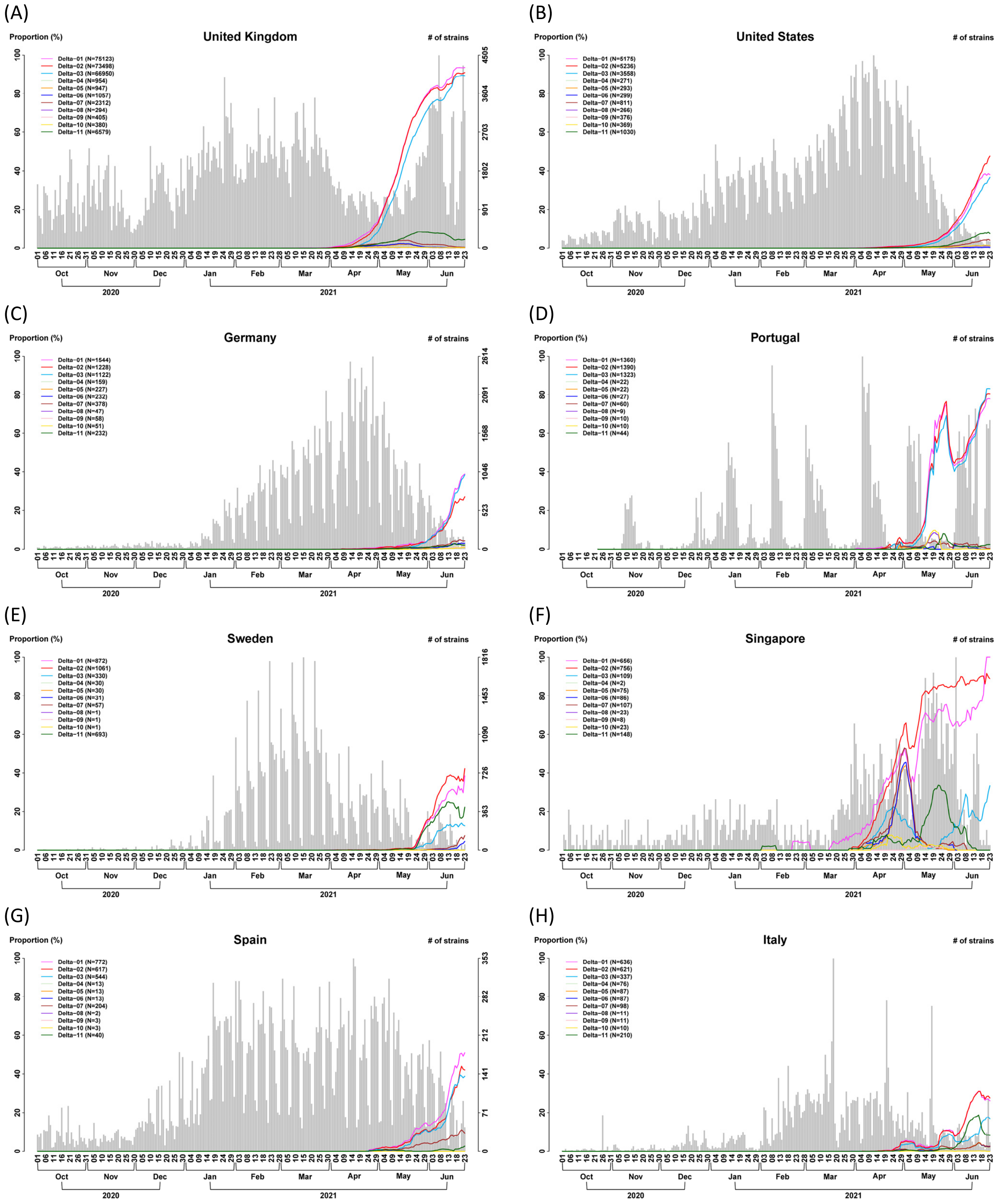
(A)
United Kingdom
Proportion (%)
### of strains
Delta−01 (N=75123)
Delta−02 (N=73498)
Delta−03 (N=66950)
Delta−04 (N=954)
Delta−05 (N=947)
Delta−06 (N=1057)
Delta−07 (N=2312)
Delta−08 (N=294)
Delta−09 (N=405)
Delta−10 (N=380)
Delta−11 (N=6579)
(B)
United States
Delta−01 (N=5175)
Delta−02 (N=5236)
Delta−03 (N=3558)
Delta−04 (N=271)
Delta−05 (N=293)
Delta−06 (N=299)
Delta−07 (N=811)
Delta−08 (N=266)
Delta−09 (N=376)
Delta−10 (N=369)
Delta−11 (N=1030)
(C)
Germany
Delta−01 (N=1544)
Delta−02 (N=1228)
Delta−03 (N=1122)
Delta−04 (N=159)
Delta−05 (N=227)
Delta−06 (N=232)
Delta−07 (N=378)
Delta−08 (N=47)
Delta−09 (N=58)
Delta−10 (N=51)
Delta−11 (N=232)
(D)
Portugal
Delta−01 (N=1360)
Delta−02 (N=1390)
Delta−03 (N=1323)
Delta−04 (N=22)
Delta−05 (N=22)
Delta−06 (N=27)
Delta−07 (N=60)
Delta−08 (N=9)
Delta−09 (N=10)
Delta−10 (N=10)
Delta−11 (N=44)
(E)
Sweden
Delta−01 (N=872)
Delta−02 (N=1061)
Delta−03 (N=330)
Delta−04 (N=30)
Delta−05 (N=30)
Delta−06 (N=31)
Delta−07 (N=57)
Delta−08 (N=1)
Delta−09 (N=1)
Delta−10 (N=1)
Delta−11 (N=693)
(F)
Singapore
Delta−01 (N=656)
Delta−02 (N=756)
Delta−03 (N=109)
Delta−04 (N=2)
Delta−05 (N=75)
Delta−06 (N=86)
Delta−07 (N=107)
Delta−08 (N=23)
Delta−09 (N=8)
Delta−10 (N=23)
Delta−11 (N=148)
(G)
Spain
Delta−01 (N=772)
Delta−02 (N=617)
Delta−03 (N=544)
Delta−04 (N=13)
Delta−05 (N=13)
Delta−06 (N=13)
Delta−07 (N=204)
Delta−08 (N=2)
Delta−09 (N=3)
Delta−10 (N=3)
Delta−11 (N=40)
(H)
Italy
Delta−01 (N=636)
Delta−02 (N=621)
Delta−03 (N=337)
Delta−04 (N=76)
Delta−05 (N=87)
Delta−06 (N=87)
Delta−07 (N=98)
Delta−08 (N=11)
Delta−09 (N=11)
Delta−10 (N=10)
Delta−11 (N=210)
Oct
Nov
Dec
Jan
Feb
Mar
Apr
May
Jun
2020
2021

(I)

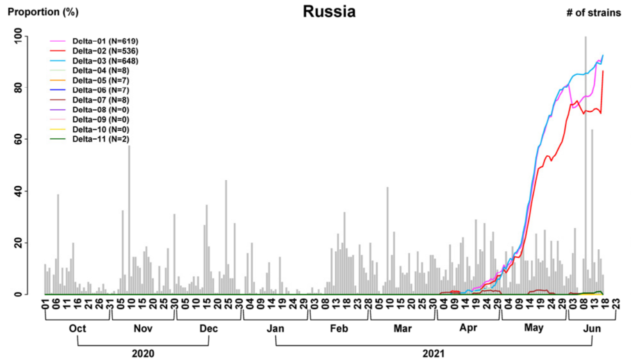
Russia
Proportion (%)
### of strains
Delta−01 (N=619)
Delta−02 (N=536)
Delta−03 (N=648)
Delta−04 (N=8)
Delta−05 (N=7)
Delta−06 (N=7)
Delta−07 (N=8)
Delta−08 (N=0)
Delta−09 (N=0)
Delta−10 (N=0)
Delta−11 (N=2)
144 115 86 58 29 0
Oct Nov Dec Jan Feb Mar Apr May Jun
2020 2021

(J)

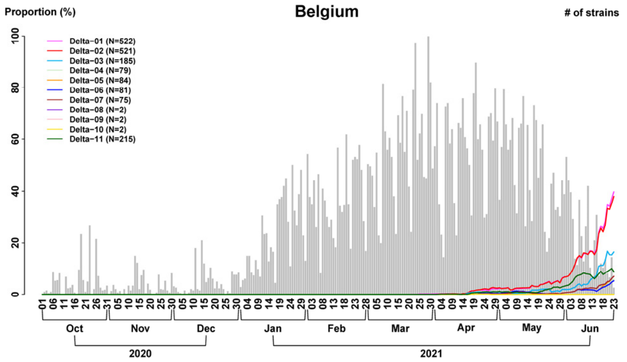
Belgium
Proportion (%)
### of strains
Delta−01 (N=522)
Delta−02 (N=521)
Delta−03 (N=185)
Delta−04 (N=79)
Delta−05 (N=84)
Delta−06 (N=81)
Delta−07 (N=75)
Delta−08 (N=2)
Delta−09 (N=2)
Delta−10 (N=2)
Delta−11 (N=215)
375 300 225 150 75 0
Oct Nov Dec Jan Feb Mar Apr May Jun
2020 2021

(K)

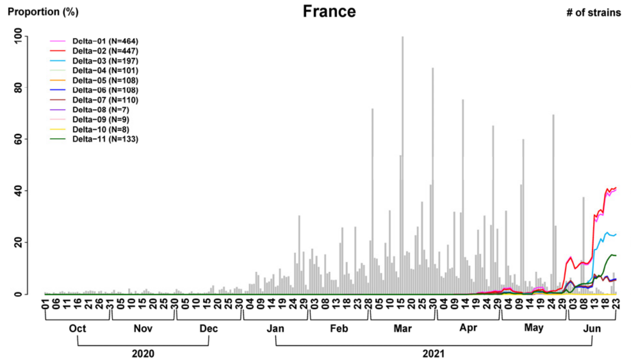
France
Proportion (%)
### of strains
Delta−01 (N=464)
Delta−02 (N=447)
Delta−03 (N=197)
Delta−04 (N=101)
Delta−05 (N=108)
Delta−06 (N=108)
Delta−07 (N=110)
Delta−08 (N=7)
Delta−09 (N=9)
Delta−10 (N=8)
Delta−11 (N=133)
1818 1454 1091 727 364 0
Oct Nov Dec Jan Feb Mar Apr May Jun
2020 2021

(L)

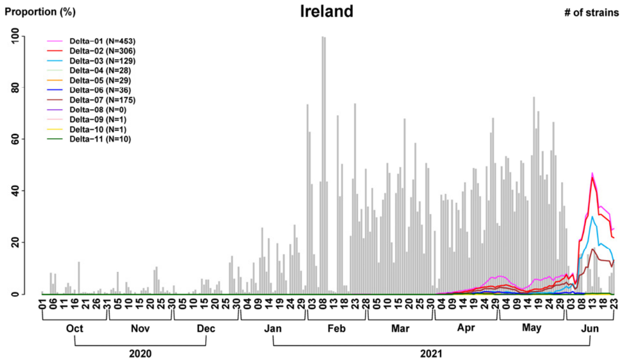
Ireland
Proportion (%)
### of strains
Delta−01 (N=453)
Delta−02 (N=306)
Delta−03 (N=129)
Delta−04 (N=28)
Delta−05 (N=29)
Delta−06 (N=36)
Delta−07 (N=175)
Delta−08 (N=0)
Delta−09 (N=1)
Delta−10 (N=1)
Delta−11 (N=10)
319 255 191 128 64 0
Oct Nov Dec Jan Feb Mar Apr May Jun
2020 2021

(M)

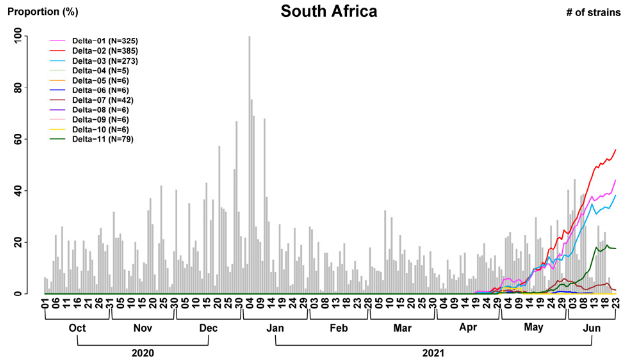
South Africa
Proportion (%)
### of strains
Delta−01 (N=325)
Delta−02 (N=385)
Delta−03 (N=273)
Delta−04 (N=5)
Delta−05 (N=6)
Delta−06 (N=6)
Delta−07 (N=42)
Delta−08 (N=6)
Delta−09 (N=6)
Delta−10 (N=6)
Delta−11 (N=79)
188 150 113 75 38 0
Oct Nov Dec Jan Feb Mar Apr May Jun
2020 2021

(N)

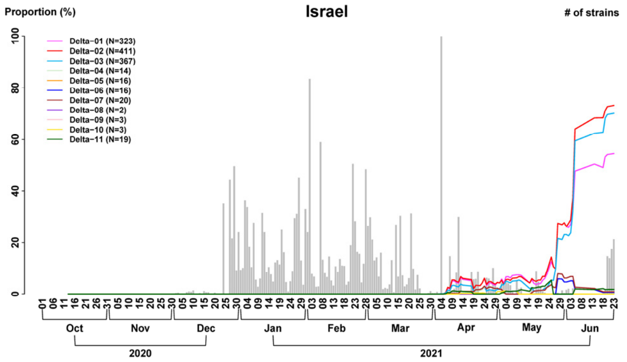
Israel
Proportion (%)
### of strains
Delta−01 (N=323)
Delta−02 (N=411)
Delta−03 (N=367)
Delta−04 (N=14)
Delta−05 (N=16)
Delta−06 (N=16)
Delta−07 (N=20)
Delta−08 (N=2)
Delta−09 (N=3)
Delta−10 (N=3)
Delta−11 (N=19)
660 528 396 264 132 0
Oct Nov Dec Jan Feb Mar Apr May Jun
2020 2021

(O)

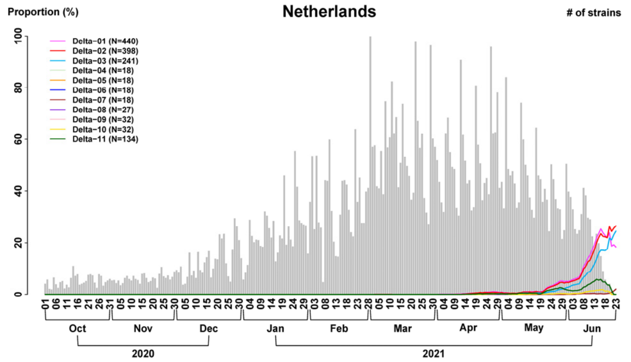
Netherlands
Proportion (%)
### of strains
Delta−01 (N=440)
Delta−02 (N=398)
Delta−03 (N=241)
Delta−04 (N=18)
Delta−05 (N=18)
Delta−06 (N=18)
Delta−07 (N=18)
Delta−08 (N=27)
Delta−09 (N=32)
Delta−10 (N=32)
Delta−11 (N=134)
497 398 298 199 99 0
Oct Nov Dec Jan Feb Mar Apr May Jun
2020 2021

(P)

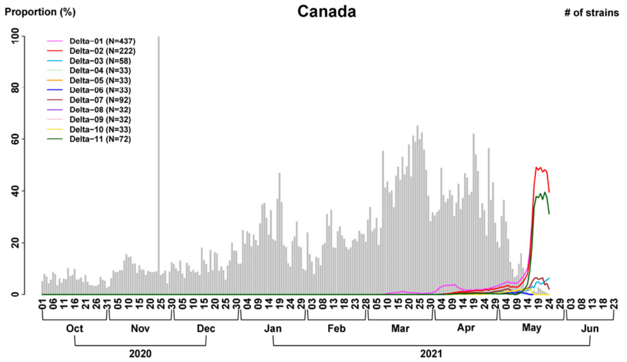
Canada
Proportion (%)
### of strains
Delta−01 (N=437)
Delta−02 (N=222)
Delta−03 (N=58)
Delta−04 (N=33)
Delta−05 (N=33)
Delta−06 (N=33)
Delta−07 (N=92)
Delta−08 (N=32)
Delta−09 (N=32)
Delta−10 (N=33)
Delta−11 (N=72)
870 696 522 348 174 0
Oct Nov Dec Jan Feb Mar Apr May Jun
2020 2021

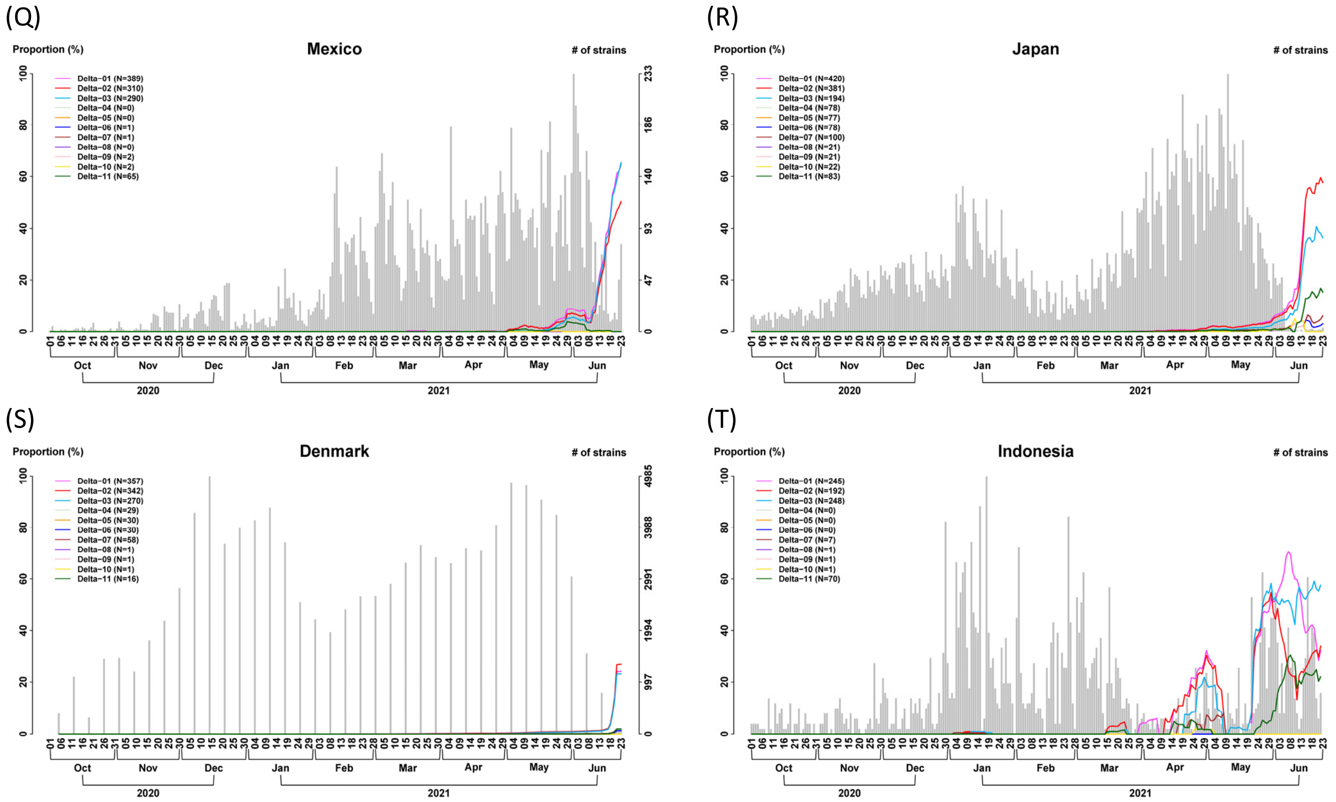


**Figure S9. Differential temporal trajectories for the eleven Delta subtypes with transmission enhancer and/or suppressor SNVs in many countries (n = 2,119K).** The first three increasing subtypes have rising temporal trajectories (Delta-01, Delta-02, and Delta-03). The remaining eight subtypes (Delta-04, Delta-05, Delta-06, Delta-07, Delta-08, Delta-09, Delta-10, and Delta-11) have a lower temporal trajectory. Remarkably, the same pattern of the differential temporal trajectories for the eleven subtypes was found not only in India but also in many other countries consistently. (**A**) The United Kingdom; (**B**) The United States; (**C**) Germany; (**D**) Portugal; (**E**) Sweden; (**F**) Singapore; (**G**) Spain; (**H**) Italy; (**I**) Russia; (**J**) Belgium; (**K**) France; (**L**) Ireland; (**M**) South Africa; (**N**) Israel; (**O**) The Netherlands; (**P**) Canada; (**Q**) Mexico; (**R**) Japan; (**S**) Denmark; (**T**) Indonesia.

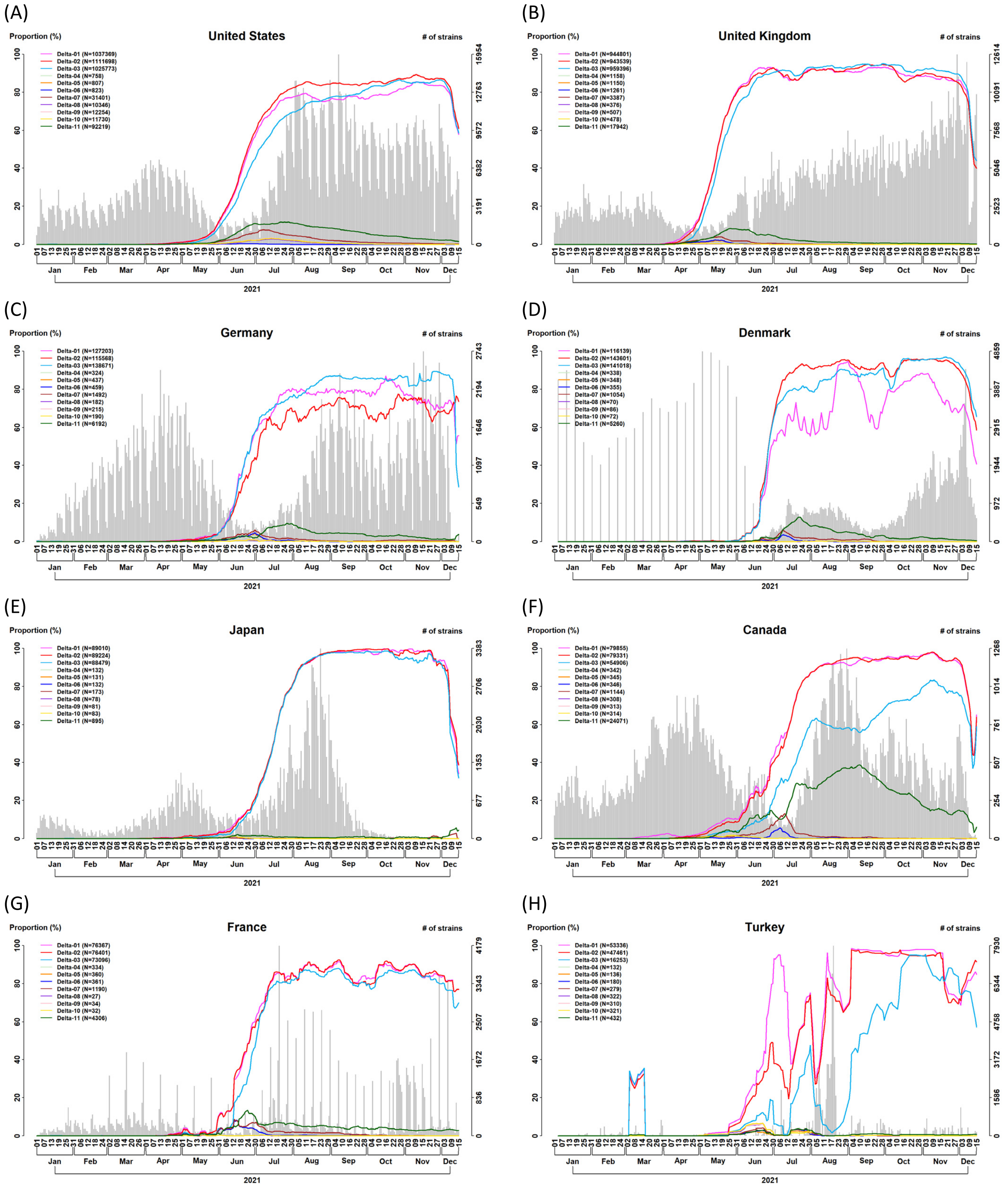
(A)
United States
Proportion (%)
### of strains
Delta-01 (N=1037369)
Delta-02 (N=1111698)
Delta-03 (N=1025773)
Delta-04 (N=758)
Delta-05 (N=807)
Delta-06 (N=823)
Delta-07 (N=31401)
Delta-08 (N=10346)
Delta-09 (N=12254)
Delta-10 (N=11730)
Delta-11 (N=92219)
100 80 60 40 20 0
15954 12763 9572 6382 3191 0
Jan Feb Mar Apr May Jun Jul Aug Sep Oct Nov Dec
2021
(B)
United Kingdom
Proportion (%)
### of strains
Delta-01 (N=944801)
Delta-02 (N=943539)
Delta-03 (N=959396)
Delta-04 (N=1158)
Delta-05 (N=1150)
Delta-06 (N=1261)
Delta-07 (N=3387)
Delta-08 (N=376)
Delta-09 (N=507)
Delta-10 (N=478)
Delta-11 (N=17942)
12614 10091 7568 5046 2523 0
2021
(C)
Germany
Proportion (%)
### of strains
Delta-01 (N=127203)
Delta-02 (N=115568)
Delta-03 (N=138671)
Delta-04 (N=324)
Delta-05 (N=437)
Delta-06 (N=459)
Delta-07 (N=1492)
Delta-08 (N=182)
Delta-09 (N=215)
Delta-10 (N=190)
Delta-11 (N=6192)
2743 2194 1646 1097 549 0
2021
(D)
Denmark
Proportion (%)
### of strains
Delta-01 (N=116139)
Delta-02 (N=143601)
Delta-03 (N=141018)
Delta-04 (N=338)
Delta-05 (N=348)
Delta-06 (N=355)
Delta-07 (N=1054)
Delta-08 (N=70)
Delta-09 (N=86)
Delta-10 (N=72)
Delta-11 (N=5260)
4859 3887 2915 1944 972 0
2021
(E)
Japan
Proportion (%)
### of strains
Delta-01 (N=89010)
Delta-02 (N=89224)
Delta-03 (N=88479)
Delta-04 (N=132)
Delta-05 (N=131)
Delta-06 (N=132)
Delta-07 (N=173)
Delta-08 (N=78)
Delta-09 (N=81)
Delta-10 (N=83)
Delta-11 (N=895)
3383 2706 2030 1353 677 0
2021
(F)
Canada
Proportion (%)
### of strains
Delta-01 (N=79855)
Delta-02 (N=79331)
Delta-03 (N=54906)
Delta-04 (N=342)
Delta-05 (N=345)
Delta-06 (N=346)
Delta-07 (N=1144)
Delta-08 (N=308)
Delta-09 (N=313)
Delta-10 (N=314)
Delta-11 (N=24071)
1268 1014 761 507 254 0
2021
(G)
France
Proportion (%)
### of strains
Delta-01 (N=76367)
Delta-02 (N=76401)
Delta-03 (N=73096)
Delta-04 (N=334)
Delta-05 (N=360)
Delta-06 (N=361)
Delta-07 (N=1190)
Delta-08 (N=27)
Delta-09 (N=34)
Delta-10 (N=32)
Delta-11 (N=4306)
4179 3343 2507 1672 836 0
2021
(H)
Turkey
Proportion (%)
### of strains
Delta-01 (N=53336)
Delta-02 (N=47461)
Delta-03 (N=16253)
Delta-04 (N=132)
Delta-05 (N=136)
Delta-06 (N=180)
Delta-07 (N=279)
Delta-08 (N=322)
Delta-09 (N=310)
Delta-10 (N=321)
Delta-11 (N=432)
7930 6344 4758 3172 1586 0
2021

(I)

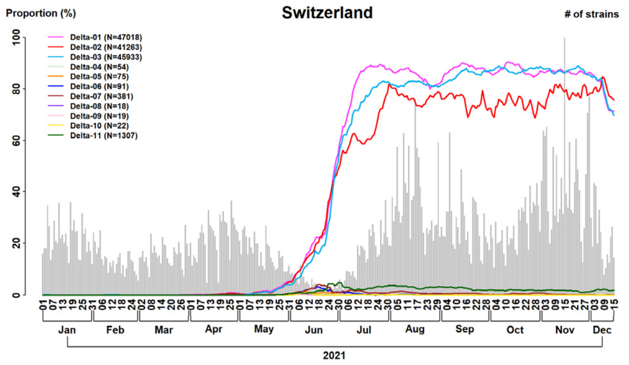
Switzerland
Proportion (%)
### of strains
Delta-01 (N=47018)
Delta-02 (N=41263)
Delta-03 (N=45933)
Delta-04 (N=54)
Delta-05 (N=75)
Delta-06 (N=91)
Delta-07 (N=381)
Delta-08 (N=18)
Delta-09 (N=19)
Delta-10 (N=22)
Delta-11 (N=1307)
1095
876
657
438
219
Jan Feb Mar Apr May Jun Jul Aug Sep Oct Nov Dec
2021

(J)

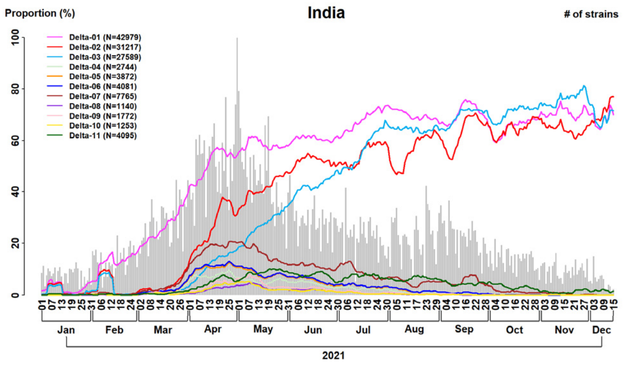
India
Proportion (%)
### of strains
Delta-01 (N=42979)
Delta-02 (N=31217)
Delta-03 (N=27589)
Delta-04 (N=2744)
Delta-05 (N=3872)
Delta-06 (N=4081)
Delta-07 (N=7765)
Delta-08 (N=1140)
Delta-09 (N=1772)
Delta-10 (N=1253)
Delta-11 (N=4095)
971
777
583
388
194
Jan Feb Mar Apr May Jun Jul Aug Sep Oct Nov Dec
2021

(K)

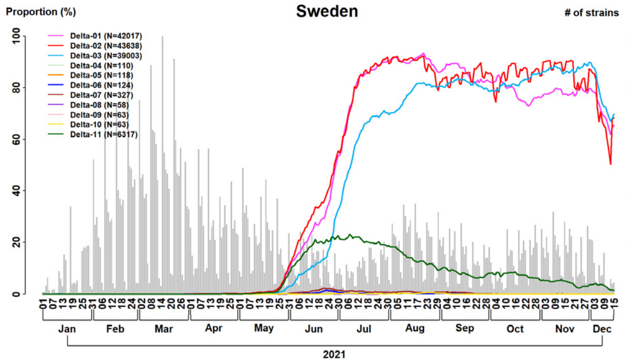
Sweden
Proportion (%)
### of strains
Delta-01 (N=42017)
Delta-02 (N=43638)
Delta-03 (N=39003)
Delta-04 (N=110)
Delta-05 (N=118)
Delta-06 (N=124)
Delta-07 (N=327)
Delta-08 (N=58)
Delta-09 (N=63)
Delta-10 (N=63)
Delta-11 (N=6317)
2111
1689
1267
844
422
Jan Feb Mar Apr May Jun Jul Aug Sep Oct Nov Dec
2021

(L)

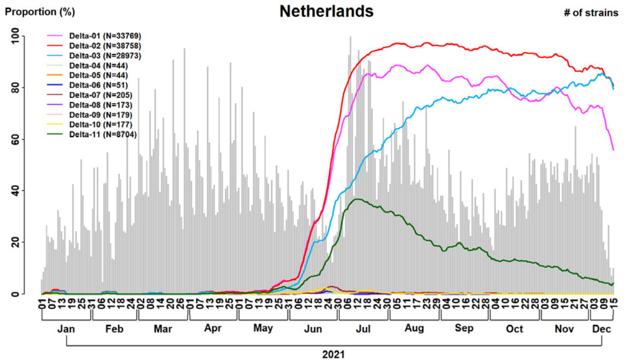
Netherlands
Proportion (%)
### of strains
Delta-01 (N=33769)
Delta-02 (N=38758)
Delta-03 (N=28973)
Delta-04 (N=44)
Delta-05 (N=44)
Delta-06 (N=51)
Delta-07 (N=205)
Delta-08 (N=173)
Delta-09 (N=179)
Delta-10 (N=177)
Delta-11 (N=8704)
540
432
324
216
108
Jan Feb Mar Apr May Jun Jul Aug Sep Oct Nov Dec
2021

(M)

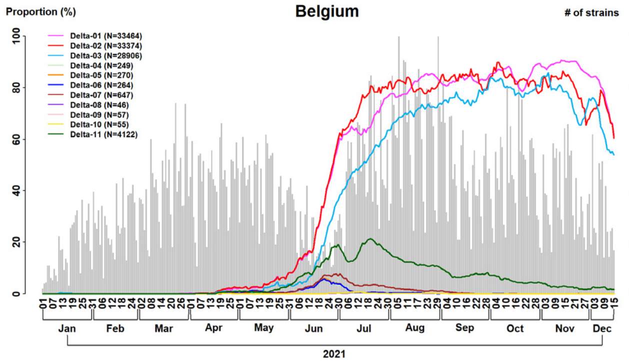
Belgium
Proportion (%)
### of strains
Delta-01 (N=33464)
Delta-02 (N=33374)
Delta-03 (N=28906)
Delta-04 (N=249)
Delta-05 (N=270)
Delta-06 (N=264)
Delta-07 (N=647)
Delta-08 (N=46)
Delta-09 (N=57)
Delta-10 (N=55)
Delta-11 (N=4122)
512
410
307
205
102
Jan Feb Mar Apr May Jun Jul Aug Sep Oct Nov Dec
2021

(N)

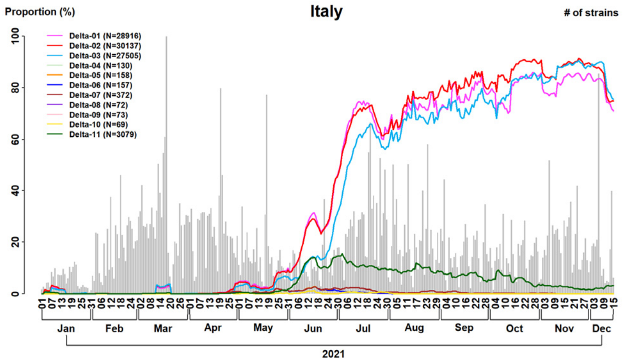
Italy
Proportion (%)
### of strains
Delta-01 (N=28916)
Delta-02 (N=30137)
Delta-03 (N=27505)
Delta-04 (N=130)
Delta-05 (N=158)
Delta-06 (N=157)
Delta-07 (N=372)
Delta-08 (N=72)
Delta-09 (N=73)
Delta-10 (N=69)
Delta-11 (N=3079)
1115
892
669
446
223
Jan Feb Mar Apr May Jun Jul Aug Sep Oct Nov Dec
2021

(O)

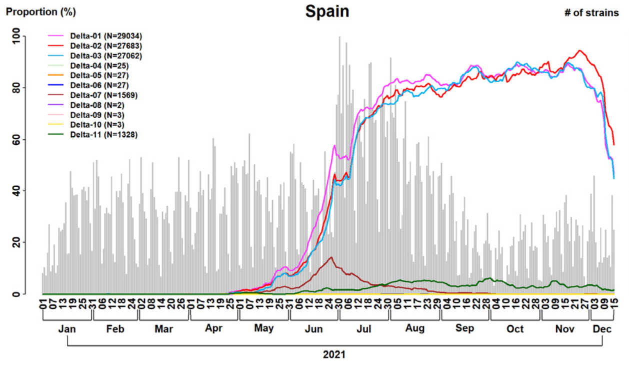
Spain
Proportion (%)
### of strains
Delta-01 (N=29034)
Delta-02 (N=27683)
Delta-03 (N=27062)
Delta-04 (N=25)
Delta-05 (N=27)
Delta-06 (N=27)
Delta-07 (N=1569)
Delta-08 (N=2)
Delta-09 (N=3)
Delta-10 (N=3)
Delta-11 (N=1328)
617
494
370
247
123
Jan Feb Mar Apr May Jun Jul Aug Sep Oct Nov Dec
2021

(P)

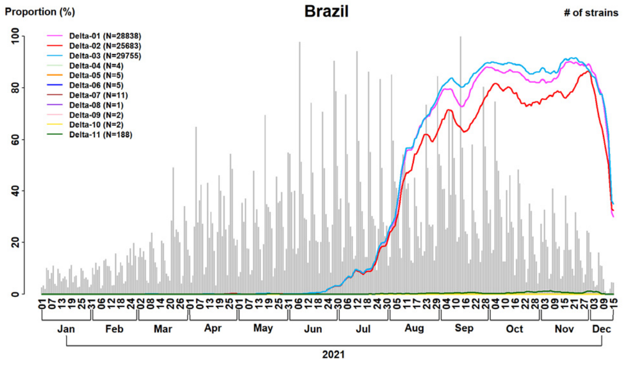
Brazil
Proportion (%)
### of strains
Delta-01 (N=28838)
Delta-02 (N=25683)
Delta-03 (N=29755)
Delta-04 (N=4)
Delta-05 (N=5)
Delta-06 (N=5)
Delta-07 (N=11)
Delta-08 (N=1)
Delta-09 (N=2)
Delta-10 (N=2)
Delta-11 (N=188)
981
785
589
392
196
Jan Feb Mar Apr May Jun Jul Aug Sep Oct Nov Dec
2021

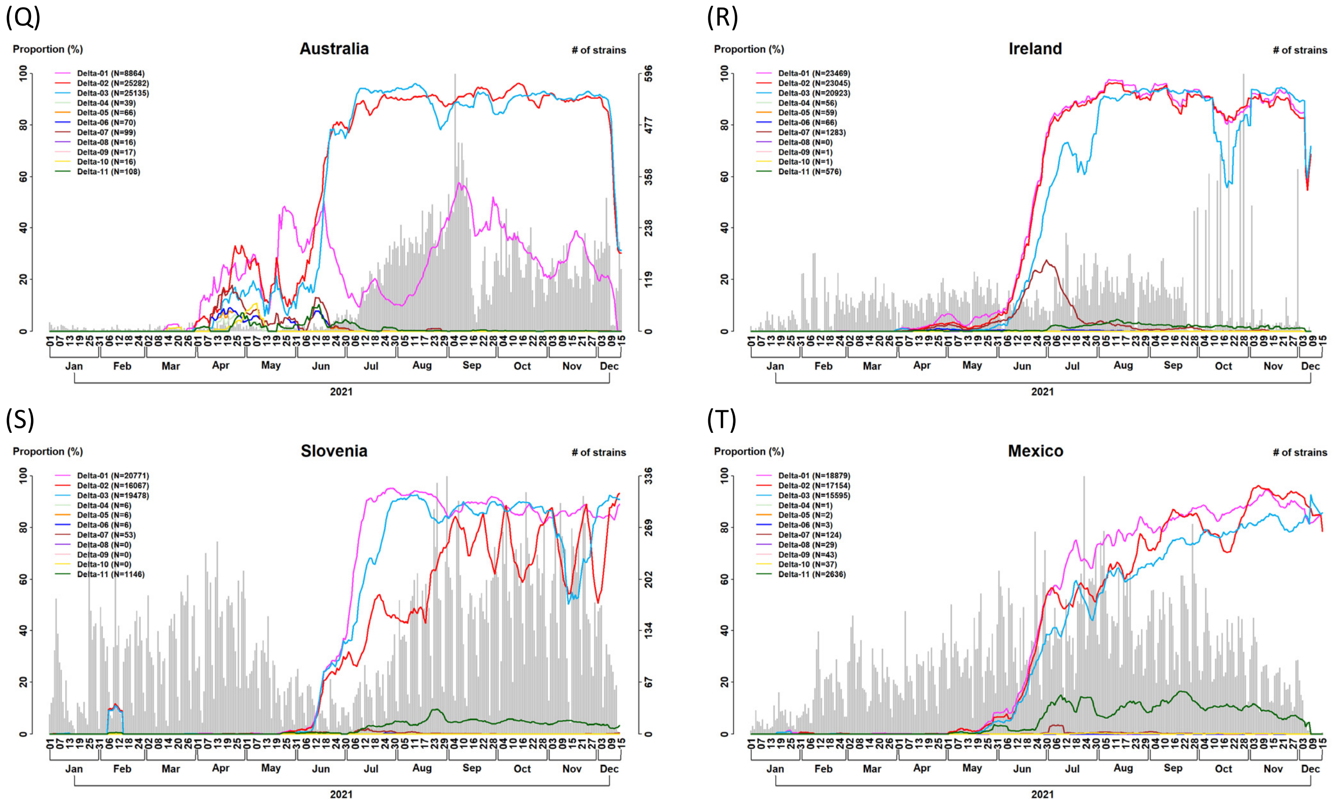


**Figure S10. Differential temporal trajectories for the eleven Delta subtypes with transmission enhancer and/or suppressor SNVs in many countries (n = 6,166K).** The first three increasing subtypes have rising temporal trajectories (Delta-01, Delta-02, and Delta-03). The remaining eight subtypes (Delta-04, Delta-05, Delta-06, Delta-07, Delta-08, Delta-09, Delta-10, and Delta-11) have a lower temporal trajectory. Remarkably, the same pattern of the differential temporal trajectories for the eleven subtypes was found not only in India but also in many other countries consistently. (A) The United States; (B) The United Kingdom; (C) Germany; (D) Denmark; (E) Japan; (F) Canada; (G) France; (H) Turkey; (I) Switzerland; (J) India; (K) Sweden; (L) The Netherlands; (M) Belgium; (N) Italy; (O) Spain; (P) Brazil; (Q) Australia; (R) Ireland; (S) Slovenia; (T) Mexico.

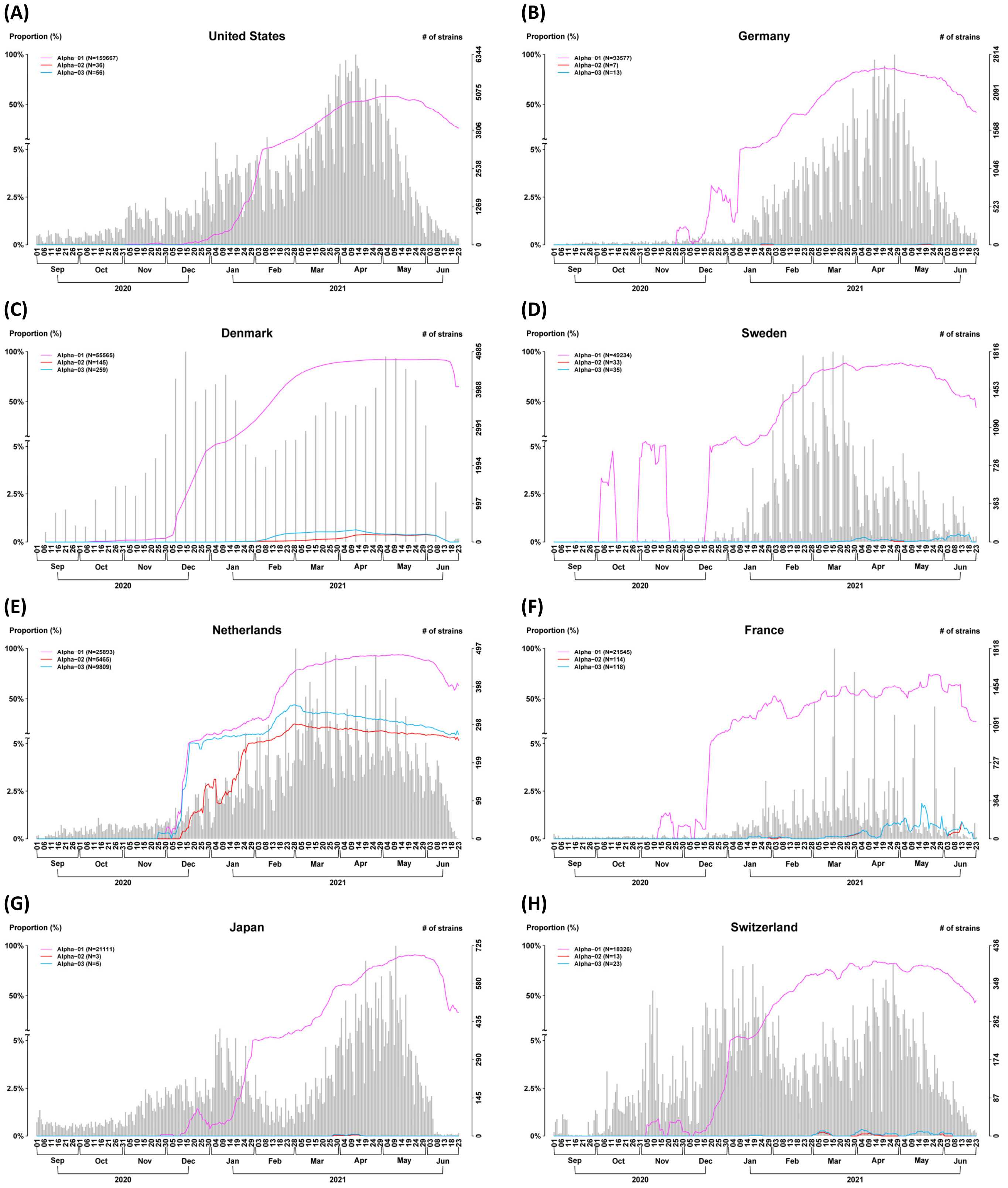
(A)
United States
Proportion (%)
### of strains
Alpha−01 (N=159667)
Alpha−02 (N=36)
Alpha−03 (N=56)
(B)
Germany
Alpha−01 (N=93577)
Alpha−02 (N=7)
Alpha−03 (N=13)
(C)
Denmark
Alpha−01 (N=55565)
Alpha−02 (N=145)
Alpha−03 (N=259)
(D)
Sweden
Alpha−01 (N=49234)
Alpha−02 (N=33)
Alpha−03 (N=35)
(E)
Netherlands
Alpha−01 (N=25893)
Alpha−02 (N=5465)
Alpha−03 (N=9809)
(F)
France
Alpha−01 (N=21545)
Alpha−02 (N=114)
Alpha−03 (N=118)
(G)
Japan
Alpha−01 (N=21111)
Alpha−02 (N=3)
Alpha−03 (N=5)
(H)
Switzerland
Alpha−01 (N=18326)
Alpha−02 (N=13)
Alpha−03 (N=23)
2020
2021

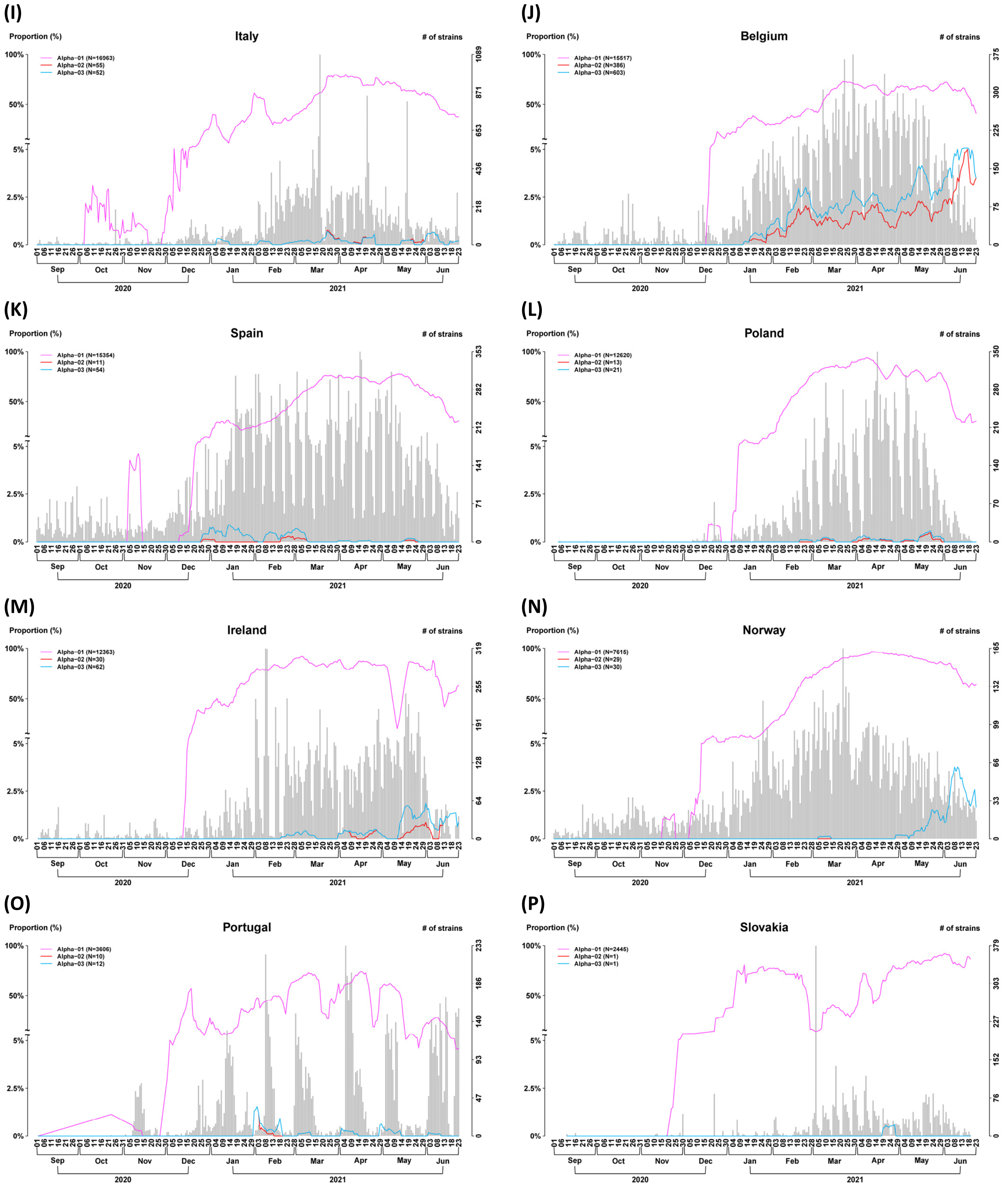
(I)
Italy
Proportion (%)
### of strains
Alpha−01 (N=16963)
Alpha−02 (N=55)
Alpha−03 (N=52)
(J)
Belgium
Proportion (%)
### of strains
Alpha−01 (N=15517)
Alpha−02 (N=386)
Alpha−03 (N=603)
(K)
Spain
Proportion (%)
### of strains
Alpha−01 (N=15354)
Alpha−02 (N=11)
Alpha−03 (N=54)
(L)
Poland
Proportion (%)
### of strains
Alpha−01 (N=12620)
Alpha−02 (N=13)
Alpha−03 (N=21)
(M)
Ireland
Proportion (%)
### of strains
Alpha−01 (N=12363)
Alpha−02 (N=30)
Alpha−03 (N=62)
(N)
Norway
Proportion (%)
### of strains
Alpha−01 (N=7615)
Alpha−02 (N=29)
Alpha−03 (N=30)
(O)
Portugal
Proportion (%)
### of strains
Alpha−01 (N=3606)
Alpha−02 (N=10)
Alpha−03 (N=12)
(P)
Slovakia
Proportion (%)
### of strains
Alpha−01 (N=2445)
Alpha−02 (N=1)
Alpha−03 (N=1)
Sep Oct Nov Dec Jan Feb Mar Apr May Jun
2020 2021

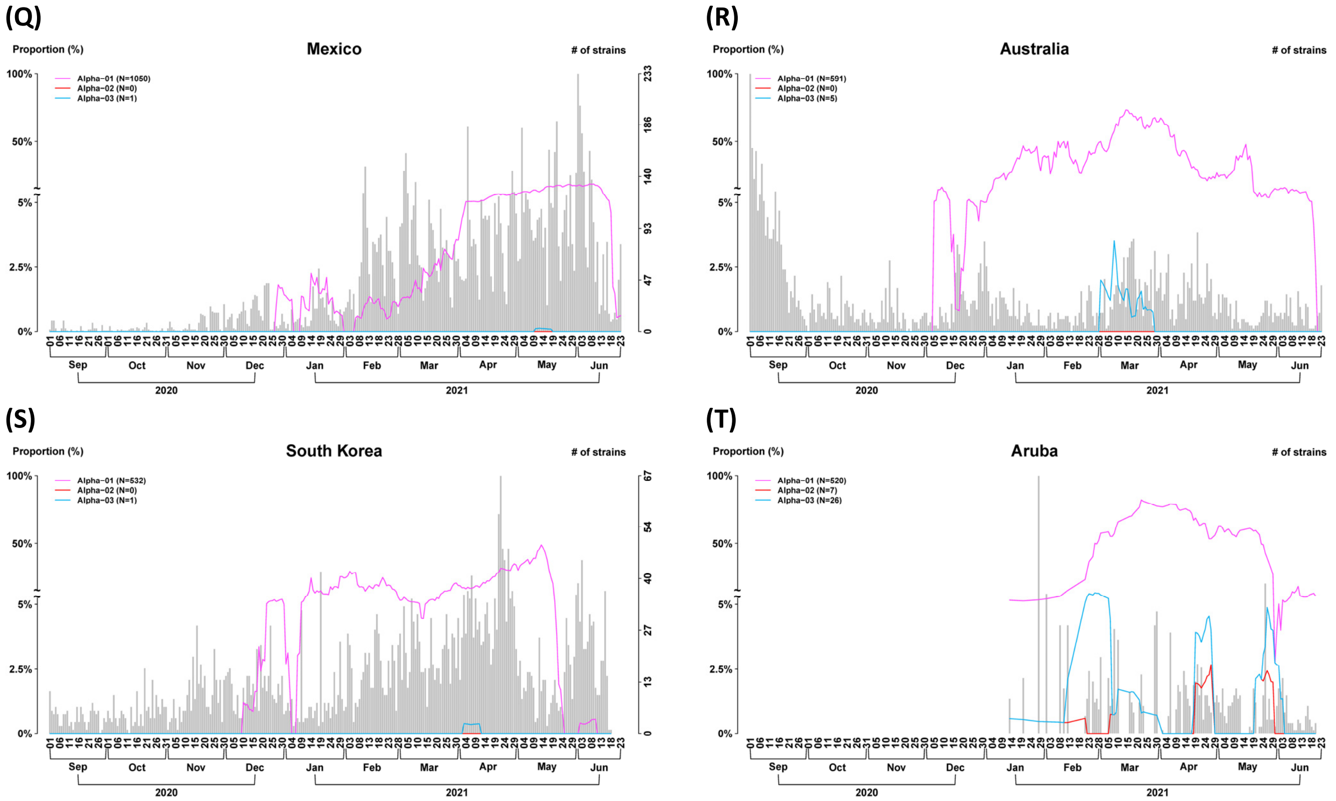


**Figure S11. Differential temporal trajectories for the three Alpha subtypes with transmission enhancer and/or suppressor SNVs in many countries (n = 2,119K).** The first increasing subtype has rising temporal trajectories (Alpha-01). The remaining two subtypes (Alpha-02 and Alpha-03) have a lower temporal trajectory. Remarkably, the same pattern of the differential temporal trajectories for the three subtypes was found not only in the United Kingdom but also in many other countries consistently. (A) The United States; (B) Germany; (C) Denmark; (D) Sweden; (E) The Netherland; (F) France; (G) Japan; (H) Switzerland; (I) Italy; (J) Belgium; (K) Spain; (L) Poland; (M) Ireland; (N) Norway; (O) Portugal; (P) Slovakia; (Q) Mexico; (R) Australia; (S) South Korea; (T) Aruba.

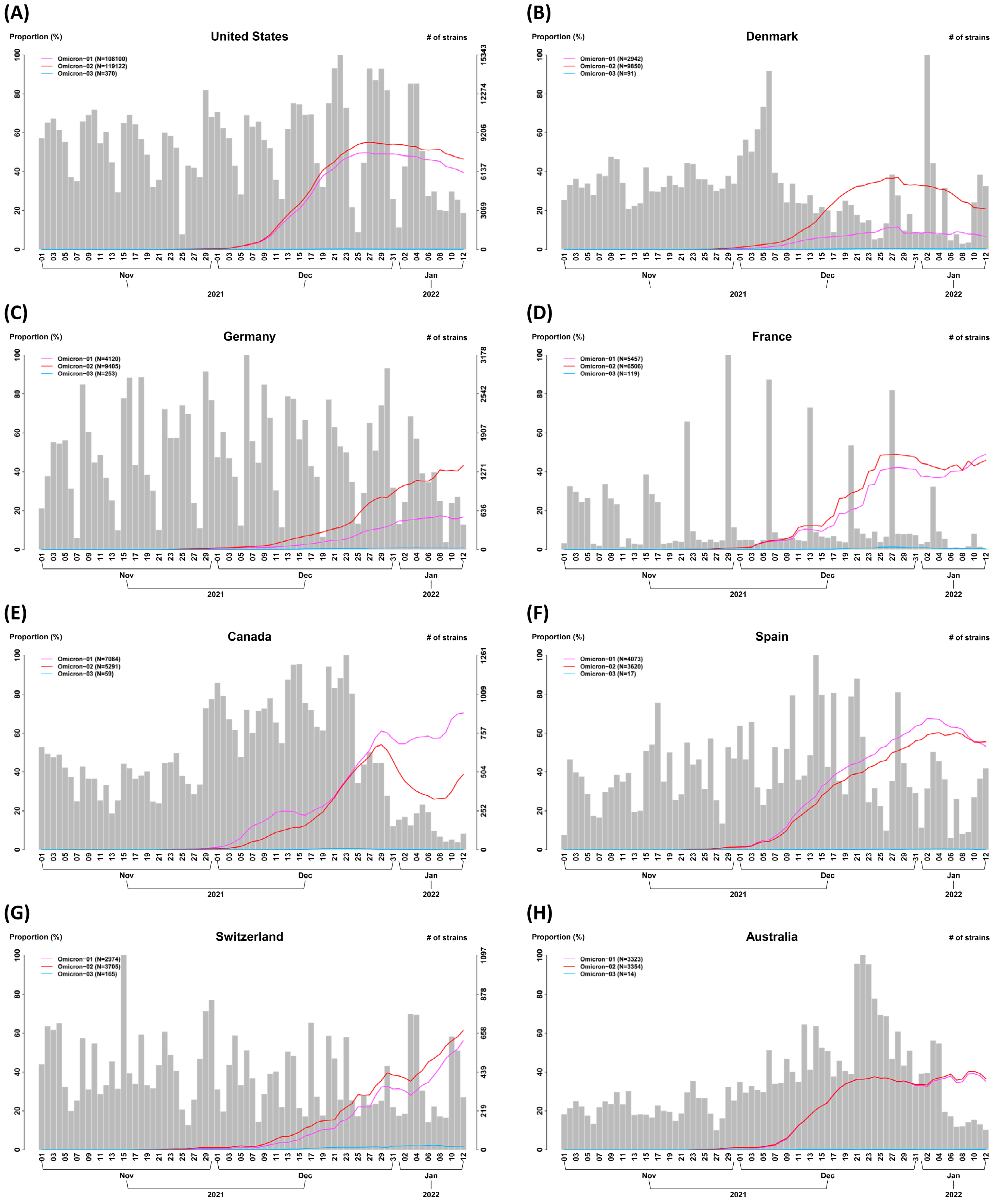
(A)
Proportion (%)
United States
### of strains
Omicron−01 (N=108100)
Omicron−02 (N=119122)
Omicron−03 (N=370)
(B)
Proportion (%)
Denmark
### of strains
Omicron−01 (N=2942)
Omicron−02 (N=9850)
Omicron−03 (N=91)
(C)
Proportion (%)
Germany
### of strains
Omicron−01 (N=4120)
Omicron−02 (N=9405)
Omicron−03 (N=253)
(D)
Proportion (%)
France
### of strains
Omicron−01 (N=5457)
Omicron−02 (N=6506)
Omicron−03 (N=119)
(E)
Proportion (%)
Canada
### of strains
Omicron−01 (N=7084)
Omicron−02 (N=5291)
Omicron−03 (N=59)
(F)
Proportion (%)
Spain
### of strains
Omicron−01 (N=4073)
Omicron−02 (N=3620)
Omicron−03 (N=17)
(G)
Proportion (%)
Switzerland
### of strains
Omicron−01 (N=2974)
Omicron−02 (N=3705)
Omicron−03 (N=165)
(H)
Proportion (%)
Australia
### of strains
Omicron−01 (N=3323)
Omicron−02 (N=3354)
Omicron−03 (N=14)
Nov
Dec
Jan
2021
2022

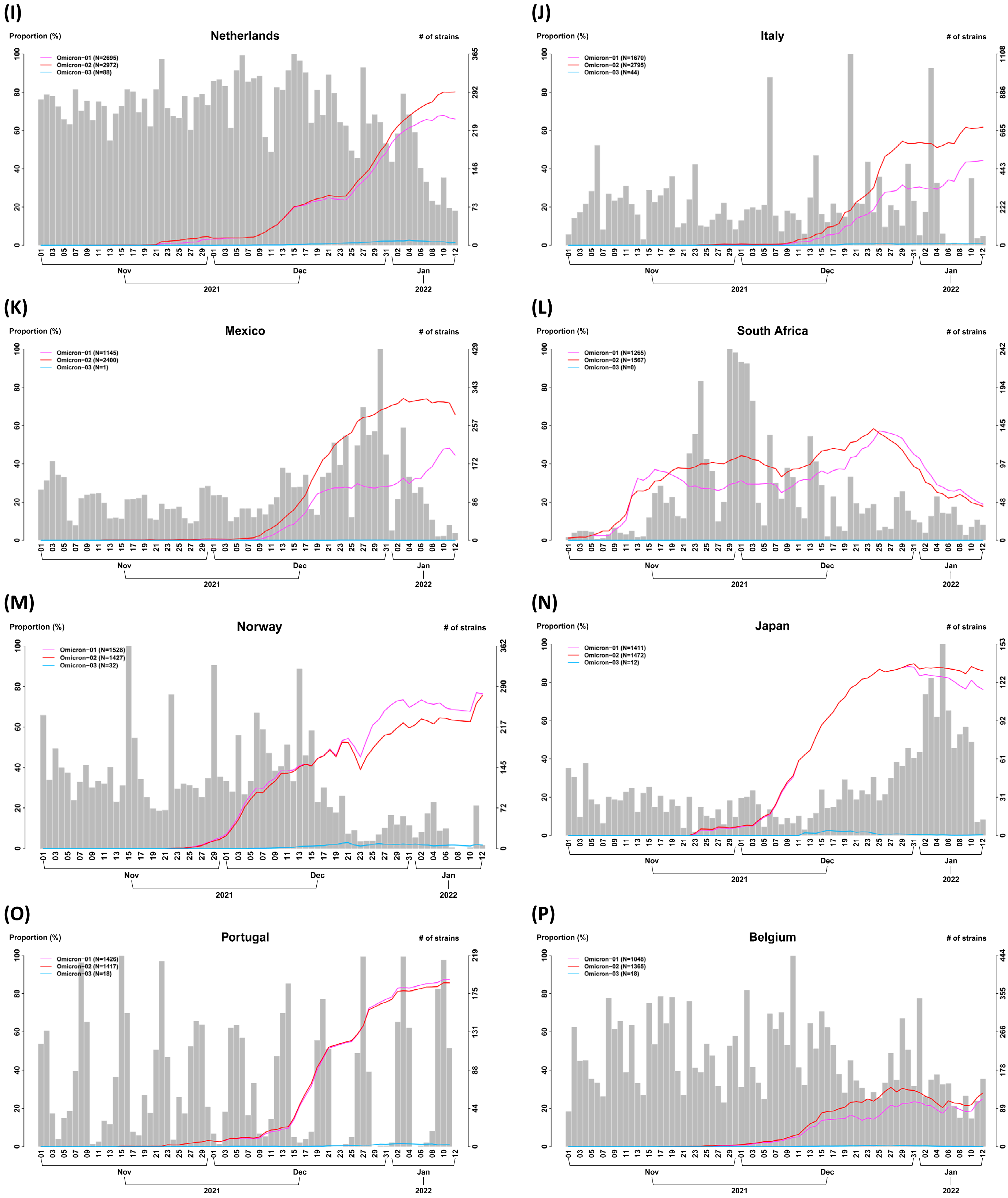
(I)
Netherlands
Proportion (%)
### of strains
Omicron−01 (N=2695)
Omicron−02 (N=2972)
Omicron−03 (N=88)
(J)
Italy
Proportion (%)
### of strains
Omicron−01 (N=1670)
Omicron−02 (N=2795)
Omicron−03 (N=44)
(K)
Mexico
Proportion (%)
### of strains
Omicron−01 (N=1145)
Omicron−02 (N=2400)
Omicron−03 (N=1)
(L)
South Africa
Proportion (%)
### of strains
Omicron−01 (N=1265)
Omicron−02 (N=1567)
Omicron−03 (N=0)
(M)
Norway
Proportion (%)
### of strains
Omicron−01 (N=1528)
Omicron−02 (N=1427)
Omicron−03 (N=32)
(N)
Japan
Proportion (%)
### of strains
Omicron−01 (N=1411)
Omicron−02 (N=1472)
Omicron−03 (N=12)
(O)
Portugal
Proportion (%)
### of strains
Omicron−01 (N=1426)
Omicron−02 (N=1417)
Omicron−03 (N=18)
(P)
Belgium
Proportion (%)
### of strains
Omicron−01 (N=1048)
Omicron−02 (N=1365)
Omicron−03 (N=18)
Nov
Dec
Jan
2021
2022

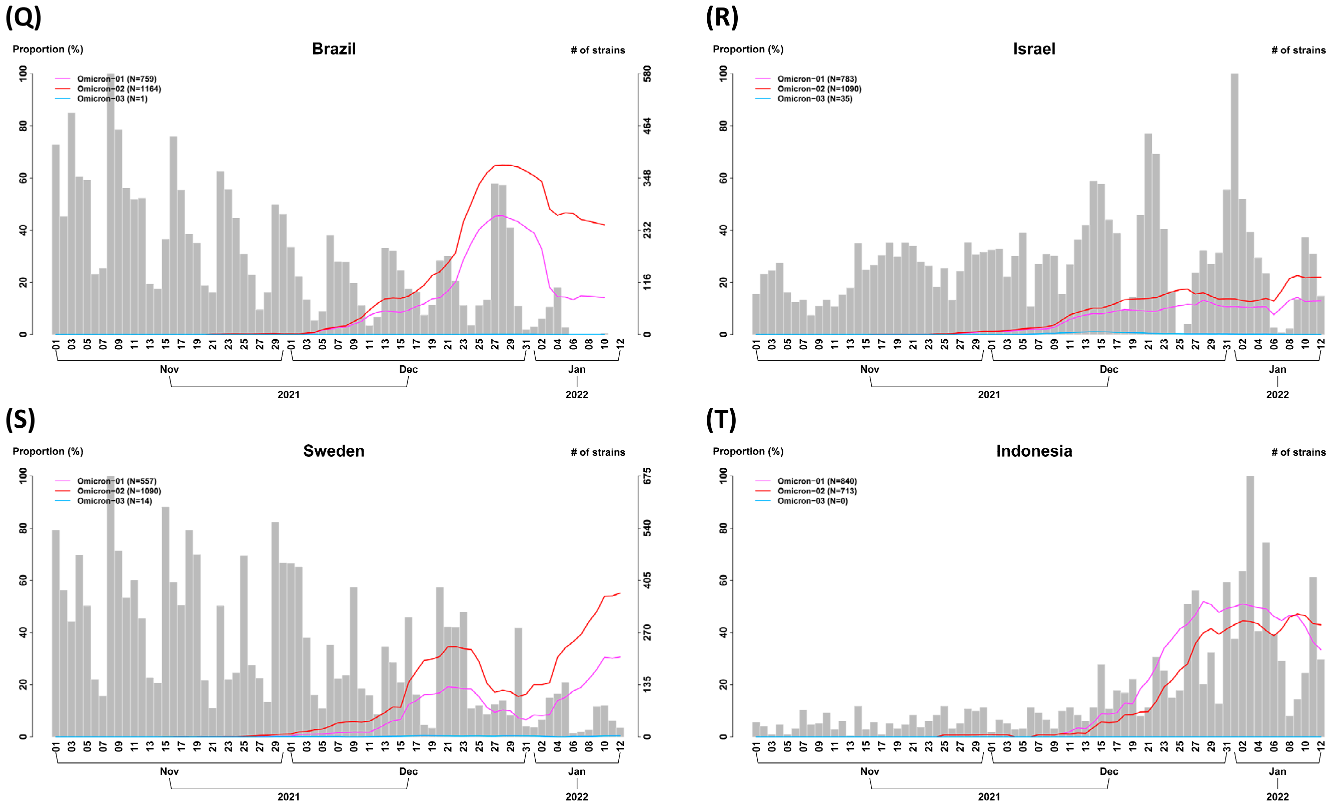


**Figure S12. Differential temporal trajectories for the three Omicron subtypes with transmission enhancer and/or suppressor SNVs in many countries (n = 7,026K).** The first two increasing subtypes have rising temporal trajectories (Omicron-01, and Omicron-02). The remaining subtype (Omicron-03) have a lower temporal trajectory. Remarkably, the same pattern of the differential temporal trajectories for the three subtypes was found not only in the United Kingdom but also in many other countries consistently. (A) The United States; (B) Denmark; (C) Germany; (D) France; (E) Canada; (F) Spain; (G) Switzerland; (H) Australia; (I) The Netherlands; (J) Italy; (K) Mexico; (L) South Africa; (M) Norway; (N) Japan; (O) Portugal; (P) Belgium; (Q) Brazil; (R) Israel; (S) Sweden; (T) Indonesia.

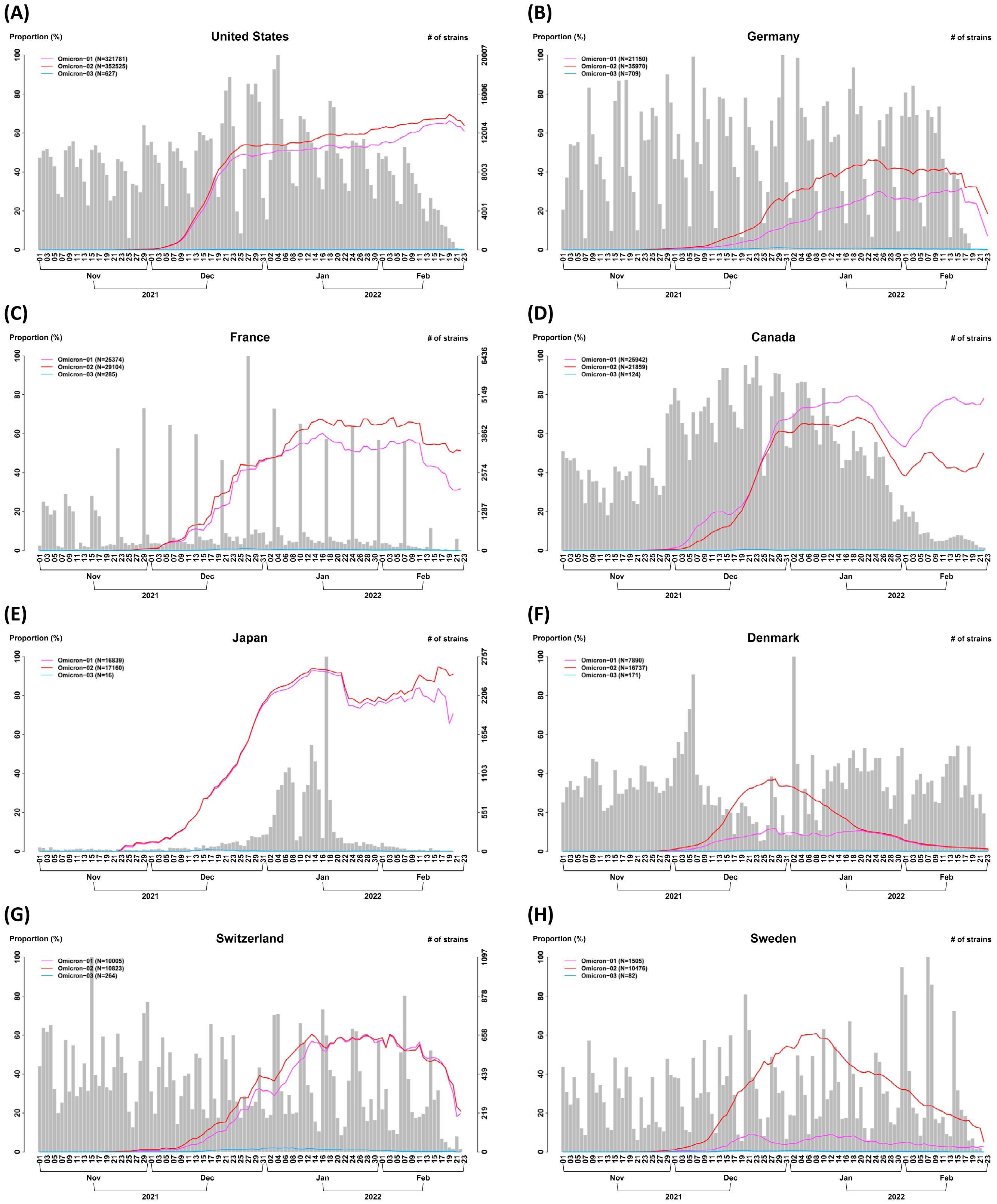
(A)
United States
Proportion (%)
### of strains
Omicron−01 (N=321781)
Omicron−02 (N=352525)
Omicron−03 (N=627)
(B)
Germany
Omicron−01 (N=21150)
Omicron−02 (N=35970)
Omicron−03 (N=709)
(C)
France
Omicron−01 (N=25374)
Omicron−02 (N=29104)
Omicron−03 (N=285)
(D)
Canada
Omicron−01 (N=25942)
Omicron−02 (N=21859)
Omicron−03 (N=124)
(E)
Japan
Omicron−01 (N=16839)
Omicron−02 (N=17160)
Omicron−03 (N=16)
(F)
Denmark
Omicron−01 (N=7890)
Omicron−02 (N=16737)
Omicron−03 (N=171)
(G)
Switzerland
Omicron−01 (N=10005)
Omicron−02 (N=10823)
Omicron−03 (N=264)
(H)
Sweden
Omicron−01 (N=1505)
Omicron−02 (N=10476)
Omicron−03 (N=82)
Nov
Dec
Jan
Feb
2021
2022

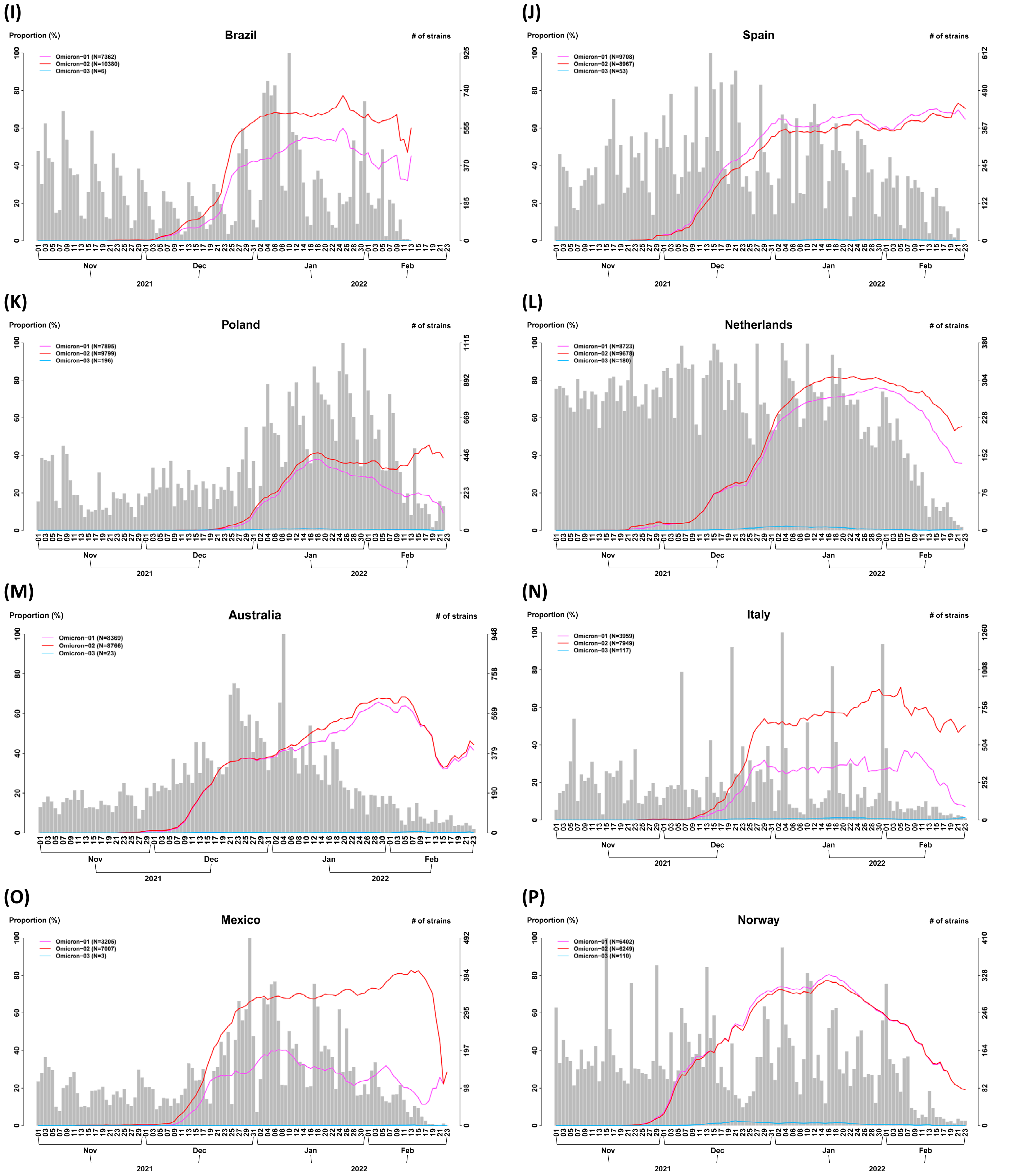
(I)
Brazil
Proportion (%)
### of strains
Omicron−01 (N=7362)
Omicron−02 (N=10380)
Omicron−03 (N=6)
(J)
Spain
Proportion (%)
### of strains
Omicron−01 (N=9708)
Omicron−02 (N=8967)
Omicron−03 (N=53)
(K)
Poland
Proportion (%)
### of strains
Omicron−01 (N=7895)
Omicron−02 (N=9799)
Omicron−03 (N=196)
(L)
Netherlands
Proportion (%)
### of strains
Omicron−01 (N=8723)
Omicron−02 (N=9678)
Omicron−03 (N=180)
(M)
Australia
Proportion (%)
### of strains
Omicron−01 (N=8369)
Omicron−02 (N=8766)
Omicron−03 (N=23)
(N)
Italy
Proportion (%)
### of strains
Omicron−01 (N=3959)
Omicron−02 (N=7949)
Omicron−03 (N=117)
(O)
Mexico
Proportion (%)
### of strains
Omicron−01 (N=3205)
Omicron−02 (N=7007)
Omicron−03 (N=3)
(P)
Norway
Proportion (%)
### of strains
Omicron−01 (N=6402)
Omicron−02 (N=6249)
Omicron−03 (N=110)
Nov
Dec
Jan
Feb
2021
2022

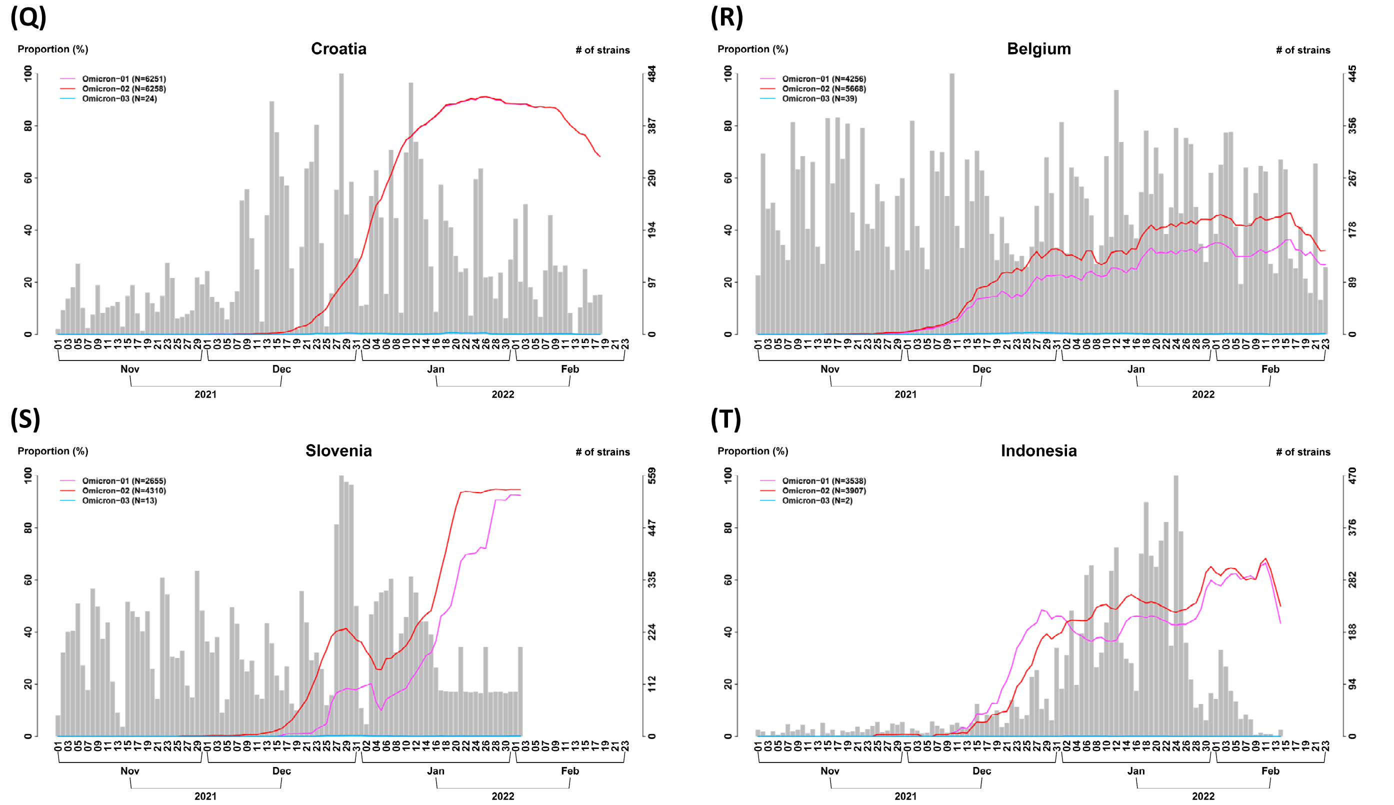


**Figure S13. Differential temporal trajectories for the three Omicron subtypes with transmission enhancer and/or suppressor SNVs in many countries (n = 8,475K).** The first two increasing subtypes have rising temporal trajectories (Omicron-01, and Omicron-02). The remaining subtype (Omicron-03) have a lower temporal trajectory. Remarkably, the same pattern of the differential temporal trajectories for the three subtypes was found not only in the United Kingdom but also in many other countries consistently. (A) The United States; (B) Germany; (C) France; (D) Canada; (E) Spain; (F) Denmark; (G) Switzerland; (H) Sweden; (I) Brazil; (J) Spain; (K) Poland; (L) Netherlands; (M) Australia; (N) Italy; (O) Mexico; (P) Norway; (Q) Croatia; (R) Belgium; (S) Slovenia; (T) Indonesia.

**(A)**

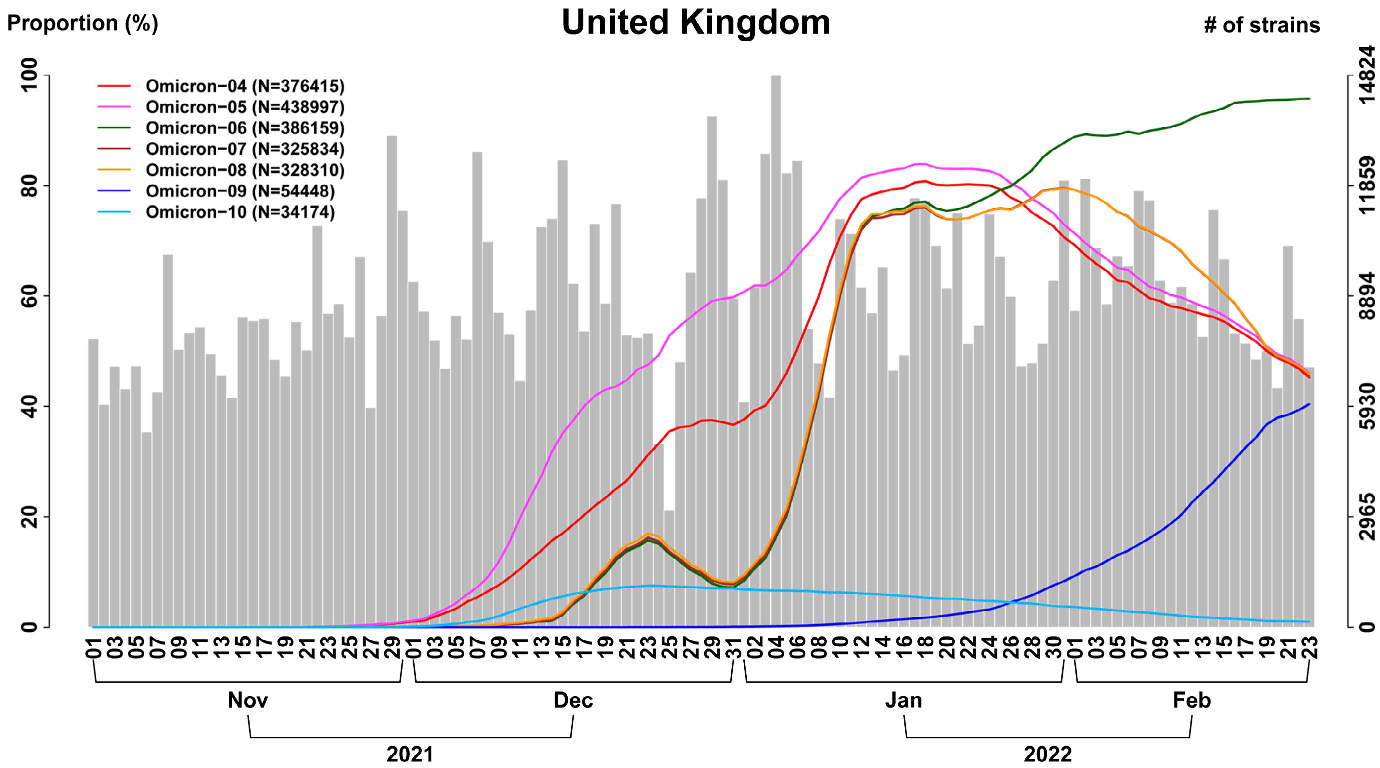


**(B)**

Defining SNVs for Omicron

Transmission enhancer

Transmission suppressor

Defining SNVs for CSSs

Red number — Proportion of SNV in a CSS is >0.5

Blue number — Proportion of SNV in a CSS is <0.5

Italic and bold: Synonymous SNV

| Omicron variant | | | Defining SNVs for Omicron (Y: yes) | Omicron-04 | Omicron-05 | Omicron-06 | Omicron-07 | Omicron-08 | Omicron-09 | Omicron-10 | Transmission related SNVs |
|---|---|---|---|---|---|---|---|---|---|---|---|
| | | | | High | High | High | High | High | High | Low | |
| A2832G | nsp3 | K856R | Y | 100.00% | 99.38% | 83.60% | 99.09% | 98.67% | 0.00% | 99.53% | Enhancer |
| T5386G | nsp3 | ***A1707A*** | Y | 100.00% | 99.89% | 84.12% | 99.83% | 99.64% | 0.06% | 99.87% | Enhancer |
| G8393A | nsp3 | A2710T | Y | 100.00% | 99.82% | 84.10% | 99.79% | 99.55% | 0.00% | 99.90% | Enhancer |
| C10449A | nsp5 | P3395H | Y | 100.00% | 100.00% | 100.00% | 99.76% | 99.49% | 99.97% | 99.60% | Enhancer |
| A11537G | nsp6 | I3758V | Y | 100.00% | 99.95% | 84.17% | 99.90% | 99.72% | 0.19% | 99.95% | Enhancer |
| T13195C | nsp10 | ***V4310V*** | Y | 100.00% | 99.88% | 84.06% | 99.80% | 99.66% | 0.00% | 99.39% | Enhancer |
| C15240T | nsp12 | ***N4992N*** | Y | 100.00% | 99.94% | 84.23% | 99.86% | 99.64% | 0.66% | 99.79% | Enhancer |
| A18163G | nsp14 | I5967V | Y | 100.00% | 100.00% | 100.00% | 99.52% | 99.41% | 99.83% | 89.06% | Enhancer |
| C21762T | S (NTD) | A67V | Y | 100.00% | 99.37% | 83.26% | 98.22% | 97.88% | 0.06% | 99.50% | Enhancer |
| C21767- | S (NTD) | DEL | | 100.00% | 98.73% | 82.64% | 97.46% | 96.85% | 0.12% | 97.91% | Enhancer |
| G22578A | S (RBD) | G339D | Y | 100.00% | 100.00% | 100.00% | 99.45% | 92.69% | 99.39% | 98.93% | Enhancer |
| T22673C | S (RBD) | S371P | Y | 100.00% | 100.00% | 84.10% | 100.00% | 92.03% | 0.00% | 93.79% | Enhancer |
| C22674T | S (RBD) | S371F | Y | 100.00% | 100.00% | 100.00% | 100.00% | 92.17% | 99.77% | 93.85% | Enhancer |
| T22679C | S (RBD) | S373P | Y | 100.00% | 100.00% | 100.00% | 99.96% | 92.52% | 100.00% | 94.02% | Enhancer |
| C22686T | S (RBD) | S375F | | 100.00% | 100.00% | 100.00% | 99.57% | 91.99% | 99.98% | 93.88% | Enhancer |
| G22992A | S (RBD, RBM) | S477N | Y | 100.00% | 100.00% | 100.00% | 99.60% | 97.67% | 100.00% | 93.68% | Enhancer |
| A23013C | S (RBD, RBM) | E484A | Y | 100.00% | 100.00% | 100.00% | 99.53% | 97.45% | 99.84% | 93.49% | Enhancer |
| A23040G | S (RBD, RBM) | Q493R | Y | 100.00% | 100.00% | 100.00% | 99.79% | 96.47% | 99.67% | 93.47% | Enhancer |
| G23048A | S (RBD, RBM) | G496S | Y | 100.00% | 100.00% | 83.63% | 100.00% | 95.98% | 0.02% | 93.45% | Enhancer |
| A23055G | S (RBD, RBM) | Q498R | Y | 100.00% | 100.00% | 92.88% | 93.36% | 89.93% | 95.85% | 87.00% | Enhancer |
| T23075C | S (RBD, RBM) | Y505H | | 100.00% | 100.00% | 92.85% | 92.99% | 90.90% | 95.96% | 87.26% | Enhancer |
| C23202A | S (S1) | T547K | Y | 100.00% | 100.00% | 84.06% | 99.77% | 99.52% | 0.03% | 99.93% | Enhancer |
| C23525T | S (S1) | H655Y | Y | 100.00% | 100.00% | 100.00% | 99.78% | 99.74% | 100.00% | 100.00% | Enhancer |
| T23599G | S (S1) | N679K | Y | 100.00% | 100.00% | 100.00% | 99.77% | 99.77% | 100.00% | 99.99% | Enhancer |
| C23604A | S (S1) | P681H | Y | 100.00% | 100.00% | 99.58% | 99.30% | 99.33% | 100.00% | 100.00% | Enhancer |
| C23854A | S (S2) | N764K | Y | 100.00% | 91.69% | 100.00% | 100.00% | 100.00% | 99.86% | 78.25% | Enhancer |
| G23948T | S (FP) | D796Y | Y | 100.00% | 100.00% | 100.00% | 99.65% | 99.34% | 100.00% | 100.00% | Enhancer |
| C24130A | S (S2) | N856K | Y | 100.00% | 99.89% | 84.06% | 99.71% | 99.43% | 0.00% | 100.00% | Enhancer |
| A24424T | S (HR1) | Q954H | Y | 100.00% | 100.00% | 100.00% | 100.00% | 99.64% | 100.00% | 99.96% | Enhancer |
| T24469A | S (HR1) | N969K | Y | 100.00% | 100.00% | 100.00% | 99.87% | 99.55% | 99.99% | 99.99% | Enhancer |
| C24503T | S (HR1) | L981F | | 100.00% | 99.93% | 84.12% | 99.70% | 99.31% | 0.00% | 100.00% | Enhancer |
| C25000T | S (S2) | ***D1146D*** | Y | 100.00% | 100.00% | 100.00% | 99.29% | 98.74% | 100.00% | 99.55% | Enhancer |
| C25584T | ORF3a | ***T64T*** | | 100.00% | 100.00% | 100.00% | 99.88% | 99.68% | 99.99% | 99.93% | Enhancer |

| | | | | | | | | | | | |
|---|---|---|---|---|---|---|---|---|---|---|---|
| C26270T | E | T9I | Y | 100.00% | 100.00% | 100.00% | 99.66% | 98.93% | 99.85% | 98.67% | Enhancer |
| A26530G | M | D3G | Y | 100.00% | 98.23% | 82.70% | 100.00% | 96.36% | 0.00% | 95.76% | Enhancer |
| C26577 G | M | Q19E | Y | 100.00% | 100.00% | 100.00% | 98.62% | 97.24% | 92.81% | 96.10% | Enhancer |
| G26709A | M | A63T | Y | 100.00% | 100.00% | 100.00% | 99.83% | 99.32% | 99.97% | 96.53% | Enhancer |
| A27259C | ORF6 | ***R20R*** | Y | 100.00% | 100.00% | 100.00% | 99.50% | 99.62% | 99.72% | 99.72% | Enhancer |
| C27807T | ORF7b | ***L18L*** | Y | 100.00% | 100.00% | 100.00% | 99.43% | 99.31% | 99.79% | 97.88% | Enhancer |
| A28271T | - | - | | 100.00% | 100.00% | 100.00% | 96.45% | 96.56% | 99.28% | 99.02% | Enhancer |
| C28311T | N | P13L | | 100.00% | 100.00% | 100.00% | 96.30% | 96.46% | 99.31% | 99.01% | Enhancer |
| G28881A | N | R203K | Y | 100.00% | 100.00% | 99.76% | 99.62% | 99.52% | 99.78% | 99.85% | Enhancer |
| G28882A | N | ***R203R*** | Y | 100.00% | 100.00% | 99.70% | 99.59% | 99.34% | 99.78% | 99.77% | Enhancer |
| G28883 C | N | G204R | Y | 100.00% | 100.00% | 99.70% | 99.59% | 99.36% | 99.78% | 99.76% | Enhancer |
| A23063T | S (RBD, RBM) | N501Y | Y | 99.99% | 100.00% | 92.96% | 93.43% | 90.23% | 95.99% | 87.15% | Enhancer |
| G22813T | S (RBD) | K417N | Y | 75.12% | 69.59% | 100.00% | 94.11% | 100.00% | 100.00% | 49.16% | Enhancer |
| T22882G | S (RBD, RBM) | N440K | Y | 79.05% | 73.36% | 100.00% | 98.88% | 100.00% | 96.96% | 49.22% | Enhancer |
| G22898A | S (RBD, RBM) | G446S | Y | 79.48% | 73.79% | 83.98% | 100.00% | 100.00% | 0.02% | 49.41% | Enhancer |
| T670G | nsp1 | S135R | | 0.00% | 0.00% | 15.58% | 0.00% | 0.06% | 100.00% | 0.00% | Enhancer |
| C2790T | nsp3 | T842I | | 0.00% | 0.01% | 15.67% | 0.01% | 0.03% | 100.00% | 0.00% | Enhancer |
| G4184A | nsp3 | G1307S | | 0.01% | 0.01% | 15.73% | 0.01% | 0.07% | 100.00% | 0.00% | Enhancer |
| C4321T | nsp3 | ***A1352A*** | | 0.03% | 0.03% | 15.68% | 0.03% | 0.08% | 100.00% | 0.07% | Enhancer |
| C9344T | nsp4 | ***L3027F*** | | 0.06% | 0.06% | 15.74% | 0.07% | 0.09% | 100.00% | 0.03% | Enhancer |
| A9424G | nsp4 | ***V3053V*** | | 0.01% | 0.01% | 15.63% | 0.01% | 0.04% | 100.00% | 0.00% | Enhancer |
| C9534T | nsp4 | T3090I | | 0.04% | 0.04% | 15.54% | 0.04% | 0.09% | 100.00% | 0.04% | Enhancer |
| C9866T | nsp4 | L3201F | | 0.01% | 0.01% | 15.52% | 0.01% | 0.04% | 100.00% | 0.00% | Enhancer |
| C10198T | nsp5 | ***D3311D*** | | 0.02% | 0.02% | 14.65% | 0.02% | 0.04% | 100.00% | 0.01% | Enhancer |
| G10447A | nsp5 | ***R3394R*** | | 0.01% | 0.01% | 15.71% | 0.01% | 0.07% | 100.00% | 0.00% | Enhancer |
| C12880T | nsp9 | ***I4205I*** | | 0.13% | 0.12% | 15.87% | 0.16% | 0.21% | 100.00% | 0.01% | Enhancer |
| C15714T | nsp12 | ***L5150L*** | | 0.03% | 0.03% | 15.78% | 0.03% | 0.09% | 100.00% | 0.04% | Enhancer |
| C17410T | nsp13 | R5716C | | 0.15% | 0.14% | 15.89% | 0.14% | 0.16% | 100.00% | 0.03% | Enhancer |
| C19955T | nsp15 | T6564I | | 0.04% | 0.04% | 14.90% | 0.04% | 0.06% | 100.00% | 0.03% | Enhancer |
| A20055G | nsp15 | ***E6597E*** | | 0.04% | 0.04% | 14.56% | 0.05% | 0.07% | 100.00% | 0.02% | Enhancer |
| C21618T | S (NTD) | T19I | | 0.02% | 0.02% | 15.73% | 0.02% | 0.04% | 100.00% | 0.01% | Enhancer |
| A21634- | S (NTD) | DEL | | 0.00% | 0.01% | 14.76% | 0.05% | 0.06% | 100.00% | 0.00% | Enhancer |
| C21635- | S (NTD) | DEL | | 0.01% | 0.01% | 14.77% | 0.05% | 0.07% | 100.00% | 0.00% | Enhancer |
| C21636- | S (NTD) | DEL | | 0.00% | 0.01% | 14.76% | 0.05% | 0.06% | 100.00% | 0.00% | Enhancer |
| C21637- | S (NTD) | DEL | | 0.00% | 0.01% | 14.76% | 0.05% | 0.06% | 100.00% | 0.00% | Enhancer |
| C21638- | S (NTD) | DEL | | 0.00% | 0.01% | 14.76% | 0.04% | 0.06% | 100.00% | 0.00% | Enhancer |
| C21639- | S (NTD) | DEL | | 0.00% | 0.01% | 14.76% | 0.04% | 0.05% | 100.00% | 0.00% | Enhancer |
| T21640- | S (NTD) | DEL | | 0.00% | 0.01% | 14.74% | 0.04% | 0.05% | 100.00% | 0.00% | Enhancer |
| G21641- | S (NTD) | DEL | | 0.00% | 0.01% | 14.74% | 0.04% | 0.05% | 100.00% | 0.00% | Enhancer |
| G21987A | S (NTD) | G142D | | 0.01% | 0.02% | 15.71% | 0.06% | 0.12% | 100.00% | 0.02% | Enhancer |
| T22200G | S (NTD) | V213G | | 0.00% | 0.01% | 15.69% | 0.01% | 0.04% | 100.00% | 0.01% | Enhancer |
| A22688G | S (RBD) | T376A | | 0.01% | 0.01% | 15.72% | 0.01% | 0.04% | 100.00% | 0.01% | Enhancer |
| G22775A | S (RBD) | D405N | | 0.01% | 0.01% | 15.74% | 0.02% | 0.08% | 100.00% | 0.01% | Enhancer |
| A22786C | S (RBD) | R408S | | 0.00% | 0.00% | 15.69% | 0.00% | 0.03% | 100.00% | 0.01% | Enhancer |
| C22995A | S (RBD, RBM) | T478K | Y | 99.98% | 99.97% | 99.98% | 99.59% | 97.74% | 100.00% | 93.72% | Enhancer |
| A23403G | S (S1) | D614G | Y | 99.99% | 99.99% | 99.98% | 99.92% | 99.79% | 100.00% | 99.98% | Enhancer |
| C26060T | ORF3a | T223I | | 0.10% | 0.10% | 15.84% | 0.11% | 0.16% | 100.00% | 0.02% | Enhancer |
| C26858T | M | ***F112F*** | | 0.03% | 0.03% | 15.78% | 0.03% | 0.09% | 100.00% | 0.01% | Enhancer |
| G27382 C | ORF6 | D61H | | 0.29% | 0.28% | 15.99% | 0.32% | 0.33% | 100.00% | 0.31% | Enhancer |
| A27383T | ORF6 | D61V | | 0.29% | 0.28% | 15.95% | 0.31% | 0.33% | 100.00% | 0.28% | Enhancer |
| T27384C | ORF6 | ***D61D*** | | 1.52% | 1.48% | 16.98% | 1.47% | 1.48% | 100.00% | 0.62% | Enhancer |
| A29510C | N | S413R | | 0.00% | 0.00% | 15.71% | 0.00% | 0.06% | 100.00% | 0.00% | Enhancer |
| T10135C | nsp5 | ***L3290L*** | | 10.68% | 10.58% | 7.89% | 9.74% | 9.43% | 0.00% | 100.00% | Suppressor |
| A24803G | S (CD) | I1081V | | 2.41% | 2.63% | 1.84% | 2.01% | 1.92% | 0.00% | 100.00% | Suppressor |
| C25708T | ORF3a | L106F | | 10.78% | 10.67% | 7.97% | 9.83% | 9.52% | 0.03% | 100.00% | Suppressor |
| A29301G | N | D343G | | 10.70% | 10.60% | 7.92% | 9.76% | 9.48% | 0.04% | 100.00% | Suppressor |
| N: # of strains in each of the six CSSs in the world | | | | 896825 | 1022153 | 907618 | 875834 | 962632 | 120063 | 38318 | |
| n: # of strains in each of the six CSSs in UK | | | | 376415 | 438997 | 386159 | 325834 | 328310 | 54448 | 34174 | |

**(C)**

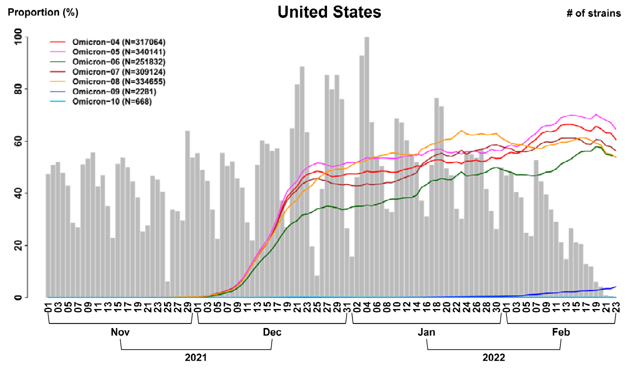


**(D)**

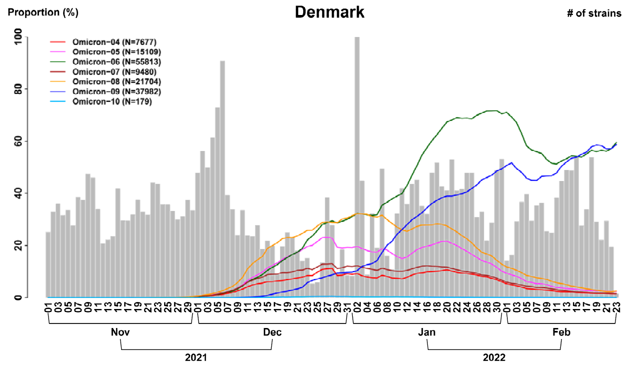

(E)

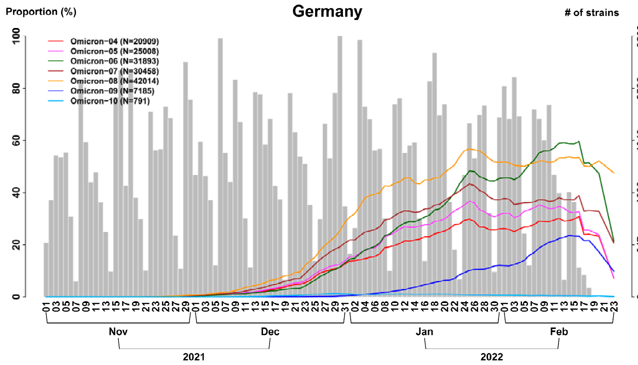
Proportion (%)
Germany
### of strains
Omicron−04 (N=20909)
Omicron−05 (N=25008)
Omicron−06 (N=31893)
Omicron−07 (N=30458)
Omicron−08 (N=42014)
Omicron−09 (N=7185)
Omicron−10 (N=791)
Nov
Dec
Jan
Feb
2021
2022

(F)

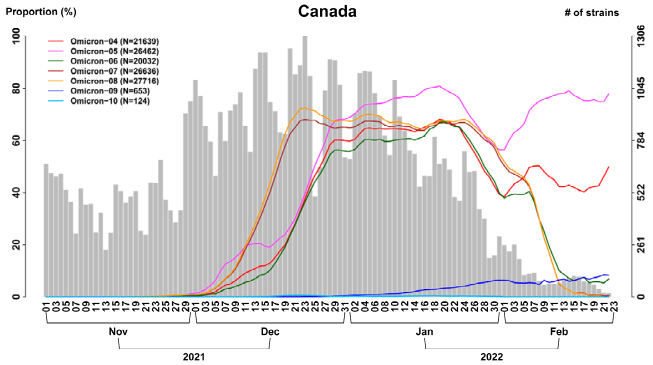
Proportion (%)
Canada
### of strains
Omicron−04 (N=21639)
Omicron−05 (N=26462)
Omicron−06 (N=20032)
Omicron−07 (N=26636)
Omicron−08 (N=27716)
Omicron−09 (N=653)
Omicron−10 (N=124)
Nov
Dec
Jan
Feb
2021
2022

(G)

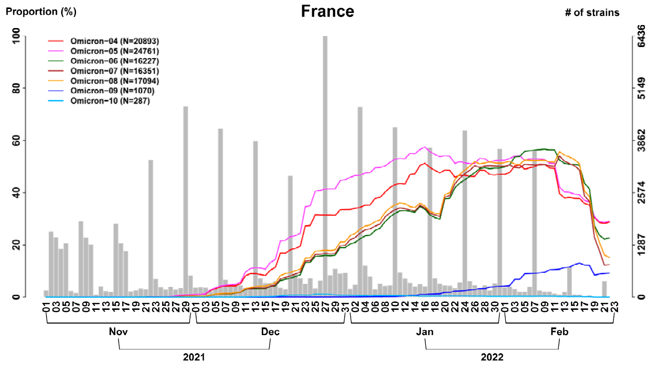
Proportion (%)
France
### of strains
Omicron−04 (N=20893)
Omicron−05 (N=24761)
Omicron−06 (N=16227)
Omicron−07 (N=16351)
Omicron−08 (N=17094)
Omicron−09 (N=1070)
Omicron−10 (N=287)
Nov
Dec
Jan
Feb
2021
2022

(H)

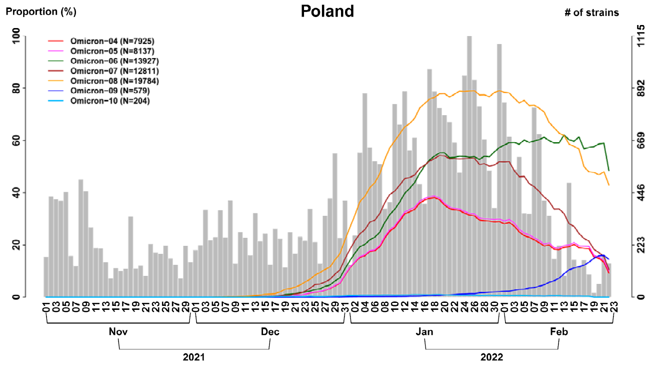
Proportion (%)
Poland
### of strains
Omicron−04 (N=7925)
Omicron−05 (N=8137)
Omicron−06 (N=13927)
Omicron−07 (N=12811)
Omicron−08 (N=19784)
Omicron−09 (N=579)
Omicron−10 (N=204)
Nov
Dec
Jan
Feb
2021
2022

(I)

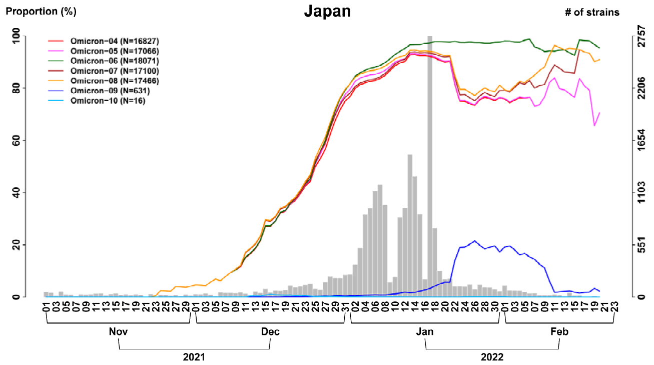
Proportion (%)
Japan
### of strains
Omicron−04 (N=16827)
Omicron−05 (N=17066)
Omicron−06 (N=18071)
Omicron−07 (N=17100)
Omicron−08 (N=17466)
Omicron−09 (N=631)
Omicron−10 (N=16)
Nov
Dec
Jan
Feb
2021
2022

(J)

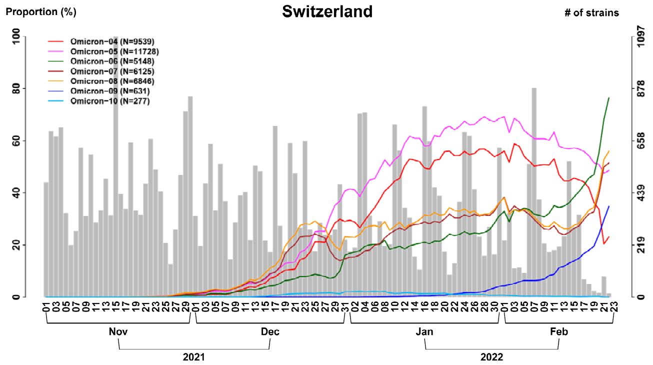
Proportion (%)
Switzerland
### of strains
Omicron−04 (N=9539)
Omicron−05 (N=11728)
Omicron−06 (N=5148)
Omicron−07 (N=6125)
Omicron−08 (N=6846)
Omicron−09 (N=631)
Omicron−10 (N=277)
Nov
Dec
Jan
Feb
2021
2022

(K)

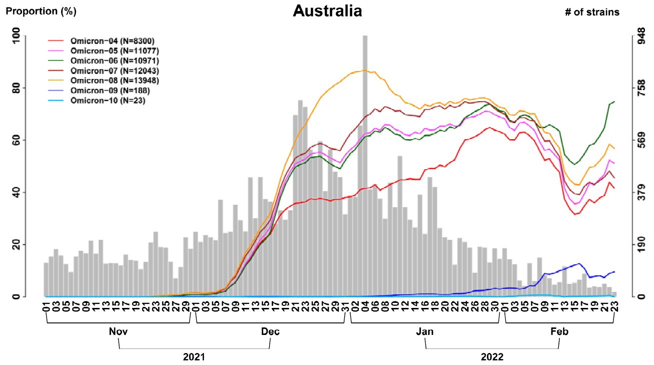
Proportion (%)
Australia
### of strains
Omicron−04 (N=8300)
Omicron−05 (N=11077)
Omicron−06 (N=10971)
Omicron−07 (N=12043)
Omicron−08 (N=13948)
Omicron−09 (N=188)
Omicron−10 (N=23)
Nov
Dec
Jan
Feb
2021
2022

(L)

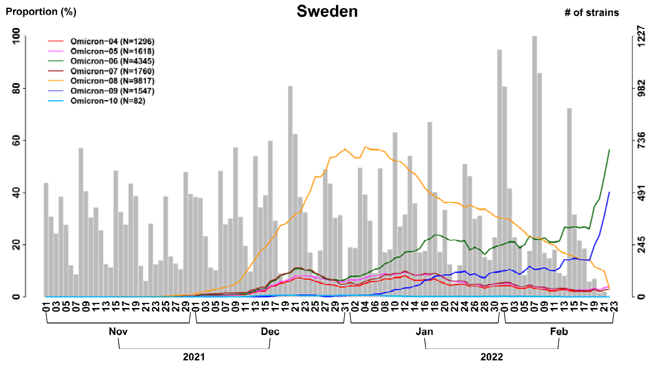
Proportion (%)
Sweden
### of strains
Omicron−04 (N=1296)
Omicron−05 (N=1618)
Omicron−06 (N=4345)
Omicron−07 (N=1760)
Omicron−08 (N=9817)
Omicron−09 (N=1547)
Omicron−10 (N=82)
Nov
Dec
Jan
Feb
2021
2022

(M)

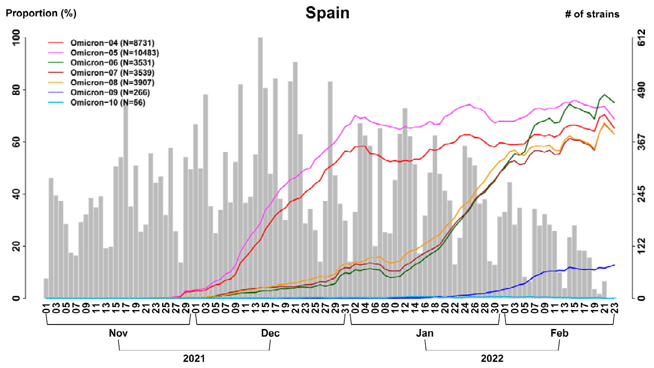
Proportion (%)
Spain
### of strains
Omicron−04 (N=8731)
Omicron−05 (N=10483)
Omicron−06 (N=3531)
Omicron−07 (N=3539)
Omicron−08 (N=3907)
Omicron−09 (N=266)
Omicron−10 (N=56)
Nov
Dec
Jan
Feb
2021
2022

(N)

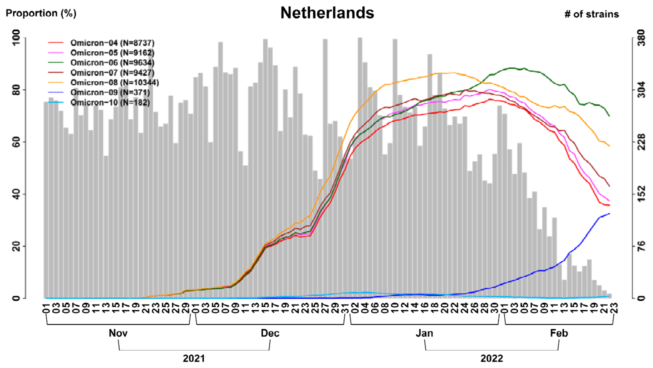
Proportion (%)
Netherlands
### of strains
Omicron−04 (N=8737)
Omicron−05 (N=9162)
Omicron−06 (N=9634)
Omicron−07 (N=9427)
Omicron−08 (N=10344)
Omicron−09 (N=371)
Omicron−10 (N=182)
Nov
Dec
Jan
Feb
2021
2022

(O)

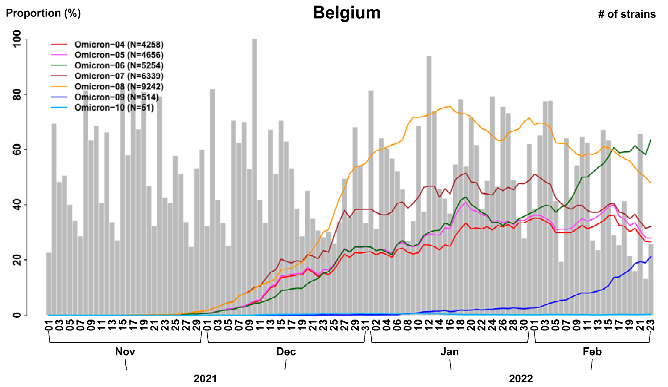
Proportion (%)
Belgium
### of strains
Omicron−04 (N=4258)
Omicron−05 (N=4656)
Omicron−06 (N=5254)
Omicron−07 (N=6339)
Omicron−08 (N=9242)
Omicron−09 (N=514)
Omicron−10 (N=51)
Nov
Dec
Jan
Feb
2021
2022

(P)

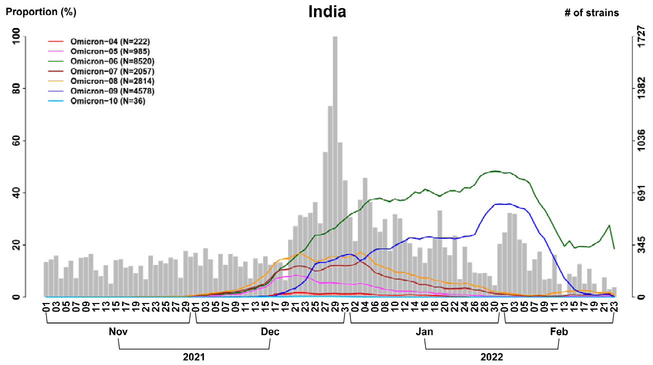
Proportion (%)
India
### of strains
Omicron−04 (N=222)
Omicron−05 (N=985)
Omicron−06 (N=8520)
Omicron−07 (N=2057)
Omicron−08 (N=2814)
Omicron−09 (N=4578)
Omicron−10 (N=36)
Nov
Dec
Jan
Feb
2021
2022

(Q)

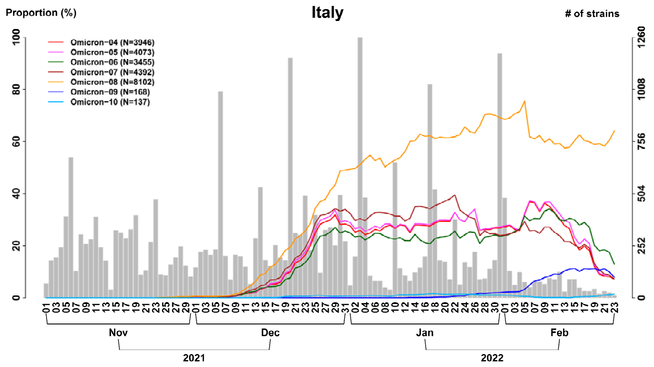
Proportion (%)
Italy
### of strains
Omicron−04 (N=3946)
Omicron−05 (N=4073)
Omicron−06 (N=3455)
Omicron−07 (N=4392)
Omicron−08 (N=8102)
Omicron−09 (N=168)
Omicron−10 (N=137)
Nov
Dec
Jan
Feb
2021
2022

(R)

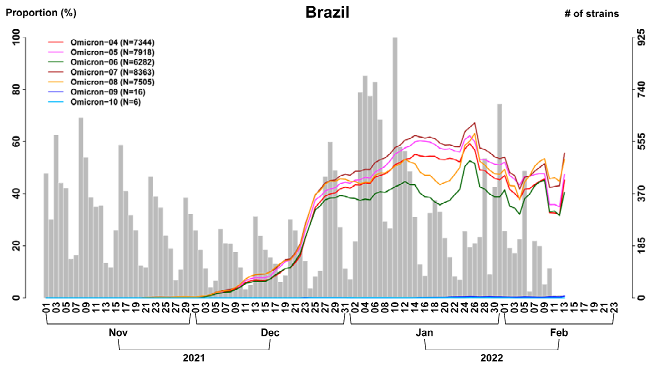
Proportion (%)
Brazil
### of strains
Omicron−04 (N=7344)
Omicron−05 (N=7918)
Omicron−06 (N=6282)
Omicron−07 (N=8363)
Omicron−08 (N=7505)
Omicron−09 (N=16)
Omicron−10 (N=6)
Nov
Dec
Jan
Feb
2021
2022

(S)

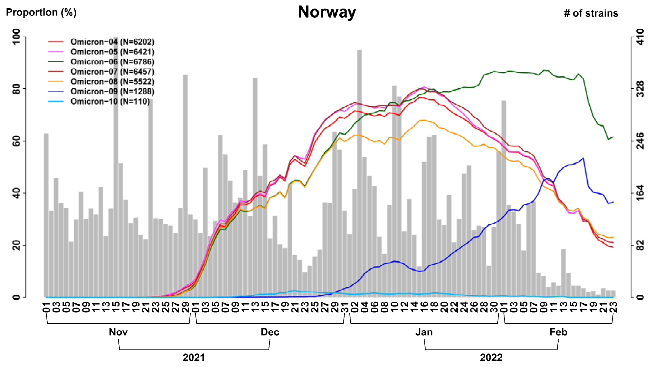
Proportion (%)
Norway
### of strains
Omicron−04 (N=6202)
Omicron−05 (N=6421)
Omicron−06 (N=6786)
Omicron−07 (N=6457)
Omicron−08 (N=5522)
Omicron−09 (N=1288)
Omicron−10 (N=110)
Nov
Dec
Jan
Feb
2021
2022

(T)

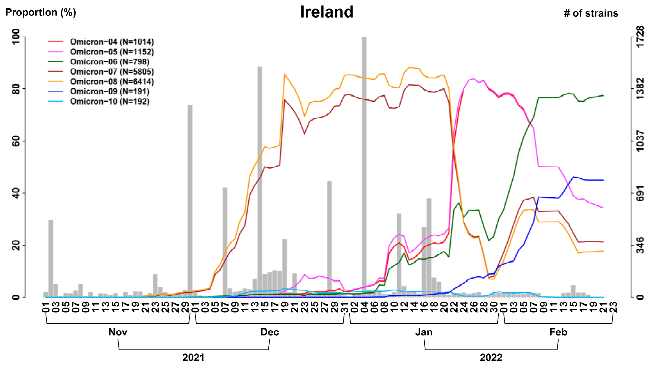
Proportion (%)
Ireland
### of strains
Omicron−04 (N=1014)
Omicron−05 (N=1152)
Omicron−06 (N=798)
Omicron−07 (N=5805)
Omicron−08 (N=6414)
Omicron−09 (N=191)
Omicron−10 (N=192)
Nov
Dec
Jan
Feb
2021
2022

(U)

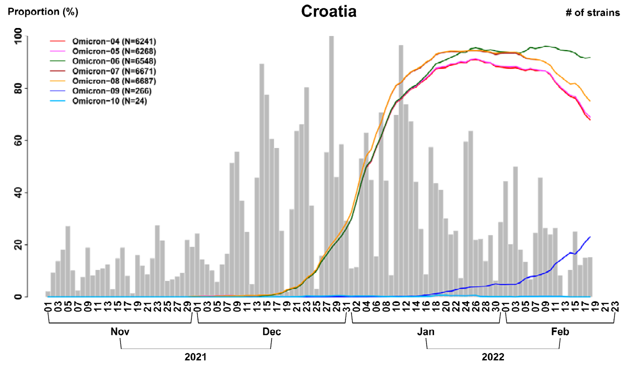


(V)

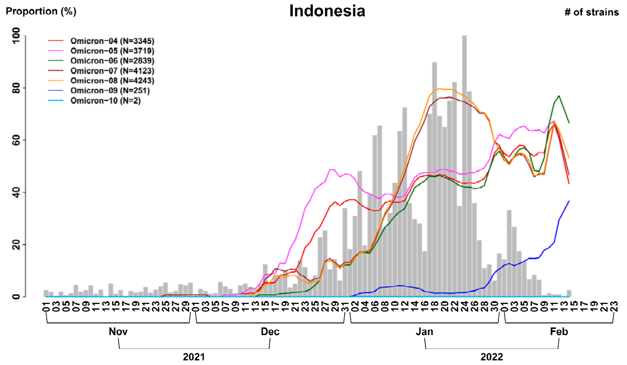


**Figure S14. Differential temporal trajectories for the seven Omicron subtypes with transmission enhancer and/or suppressor SNVs in a larger sample size (n = 8,475K). (A) Seven Omicron (aka B.1.1.529) subtypes (i.e., correlated SNV sets; CSSs) in the United Kingdom.** Six CSSs (Omicron-04 ~ Omicron-09) have an increasing temporal trajectory and one CSS (Omicron-10) have a much lower temporal trajectory, revealing that CSSs provide a more detailed information for the subtypes and their transmission patterns. **(B) Transmission enhancer and suppressor SNVs.** Seven CSSs are separated into two categories: the first six CSSs (Omicron-04 ~ Omicron-09) have a high temporal trajectory and the remaining one CSS has a low temporal trajectory. For each of the seven CSSs, nucleotide change, the protein that a SNV is located, and amino acid change for the signature SNVs are listed and followed by the proportions of the signature SNVs in each of the three CSSs. If the proportion is greater than 80%, then the numbers are marked by red color, otherwise, the numbers are marked by blue color. The signature SNVs for each of the seven CSSs are marked by yellow color. The number of strains in each of the seven CSSs in the world (N) and in the United Kingdom (n) are listed. Differential temporal trajectories for the seven Omicron subtypes with transmission enhancer and/or suppressor SNVs in many countries are provided as follows: (**C**) The United States; (**D**) Denmark; (**E**) Germany; (**F**) Canada; (**G**) France; (**H**) Poland; (**I**) Japan; (**J**) Switzerland; (**K**) Australia; (**L**) Sweden; (**M**) Spain; (**N**) Netherlands; (**O**) Belgium; (**P**) India; (**Q**) Italy; (**R**) Brazil; (**S**) Norway; (**T**) Ireland; (**U**) Croatia; (**V**) Indonesia.

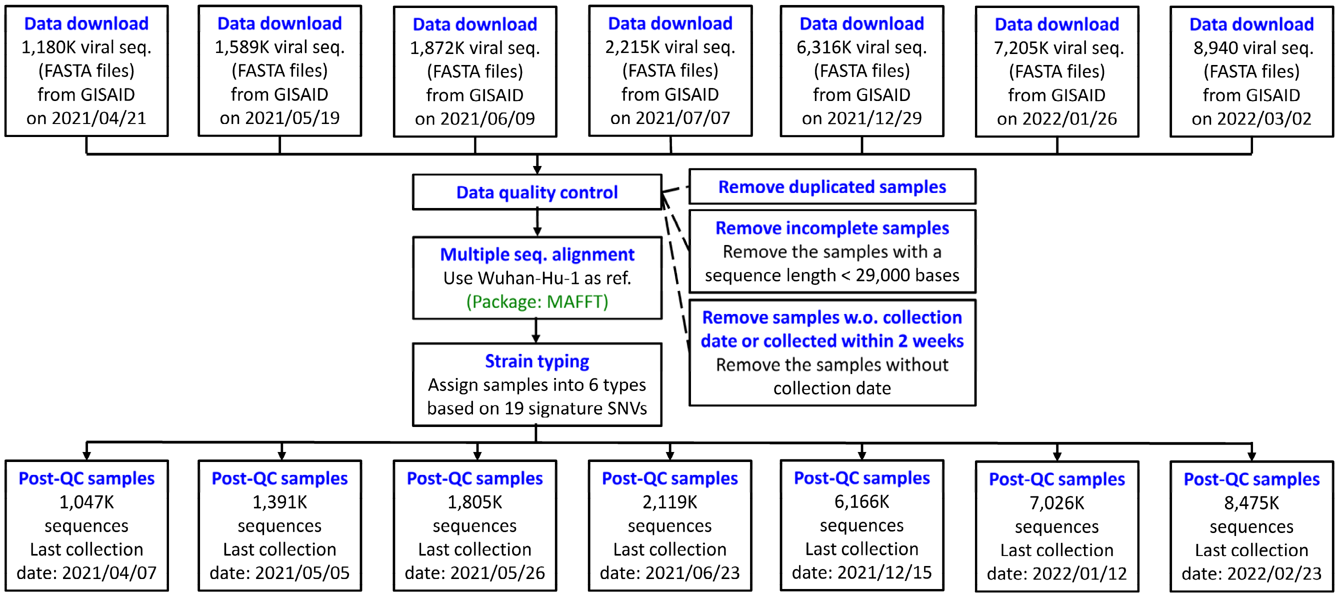


**Figure S15. Data download and preprocessing.** We downloaded 1,180K, 1,589K, 1,872K, 2,215K, 6316K, 7,205K, and 8,475K whole-genome sequences data from the Global Initiative on Sharing Avian Influenza Data (GISAID) database (https://www.gisaid.org/) on 7 Apr, 5 May, 26 May, 23 Jun, 15 Dec in year 2021 and 12 Jan and 23 Feb in year 2022 respectively. We discarded the duplicated samples, the samples with an aligned sequence of <29K bases, and the samples without sample recruitment date. The remained sequences contained 1,047K, 1,391K, 1,805K, 2,119K, 6,166K, 7,026K, and 8,475K genomes, respectively. Multiple sequence alignment was performed by using MAFFT v.7. The Wuhan-Hu-1 that the strain was originally isolated in China and had 29,903 nucleotides was employed as the reference genome. Our sequence analysis discarded two ends (5' leader and 3' terminal sequences) and focused on the SNV base positions from 266 to 29,674. Nucleotides different from the Wuhan-Hu-1 strain were assigned as a SNV. Deletions were also detected.

**Table S1. Proportions of intragenic and intergenic allelic association (n = 2,119K).** The number and proportion of SNVs having allelic association with another SNVs in the same gene (i.e., intragenic allelic association) and with other SNVs in the other gene (i.e., intergenic allelic association). The regions that they contain no SNVs with allelic association are marked by light green. The regions that they are top in the number of SNVs with allelic association are marked by red; more red, more SNVs with allelic association.

| | nsp1 | nsp2 | nsp3 | nsp4 | nsp5 | nsp6 | nsp7 | nsp8 | nsp9 | nsp10 | nsp11 | nsp12 | nsp13 |
|---|---|---|---|---|---|---|---|---|---|---|---|---|---|
| nsp1 | **0 (0%)** | 0 (0%) | 5 (1.82%) | 0 (0%) | 0 (0%) | 0 (0%) | 0 (NaN%) | 0 (0%) | 1 (2.17%) | 0 (0%) | 0 (NaN%) | 1 (0.72%) | 1 (1.18%) |
| nsp2 | 0 (0%) | **2 (1.67%)** | 12 (4.38%) | 3 (4.05%) | 4 (10.26%) | 1 (4%) | 0 (NaN%) | 2 (11.11%) | 3 (6.52%) | 0 (0%) | 0 (NaN%) | 7 (5.07%) | 4 (4.71%) |
| nsp3 | 5 (22.73%) | 12 (10%) | **17 (6.2%)** | 4 (5.41%) | 4 (10.26%) | 2 (8%) | 0 (NaN%) | 1 (5.56%) | 6 (13.04%) | 1 (25%) | 0 (NaN%) | 24 (17.39%) | 13 (15.29%) |
| nsp4 | 0 (0%) | 3 (2.5%) | 4 (1.46%) | **3 (4.05%)** | 1 (2.56%) | 4 (16%) | 0 (NaN%) | 1 (5.56%) | 2 (4.35%) | 0 (0%) | 0 (NaN%) | 1 (0.72%) | 5 (5.88%) |
| nsp5 | 0 (0%) | 4 (3.33%) | 4 (1.46%) | 1 (1.35%) | **0 (0%)** | 1 (4%) | 0 (NaN%) | 0 (0%) | 1 (2.17%) | 1 (25%) | 0 (NaN%) | 3 (2.17%) | 0 (0%) |
| nsp6 | 0 (0%) | 1 (0.83%) | 2 (0.73%) | 4 (5.41%) | 1 (2.56%) | **1 (4%)** | 0 (NaN%) | 0 (0%) | 1 (2.17%) | 0 (0%) | 0 (NaN%) | 1 (0.72%) | 0 (0%) |
| nsp7 | 0 (0%) | 0 (0%) | 0 (0%) | 0 (0%) | 0 (0%) | 0 (0%) | 0 (NaN%) | 0 (0%) | 0 (0%) | 0 (0%) | 0 (NaN%) | 0 (0%) | 0 (0%) |
| nsp8 | 0 (0%) | 2 (1.67%) | 1 (0.36%) | 1 (1.35%) | 0 (0%) | 0 (0%) | 0 (NaN%) | **0 (0%)** | 1 (2.17%) | 0 (0%) | 0 (NaN%) | 1 (0.72%) | 2 (2.35%) |
| nsp9 | 1 (4.55%) | 3 (2.5%) | 6 (2.19%) | 2 (2.7%) | 1 (2.56%) | 1 (4%) | 0 (NaN%) | 1 (5.56%) | **0 (0%)** | 0 (0%) | 0 (NaN%) | 1 (0.72%) | 2 (2.35%) |
| nsp10 | 0 (0%) | 0 (0%) | 1 (0.36%) | 0 (0%) | 1 (2.56%) | 0 (0%) | 0 (NaN%) | 0 (0%) | 0 (0%) | **0 (0%)** | 0 (NaN%) | 1 (0.72%) | 0 (0%) |
| nsp11 | 0 (0%) | 0 (0%) | 0 (0%) | 0 (0%) | 0 (0%) | 0 (0%) | 0 (NaN%) | 0 (0%) | 0 (0%) | 0 (0%) | 0 (NaN%) | 0 (0%) | 0 (0%) |
| nsp12 | 1 (4.55%) | 7 (5.83%) | 24 (8.76%) | 1 (1.35%) | 3 (7.69%) | 1 (4%) | 0 (NaN%) | 1 (5.56%) | 1 (2.17%) | 1 (25%) | 0 (NaN%) | **3 (2.17%)** | 6 (7.06%) |
| nsp13 | 1 (4.55%) | 4 (3.33%) | 13 (4.74%) | 5 (6.76%) | 0 (0%) | 0 (0%) | 0 (NaN%) | 2 (11.11%) | 2 (4.35%) | 0 (0%) | 0 (NaN%) | 6 (4.35%) | **2 (2.35%)** |
| nsp14 | 0 (0%) | 2 (1.67%) | 1 (0.36%) | 0 (0%) | 1 (2.56%) | 0 (0%) | 0 (NaN%) | 0 (0%) | 0 (0%) | 0 (0%) | 0 (NaN%) | 2 (1.45%) | 0 (0%) |
| nsp15 | 0 (0%) | 1 (0.83%) | 4 (1.46%) | 3 (4.05%) | 2 (5.13%) | 2 (8%) | 0 (NaN%) | 0 (0%) | 2 (4.35%) | 0 (0%) | 0 (NaN%) | 0 (0%) | 1 (1.18%) |
| nsp16 | 0 (0%) | 2 (1.67%) | 2 (0.73%) | 2 (2.7%) | 2 (5.13%) | 1 (4%) | 0 (NaN%) | 1 (5.56%) | 1 (2.17%) | 0 (0%) | 0 (NaN%) | 2 (1.45%) | 0 (0%) |
| S | 9 (40.91%) | 29 (24.17%) | 83 (30.29%) | 19 (25.68%) | 7 (17.95%) | 4 (16%) | 0 (NaN%) | 5 (27.78%) | 15 (32.61%) | 0 (0%) | 0 (NaN%) | 31 (22.46%) | 25 (29.41%) |
| ORF3a | 1 (4.55%) | 4 (3.33%) | 14 (5.11%) | 11 (14.86%) | 3 (7.69%) | 3 (12%) | 0 (NaN%) | 0 (0%) | 1 (2.17%) | 0 (0%) | 0 (NaN%) | 5 (3.62%) | 4 (4.71%) |
| E | 0 (0%) | 1 (0.83%) | 1 (0.36%) | 0 (0%) | 1 (2.56%) | 0 (0%) | 0 (NaN%) | 0 (0%) | 0 (0%) | 0 (0%) | 0 (NaN%) | 0 (0%) | 0 (0%) |
| M | 0 (0%) | 2 (1.67%) | 0 (0%) | 2 (2.7%) | 0 (0%) | 0 (0%) | 0 (NaN%) | 1 (5.56%) | 1 (2.17%) | 0 (0%) | 0 (NaN%) | 1 (0.72%) | 2 (2.35%) |
| ORF6 | 0 (0%) | 0 (0%) | 0 (0%) | 0 (0%) | 0 (0%) | 0 (0%) | 0 (NaN%) | 0 (0%) | 0 (0%) | 0 (0%) | 0 (NaN%) | 0 (0%) | 0 (0%) |
| ORF7a | 0 (0%) | 0 (0%) | 2 (0.73%) | 2 (2.7%) | 0 (0%) | 0 (0%) | 0 (NaN%) | 0 (0%) | 0 (0%) | 0 (0%) | 0 (NaN%) | 0 (0%) | 1 (1.18%) |
| ORF7b | 0 (0%) | 0 (0%) | 0 (0%) | 0 (0%) | 0 (0%) | 0 (0%) | 0 (NaN%) | 0 (0%) | 0 (0%) | 0 (0%) | 0 (NaN%) | 0 (0%) | 0 (0%) |
| ORF8 | 1 (4.55%) | 18 (15%) | 26 (9.49%) | 6 (8.11%) | 3 (7.69%) | 2 (8%) | 0 (NaN%) | 1 (5.56%) | 2 (4.35%) | 0 (0%) | 0 (NaN%) | 17 (12.32%) | 6 (7.06%) |
| N | 3 (13.64%) | 23 (19.17%) | 52 (18.98%) | 5 (6.76%) | 5 (12.82%) | 2 (8%) | 0 (NaN%) | 2 (11.11%) | 6 (13.04%) | 1 (25%) | 0 (NaN%) | 31 (22.46%) | 11 (12.94%) |
| ORF10 | 0 (0%) | 0 (0%) | 0 (0%) | 0 (0%) | 0 (0%) | 0 (0%) | 0 (NaN%) | 0 (0%) | 0 (0%) | 0 (0%) | 0 (NaN%) | 0 (0%) | 0 (0%) |
| Sums | 22 | 120 | 274 | 74 | 39 | 25 | 0 | 18 | 46 | 4 | 0 | 138 | 85 |

(cont'd)

| | nsp14 | nsp15 | nsp16 | S | ORF3a | E | M | ORF6 | ORF7a | ORF7b | ORF8 | N | ORF10 |
|---|---|---|---|---|---|---|---|---|---|---|---|---|---|
| nsp1 | 0 (0%) | 0 (0%) | 0 (0%) | 9 (1.84%) | 1 (0.8%) | 0 (0%) | 0 (0%) | 0 (NaN%) | 0 (0%) | 0 (NaN%) | 1 (0.5%) | 3 (0.95%) | 0 (NaN%) |
| nsp2 | 2 (12.5%) | 1 (3.12%) | 2 (7.14%) | 29 (5.94%) | 4 (3.2%) | 1 (9.09%) | 2 (12.5%) | 0 (NaN%) | 0 (0%) | 0 (NaN%) | 18 (9.05%) | 23 (7.28%) | 0 (NaN%) |
| nsp3 | 1 (6.25%) | 4 (12.5%) | 2 (7.14%) | 83 (17.01%) | 14 (11.2%) | 1 (9.09%) | 0 (0%) | 0 (NaN%) | 2 (14.29%) | 0 (NaN%) | 26 (13.07%) | 52 (16.46%) | 0 (NaN%) |
| nsp4 | 0 (0%) | 3 (9.38%) | 2 (7.14%) | 19 (3.89%) | 11 (8.8%) | 0 (0%) | 2 (12.5%) | 0 (NaN%) | 2 (14.29%) | 0 (NaN%) | 6 (3.02%) | 5 (1.58%) | 0 (NaN%) |
| nsp5 | 1 (6.25%) | 2 (6.25%) | 2 (7.14%) | 7 (1.43%) | 3 (2.4%) | 1 (9.09%) | 0 (0%) | 0 (NaN%) | 0 (0%) | 0 (NaN%) | 3 (1.51%) | 5 (1.58%) | 0 (NaN%) |
| nsp6 | 0 (0%) | 2 (6.25%) | 1 (3.57%) | 4 (0.82%) | 3 (2.4%) | 0 (0%) | 0 (0%) | 0 (NaN%) | 0 (0%) | 0 (NaN%) | 2 (1.01%) | 2 (0.63%) | 0 (NaN%) |
| nsp7 | 0 (0%) | 0 (0%) | 0 (0%) | 0 (0%) | 0 (0%) | 0 (0%) | 0 (0%) | 0 (NaN%) | 0 (0%) | 0 (NaN%) | 0 (0%) | 0 (0%) | 0 (NaN%) |
| nsp8 | 0 (0%) | 0 (0%) | 1 (3.57%) | 5 (1.02%) | 0 (0%) | 0 (0%) | 1 (6.25%) | 0 (NaN%) | 0 (0%) | 0 (NaN%) | 1 (0.5%) | 2 (0.63%) | 0 (NaN%) |
| nsp9 | 0 (0%) | 2 (6.25%) | 1 (3.57%) | 15 (3.07%) | 1 (0.8%) | 0 (0%) | 1 (6.25%) | 0 (NaN%) | 0 (0%) | 0 (NaN%) | 2 (1.01%) | 6 (1.9%) | 0 (NaN%) |
| nsp10 | 0 (0%) | 0 (0%) | 0 (0%) | 0 (0%) | 0 (0%) | 0 (0%) | 0 (0%) | 0 (NaN%) | 0 (0%) | 0 (NaN%) | 0 (0%) | 1 (0.32%) | 0 (NaN%) |
| nsp11 | 0 (0%) | 0 (0%) | 0 (0%) | 0 (0%) | 0 (0%) | 0 (0%) | 0 (0%) | 0 (NaN%) | 0 (0%) | 0 (NaN%) | 0 (0%) | 0 (0%) | 0 (NaN%) |
| nsp12 | 2 (12.5%) | 0 (0%) | 2 (7.14%) | 31 (6.35%) | 5 (4%) | 0 (0%) | 1 (6.25%) | 0 (NaN%) | 0 (0%) | 0 (NaN%) | 17 (8.54%) | 31 (9.81%) | 0 (NaN%) |
| nsp13 | 0 (0%) | 1 (3.12%) | 0 (0%) | 25 (5.12%) | 4 (3.2%) | 0 (0%) | 2 (12.5%) | 0 (NaN%) | 1 (7.14%) | 0 (NaN%) | 6 (3.02%) | 11 (3.48%) | 0 (NaN%) |
| nsp14 | **0 (0%)** | 0 (0%) | 1 (3.57%) | 2 (0.41%) | 3 (2.4%) | 0 (0%) | 0 (0%) | 0 (NaN%) | 0 (0%) | 0 (NaN%) | 1 (0.5%) | 3 (0.95%) | 0 (NaN%) |
| nsp15 | 0 (0%) | **1 (3.12%)** | 2 (7.14%) | 8 (1.64%) | 2 (1.6%) | 0 (0%) | 0 (0%) | 0 (NaN%) | 1 (7.14%) | 0 (NaN%) | 1 (0.5%) | 2 (0.63%) | 0 (NaN%) |
| nsp16 | 1 (6.25%) | 2 (6.25%) | **0 (0%)** | 3 (0.61%) | 3 (2.4%) | 0 (0%) | 0 (0%) | 0 (NaN%) | 1 (7.14%) | 0 (NaN%) | 1 (0.5%) | 4 (1.27%) | 0 (NaN%) |
| S | 2 (12.5%) | 8 (25%) | 3 (10.71%) | **72 (14.75%)** | 33 (26.4%) | 5 (45.45%) | 4 (25%) | 0 (NaN%) | 2 (14.29%) | 0 (NaN%) | 49 (24.62%) | 83 (26.27%) | 0 (NaN%) |
| ORF3a | 3 (18.75%) | 2 (6.25%) | 3 (10.71%) | 33 (6.76%) | **4 (3.2%)** | 1 (9.09%) | 0 (0%) | 0 (NaN%) | 3 (21.43%) | 0 (NaN%) | 13 (6.53%) | 17 (5.38%) | 0 (NaN%) |
| E | 0 (0%) | 0 (0%) | 0 (0%) | 5 (1.02%) | 1 (0.8%) | **0 (0%)** | 0 (0%) | 0 (NaN%) | 0 (0%) | 0 (NaN%) | 1 (0.5%) | 1 (0.32%) | 0 (NaN%) |
| M | 0 (0%) | 0 (0%) | 0 (0%) | 4 (0.82%) | 0 (0%) | 0 (0%) | **0 (0%)** | 0 (NaN%) | 0 (0%) | 0 (NaN%) | 1 (0.5%) | 2 (0.63%) | 0 (NaN%) |
| ORF6 | 0 (0%) | 0 (0%) | 0 (0%) | 0 (0%) | 0 (0%) | 0 (0%) | 0 (0%) | 0 (NaN%) | 0 (0%) | 0 (NaN%) | 0 (0%) | 0 (0%) | 0 (NaN%) |
| ORF7a | 0 (0%) | 1 (3.12%) | 1 (3.57%) | 2 (0.41%) | 3 (2.4%) | 0 (0%) | 0 (0%) | 0 (NaN%) | **0 (0%)** | 0 (NaN%) | 1 (0.5%) | 1 (0.32%) | 0 (NaN%) |
| ORF7b | 0 (0%) | 0 (0%) | 0 (0%) | 0 (0%) | 0 (0%) | 0 (0%) | 0 (0%) | 0 (NaN%) | 0 (0%) | 0 (NaN%) | 0 (0%) | 0 (0%) | 0 (NaN%) |
| ORF8 | 1 (6.25%) | 1 (3.12%) | 1 (3.57%) | 49 (10.04%) | 13 (10.4%) | 1 (9.09%) | 1 (6.25%) | 0 (NaN%) | 1 (7.14%) | 0 (NaN%) | **14 (7.04%)** | 35 (11.08%) | 0 (NaN%) |
| N | 3 (18.75%) | 2 (6.25%) | 4 (14.29%) | 83 (17.01%) | 17 (13.6%) | 1 (9.09%) | 2 (12.5%) | 0 (NaN%) | 1 (7.14%) | 0 (NaN%) | 35 (17.59%) | **27 (8.54%)** | 0 (NaN%) |
| ORF10 | 0 (0%) | 0 (0%) | 0 (0%) | 0 (0%) | 0 (0%) | 0 (0%) | 0 (0%) | 0 (NaN%) | 0 (0%) | 0 (NaN%) | 0 (0%) | 0 (0%) | 0 (NaN%) |
| Sums | 16 | 32 | 28 | 488 | 125 | 11 | 16 | 0 | 14 | 0 | 199 | 316 | 0 |

**Table S2. Distribution of the viral strains and SNVs represented by correlated SNV sets (CSSs) (n = 2,119K).** Distribution of the viral strains and SNVs represented by CSSs. The analysis was conducted based on the genome sequences of more than 2 million of SARS-CoV-2 viral strains collected before 23 Jun, 2021 (n = 2,215K) from the GISAID (https://www.gisaid.org/). After data quality control, 2,119,724 strains remained. The majority of the viral strains can be characterized by a single CSS or a few numbers of the 1,057 CSSs consisting of 1,384 distinct signature SNVs (from 1,432 signature SNVs that some SNVs have various alternative alleles). We find that 1,053 of 1,057 CSSs can characterize >99.9% of the current dominant strain type (Type VI). Type VI became the dominant strain (2,000,622 strains and 94.38%) since March, 2020. Among 2,000,622 Type VI strains, 98.79% of the strains can be further characterized by 171 “primary” CSSs which are constructed by 557 SNVs. A primary CSS defines the subtype within which at least one strain does not belong to other subtypes. The remaining 882 “secondary” CSSs, by which the subtype strains defined already belong to one or multiple subtypes defined by primary CSSs, can be explained by the 171 primary CSSs. The primary CSSs provide a backbone for subtyping, whereas the secondary CSSs allow more detailed characterization. The proportion explained by CSSs is further increased to 99.95% if additional 42 SNVs with a variation frequency of >0.01 are accounted. The CSSs can cover the well-recognized variants of interest such as B.1.1.7 (alias as variant 501Y.v1 or Alpha), B.1.351 (alias as variant 501Y.v2 or Beta), P.1 (alias as variant 501Y.v3 or Gamma), and B.1.617.2 (alias as variant Delta) and exhibit the subtypes and heterogeneity of these variants.

| n = 2,119,724 | | | **CSS** | | **non-CSS** | | |
|---|---|---|---|---|---|---|---|
| (A) 14 signatures | | | (B) | (C) | (D) | (E) | |
| | | | haplotype | single SNV | SNV with MAF > 0.01 | SNVs with MAF ≤ 0.01 | (B)+(C)+(D) |
| | | | 22 CSSs, 88 SNVs | | 33 SNVs | 4435 SNVs | |
| **Type I** | 11,020 | 0.5199% | 141 | 6,661 | 3,755 | 463 | 10,557 |
| | | | 0.0067% | 0.3142% | 0.1771% | 0.0218% | 0.4980% |
| | | | 2 CSSs, 17 SNVs | | 36 SNVs | 1334 SNVs | |
| **Type II** | 9,835 | 0.4640% | 1052 | 7,577 | 1,190 | 16 | 9,819 |
| | | | 0.0496% | 0.3575% | 0.0561% | 0.0008% | 0.4632% |
| | | | - | | 24 SNVs | 3906 SNVs | |
| **Type III** | 6,259 | 0.2953% | 0 | 1,876 | 4,339 | 44 | 6,215 |
| | | | 0.0000% | 0.0885% | 0.2047% | 0.0021% | 0.2932% |
| | | | - | | 36 SNVs | 1574 SNVs | |
| **Type IV** | 2,891 | 0.1364% | 0 | 757 | 1,751 | 383 | 2,508 |
| | | | 0.0000% | 0.0357% | 0.0826% | 0.0181% | 0.1183% |
| | | | - | | 67 SNVs | - | |
| **Type V** | 637 | 0.0301% | 0 | 601 | 36 | 0 | 637 |
| | | | 0.0000% | 0.0284% | 0.0017% | 0.0000% | 0.0301% |
| | | | 171 CSSs, 577 SNVs | | 42 SNVs | 17904 SNVs | |
| **Type VI** | 2,000,622 | 94.3812% | 1521736 | 454,708 | 23,141 | 1037 | 1,999,585 |
| | | | 71.7893% | 21.4513% | 1.0917% | 0.0489% | 94.3323% |
| | | | 162 CSSs, 508 SNVs | | 46 SNVs | - | |
| **Others** | 88,460 | 4.1732% | 47423 | 40,704 | 333 | 0 | 88,460 |
| | | | 2.2372% | 1.9202% | 0.0157% | 0.0000% | 4.1732% |
| | | | 1,570,352 | 512,884 | 34,545 | | 2,117,781 |
| | | | 74.0829% | 24.1958% | 1.6297% | | 99.9083% |

***Supplemental Text 1: Allelic association is a characteristic of rising variants***

To develop a method for analyzing the big data of genome sequence, we first examined the genome sequences of 1 million of SARS-CoV-2 viral strains collected before April 7, 2021 (n = 1,180,744) from GISAID. After data quality control, 1,047,410 strains remained. We found that strains with high occurrence frequency almost always contain SNVs with strong allelic associations, and that the frequency of strains started to increase mostly after the allelic association of some pairs of SNVs increases. The results are further confirmed in the updated data of 2,119K (**Fig. S1**; data collection before June 23, 2021). For cases of low allelic associations, the strain frequency is rarely increased to a high level. The observed pattern reveals that allelic association of SNVs is a hallmark of increasing strain occurrence frequency. This characteristic allows us to use allelic association as a way to reduce the dimension of the big data and subtype the variants.

We found that ORF8, ORF7a, N (the nucleocapsid protein), S (the spike protein) especially for the receptor binding domain (RBD), and nsp9 (the non-structural protein 9) are the top genomic regions that have a high proportion of SNVs with allelic association (**Fig. S2A**). Moreover, N, S, RBD in S, ORF8, and nsp3 have the highest ratio of non-synonymous SNVs vs. synonymous SNVs with allelic association (**Fig. S2B**). We further examined the intragenic and intergenic allelic associations. S, N, and nsp3 exhibit the most abundant SNVs that they have allelic association with the SNVs in other proteins (**Fig. S3**). Among the proteins involved in intergenic allelic association, S-N association are the most frequent (**Figs. S3** and **S4**). S is also the region that exhibits the most abundant SNVs that they have within-gene allelic association (**Fig. S3**). Detailed statistics are provided (**Table S1**).

These results reveal that the SNVs in S pivotally involve in intra- and inter-gene allelic associations. Non-spike protein SNVs may enhance viral transmissibility through a direct genetic epistasis with spike SNVs (i.e., genetic buffering) or an indirect effect of genetic hitchhiking mainly contributed by spike SNVs. Therefore, the use of allelic association as a means to achieve dimension reduction and subtyping is biologically meaningful.

***Supplemental Text 2: Using correlated SNV sets (CSSs) to subtype existing variants***

To understand the evolutionary relationship among the strains, the conventional strain-based phylogenetic tree analysis requires intensive computation and becomes intractable for a data set with multi-million genome sequences. CSSs provide a computationally efficient and epidemiologically relevant dimension reduction for the viral genome dataset. Therefore, we used CSSs to construct a phylogenetic tree for the 2 million genomes, using the maximum parsimony method (Edwards & Cavallisforza, 1963). The dendrogram analysis reveals the important variants such as the Variants of Concern and Interest, their CSSs, and the differential transmission patterns and evolution relationship in the subtypes (**Fig. S8**).

The dendrogram was arranged based on the number of taxa in a descending order with the largest number of terminal taxa on top of the dendrogram. The majority of strains in this top group are Delta (aka B.1.617.2). This group has a deep-rooted branch topology, implying that this CSSs group (or subtype) has undergone a series of mutations (refer to the phylogenetic dendrogram in **Fig. S8**). Delta strains are separated into two major CSSs subgroups. The two CSSs subgroups carry similar SNVs for the defining SNVs for Delta, according to the Pango nomenclature (Rochman et al., 2021) from the metadata in GISAID (refer to the variation matrix in **Fig. S8**). However, compared to the first CSSs subgroup, the second CSSs subgroup has many more additional SNVs with allelic association in ORF1ab (**Fig. S8**).

We also examined the dendrogram for Alpha (aka B.1.1.7 or 501Y.v1). This group has a deep-rooted branch topology, implying that this CSSs group has undergone a series of mutations (**Fig. S8**). Two major CSSs subgroups were found. One is closer to the root and the other is to the leaf. The CSSs subgroup closer to the root (i.e., the recent common ancestor of Alpha) indicates that most of the CSSs are primary CSSs and have less SNVs with allelic association compared to the CSSs subgroup closer to the leaf (refer to the variation matrix in **Fig. S8**). The subgroup closer to the leaf has a wide dendrogram illustrates that this subgroup has more CSSs, implying that the CSSs have spread to more countries, and/or more variations are accumulated. Other variants of interest such as Beta (aka variant B.1.351 or 501Y.v2), Gamma (aka variant P.1 or 501Y.v3), etc. are also represented in the tree dendrogram with many subtypes and heterogeneity within these variants.
